# Cell-type-specific 3D-genome organization and transcription regulation in the brain

**DOI:** 10.1101/2023.12.04.570024

**Authors:** Shiwei Liu, Pu Zheng, Cosmos Yuqi Wang, Bojing Blair Jia, Nathan R. Zemke, Bing Ren, Xiaowei Zhuang

## Abstract

3D organization of the genome plays a critical role in regulating gene expression. However, it remains unclear how chromatin organization differs among different cell types in the brain. Here we used genome-scale DNA and RNA imaging to investigate 3D-genome organization in transcriptionally distinct cell types in the primary motor cortex of the mouse brain. We uncovered a wide spectrum of differences in the nuclear architecture and 3D-genome organization among different cell types, ranging from the physical size of the cell nucleus to the active-inactive chromatin compartmentalization and radial positioning of chromatin loci within the nucleus. These cell-type-dependent variations in nuclear architecture and chromatin organization exhibited strong correlation with both total transcriptional activity of the cell and transcriptional regulation of cell-type-specific marker genes. Moreover, we found that the methylated-DNA-binding protein MeCP2 regulates transcription in a divergent manner, depending on the nuclear radial positions of chromatin loci, through modulating active-inactive chromatin compartmentalization.

## Main Text

The eukaryotic nucleus is the central hub for essential genomic functions, ranging from transcription and gene-expression regulation to the replication of DNA. The morphology and molecular architecture of the cell nucleus are among the most distinct indicators of cell differentiation, aging, and disease progression (*1, 2*). Decades of imaging and biochemical studies have provided rich insights into the nuclear architecture and chromatin organization across scales (*3–9*). Recent development of high-throughput sequencing- and imaging-based assays for 3D-genome mapping has substantially advanced our understanding of the chromatin structure in the interphase nucleus with a genome-wide view (*3–5, 8–15*). Studies using these methods have revealed prominent chromatin structures such as chromatin compartments (*16*), topologically associating domains (TADs) (*17–19*), and chromatin loops (*20*). For example, both sequencing- and imaging-based methods have shown that chromatin in the interphase nucleus are segregated into compartments enriched for active and inactive chromatin marks (termed A and B compartments), with compartment-A and compartment-B chromatin preferentially positioned in the interior and periphery of the nucleus, respectively (*16, 21–24*).

Recent studies comparing embryonic stem cells and a few differentiated cell types in vitro, as well as comparisons between major cell types in the brain, have revealed changes in A/B compartment arrangements from cell type to cell type, which are correlated with gene expression changes (*25–31*). However, these effects tend to be subtle and exactly how the A/B compartmentalization affects the transcription activity of chromatin remains unclear. In addition, TADs and loop domains have been observed in different cell types, and the boundaries of these domains vary between cell types (*25–27, 29, 31–33*). However, genetic manipulations that abolish TADs only have a small effect on transcription activities (*34–36*). Therefore, what properties of the 3D chromatin organization are related to transcription and what molecular mechanisms underlie these connections remain incompletely understood. Moreover, most of the cell-type-dependence studies of 3D-genome organization have investigated only a small number of cells or cell types. Hence what differences in 3D chromatin organization are present between different cell types in complex tissues and how 3D chromatin organization influences or is influenced by transcription across different cell types remain poorly understood.

3D organization of the genome is regulated by a variety of protein factors that establish, maintain, or modulate chromatin structures (*37*). For example, cohesin and CTCF are known to regulate chromatin organization, in particular chromatin domains and loops, and the underlying mechanisms have been extensively characterized (*38*). However, for most transcription regulators, the mechanisms underlying how they control or modulate chromatin organization are less clear. Many of these factors, such as Med1, Brd4, and MeCP2, bind to gene regulatory elements, including histone marks and methylated DNA, and are suggested to drive 3D-genome organization through phase separation (*6, 8*). Among these proteins, MeCP2, the causal gene of the Rett syndrome – a neurological disorder (*39*), binds to methylated DNA (*40*), forms liquid-liquid phase separation (*41, 42*), and regulates gene transcription (*43, 44*). Although *Mecp2* deletion or overexpression has been reported to change the size of the nucleus and gross nuclear architecture, as reflected by DAPI and histone mark stains (*41, 42, 45, 46*), how MeCP2 affects higher-order chromatin structures and how the chromatin-organizing function of MeCP2 regulates transcription mechanistically remain unclear.

In this work, we studied the 3D-genome organization and the effect of MeCP2 on chromatin organization across different cell types in the brain using multiplexed error-robust fluorescence in situ hybridization (MERFISH), a genome-scale imaging method (*47*). We have previously demonstrated *in situ* cell-type identification in the brain using RNA-MERFISH (*48–50*) and the determination of 3D-genome organization in cultured cells using DNA-MERFISH (*24*). Here, we extended DNA-MERFISH to intact tissue samples and combined it with RNA-MERFISH to probe cell-type-specific 3D-genome organization in ∼20 major cell types in the primary motor cortex (MOp) of the mouse brain.

Using this approach, we observed multiple levels of differences in the nuclear architecture and chromatin organization between different cell types that were related to transcriptional regulation. First, we showed that the nuclear sizes and chromosome territory sizes varied drastically across different cell types, both of which were strongly correlated with the total transcriptional activity of the cells. Second, we observed cell-type-dependent changes in higher-order chromatin structures. In particular, chromatin exhibited increasingly more pronounced A/B compartments and less pronounced megadomain structures as the overall transcriptional activity increased from non-neuronal cells to inhibitory neurons and then to excitatory neurons. Third, transcriptional activity of differentially expressed genes, as well as the activity of cell-type-specific super-enhancer activities, were correlated with the local enrichment of compartment-A chromatin loci at these genes or super-enhancers across different cell types. Fourth, we observed different nuclear radial positioning of genomic loci in different cell types. In non-neuronal cells, transcriptional activity of genomic loci was strongly correlated with their radial positions in the nucleus, with active and inactive chromatin enriched in the nuclear interior and periphery, respectively. In contrast, active and inactive chromatin in neurons adopted a more intermixed radial positioning in the nucleus. Finally, we found that MeCP2 regulated nuclear radial positioning, local A/B compartmentalization, and transcriptional activity of chromatin loci in a radial-position-dependent and cell-type-specific manner. Notably, upon *Mecp2* deletion, genes located in the nuclear periphery showed increased transcriptional activity whereas genes located near nuclear centers showed decreased transcriptional activity, and these divergent changes can be explained by a common mechanism – loss of MeCP2 weakened A/B-compartment chromatin segregation. Overall, our data provided rich insights into how 3D chromatin organization is connected to transcriptional activities in a cell-type-specific manner in the brain.

### Integrated RNA- and DNA-MERFISH for cell-type-specific chromatin organization mapping

We chose the primary motor cortex (MOp) as a model system to study the cell-type-specific chromatin organization and its relationship to transcriptional regulation. A comprehensive cell-type atlas of the MOp has been generated by the BRAIN Initiative Cell Census Network (*51*). Specifically, single-cell/nucleus RNA sequencing (sc/snRNA-seq) and epigenomic sequencing studies have extensively characterized cell-type taxonomy as well as the transcriptional regulatory landscape across all identified cell types in the MOp, including the transcriptomic, chromatin accessibility, and DNA-methylation profiles of each cell type (*52–54*). In parallel, we have generated a spatially resolved cell atlas of the MOp by RNA-MERFISH, reporting a comprehensive list of cell types and their spatial organization in this brain region (*50*). Here, we sought to develop a platform that combines RNA-MERFISH for resolving transcriptionally defined cell types and DNA-MERFISH for characterizing chromatin structures in single cells to investigate the relationship between cell-type-specific chromatin organization and gene-expression regulation.

We introduced several modifications to our previous MERFISH protocols to facilitate the combination of RNA- and DNA-MERFISH. To preserve 3D chromatin structures in their native environment, we omitted the gel-embedding-based tissue clearing procedures (Materials and Methods), which were previously used in RNA-MERFISH characterizations of MOp cell types (*50*). To compensate for the decrease in signal-to-noise ratio resulted from omitting tissue clearing, we used an adaptor-probe approach to boost signal, which utilizes an adaptor probe to link dye-labeled readout probes to encoding probes, the latter in turn are bound to target RNA or DNA (*24*). This approach allows more dye-labeled readout probes to be linked to each encoding probe and hence increases signals from individual mRNA molecules and genomic loci.

Additionally, we employed photobleaching to reduce autofluorescence of the tissue slices (*49*). With these modifications, we performed RNA-MERFISH measurements on coronal mouse brain sections encompassing the MOp (fig. S1). Subsequently, we conducted DNA-MERFISH on the same tissue sections and registered the DNA and RNA images to obtain transcriptional profiles and chromatin 3D structures in the same cells (fig. S1).

To systematically study the cell-type-dependent chromatin structures, we designed encoding probe libraries to target cellular RNAs and chromatin in the following manner. For RNA-MERFISH, we utilized the same MOp encoding-probe library as in our previous work, targeting 242 marker genes for cell-type identification (*50*). For DNA-MERFISH, we designed three different encoding-probe libraries respectively targeting three groups of genomic loci (1981 loci in total): (1) 988 loci evenly distributed across the mouse genome with ∼2.5Mb spacing; (2) 28 loci centered around transcription start sites (TSSs) for representative marker genes for the major cell types in the MOp; (3) 965 loci centered around candidate super-enhancers, genomic regions that comprise multiple putative enhancers and have been implicated in cell-type and gene-expression specification (*55, 56*). We selected cell-type specific super-enhancers using the published single-nucleus Assay for Transposase-Accessible Chromatin using Sequencing (snATAC-seq) dataset (*52*) (Materials and Methods).

Using this design, we profiled transcription of the 242 genes in 46,340 cells and identified 21 cell types at the subclass level in the MOp and adjacent areas by *de novo* cell clustering (Fig. 1A). These include eight subclasses of excitatory neurons (layer 2/3 (L2/3) intratelencephalic (IT), L4/5 IT, L5 IT, and L6 IT, L5 extratelencephalic (ET), L5/6 near-projection (NP), L6 cortical-thalamic projection (CT), and L6b neurons), five subclasses of inhibitory neurons annotated by their canonical marker genes (*Pvalb*, *Sst*, *Lamp5*, *Vip* and *Sncg*), and eight subclasses of non-neuronal cells (astrocytes, oligodendrocytes, oligodendrocyte progenitor cells (OPC), microglia, endothelial cells, pericytes, vascular leptomeningeal cells (VLMCs), and smooth muscle cells (SMCs)) (Fig. 1, A and B). The overall expression profiles of MOp cells obtained using our modified RNA-MERFISH protocol agreed well with both our previous results from tissue-cleared RNA-MERFISH (*50*) (fig. S2A) and bulk RNA-seq data (fig. S2B). Our data also showed high reproducibility between experimental replicates (fig. S2C). Hence, the cell types identified using this modified RNA-MERFISH protocol showed a one-to-one correspondence with the cell types determined from our previously published RNA-MERFISH dataset (*50*), except for the low abundance perivascular macrophages (PVMs) which was not identified here presumably due to fewer cells measured here (fig. S2, D and E).

**Fig. 1.**
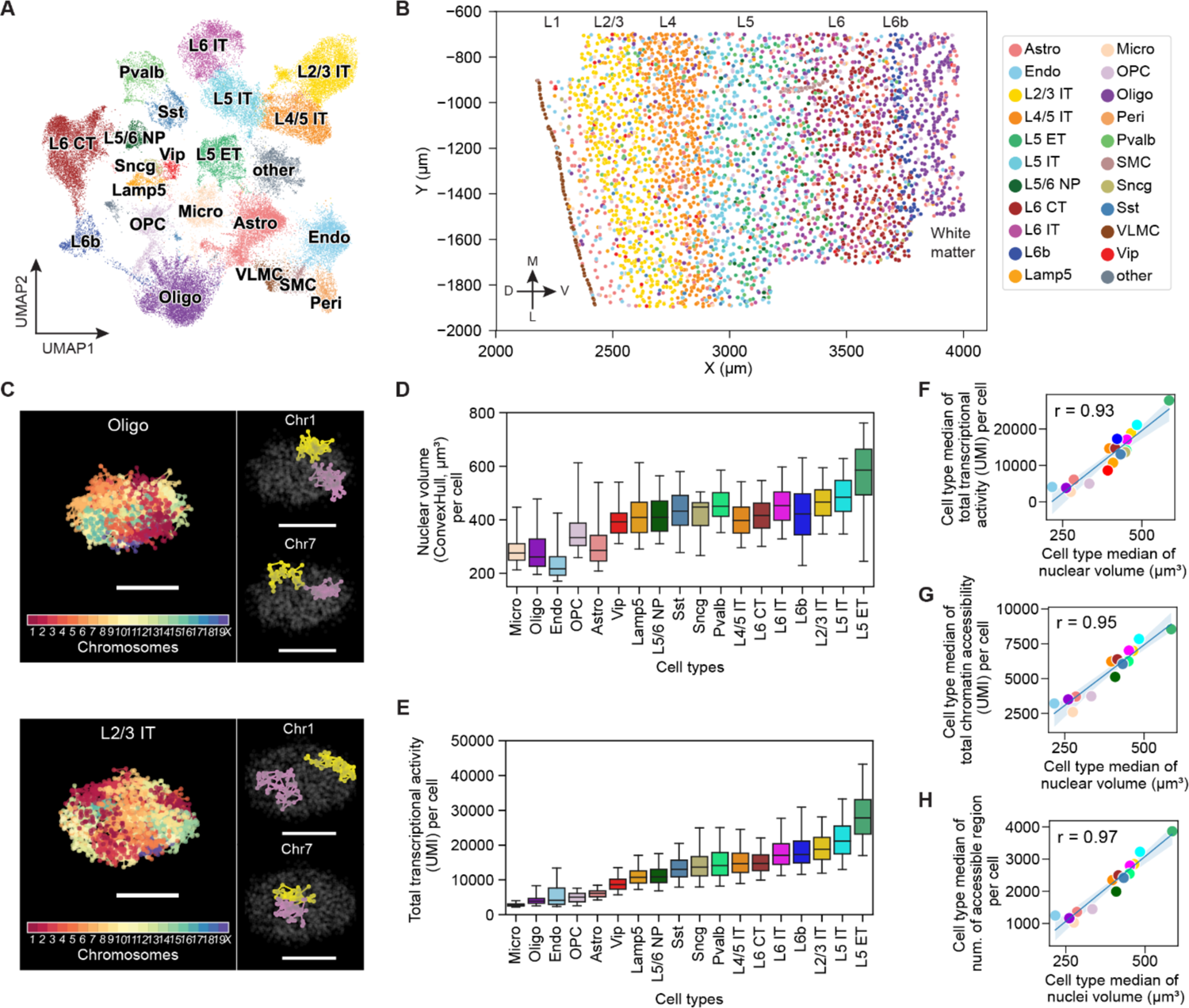
Integrated 3D-genome and transcriptome imaging revealed cell-type-dependent variations of and correlation between cell-nucleus size and transcriptional activity. (**A**) Uniform manifold approximation and projection (UMAP) representation of transcriptionally distinct cell types in the MOp determined by RNA-MERFISH. Cells are colored by their cell-type identities, and the same color code is used in all panels in this figure. Micro: microglia; Oligo: oligodendrocytes; OPC: Oligodendrocyte progenitor cells; Astro: astrocytes; Endo: endothelial cells; Peri: Pericytes; Vip: *vip*-positive inhibitory neurons; Lamp5: *lamp5*-positive inhibitory neurons; Sst: *sst*-positive inhibitory neurons; Sncg: *sncg*-positive inhibitory neurons; Pvalb: *pvalb*-positive inhibitory neurons; L2/3 IT, L4/5 IT, L5 IT, and L6 IT: L2/3 IT, L4/5 IT, L5 IT and L6 IT excitatory neurons; L5 ET: L5 ET excitatory neurons; L5/6 NP: L5/6 NP excitatory neurons; L6 CT: L6 CT excitatory neurons; L6b: L6b excitatory neurons. (**B**) Spatial map of transcriptionally distinct cell types in a coronal slice of the MOp. Different cortical layers are indicated on top (L1-L6b). D, dorsal; L, lateral; M, medial; V, ventral. (**C**) 3D renderings of chromatin organization in an oligodendrocyte (top) and L2/3 IT neuron (bottom) determined by DNA-MERFISH. Left, all chromosome loci colored according to their chromosome identities; Right, Chr1 and Chr7, whose two homologs are distinctively colored in yellow and purple. Scale bar: 5 μm. (**D**) Boxplot for the distribution of nuclear volume across individual cells in each cell type. Nuclear volume per cell was calculated by the convex hull volume of all chromosome loci. (**E**) Boxplot for the distribution of total transcriptional activity across individual cells in each cell type. Total transcriptional activity per cell was calculated using the sum of unique molecular identifiers (UMIs) from snRNA-seq data of the MOp (*52*). Cell types are ordered by their rank in the cell-type median of total transcriptional activity per cell. (**F**) Scatterplot of the cell-type medians of total transcriptional activity versus nuclear volume per cell. (**G**) Scatterplot of the cell-type medians of total chromatin accessibility versus nuclear volume per cell. Total chromatin accessibility per cell was calculated using the sum of UMIs per cell from snATAC-seq data of the MOp (*52*). (**H**) Scatterplot of the cell-type medians of the number of accessible chromatin regions versus nuclear volume per cell. The number of accessible chromatin regions per cell was obtained from snATAC-seq data of the MOp (*52*). Each dot in (F to G) corresponds to a cell type.

Next, we determined the 3D-genome organization for each identified cell type. We used 99-bit and 95-bit error-correcting Hamming codes to encode the 988 genome-wide loci and 965 super-enhancer loci, respectively, and imaged these 1953 loci using two back-to-back DNA-MERFISH measurements. In addition, we imaged the 28 marker-gene TSS loci using sequential multi-color DNA-FISH after DNA-MERFISH measurements, all on the same samples. We analyzed the DNA-MERFISH images by adapting the recently reported spatial genome aligner algorithm (*57*), which allowed us to determine the copy number of chromosomes and trace chromatin loci within each chromosome (Materials and Methods). This approach provided separate 3D structures of homologous chromosomes in individual cells (Fig. 1C). We found that the median pairwise distance between our imaged loci showed a high correlation with the pairwise contact frequency recently measured by single-nucleus methyl-3C (snm3C) (*29*) (fig. S3), demonstrating that the 3D chromosome structures were well preserved in their native tissue contexts in our integrated RNA- and DNA-MERFISH measurements.

### Cell-type-dependent variations in nucleus and chromosomal territory sizes and their relationship with transcriptional activity

We systematically characterized the size of the cell nucleus in each cell type in the MOp and observed substantially different nuclear sizes for different cell types, with non-neuronal cells, inhibitory neurons, and excitatory neurons exhibiting increasingly larger nuclear sizes (Fig. 1D and fig. S4). Within these major cell classes, different subclasses of cells also showed different nuclear sizes. For example, among excitatory neurons, L5 ET and L4/5 IT showed the largest and smallest nuclear sizes, respectively (Fig. 1D). Even among IT neurons, cells in different cortical layers showed different nuclear sizes (Fig. 1D). These results substantially expanded previous knowledge of different nuclear sizes for different cell types in the cortex (*58*).

Next, we investigated the relationship between the nuclear size and transcription by using a published snRNA-seq dataset (*52*) to obtain genome-wide RNA expression levels in different cell types in the MOp. Notably, we observed a strong correlation between the nuclear size and total transcriptional activity of the cell across different cell types (Fig. 1, D to F). Remarkably, the cell type with the largest nuclear sizes, L5 ET, showed a ∼10-fold higher total transcript counts than that observed in the cell types with the smallest nuclear sizes, i.e., microglia, oligodendrocytes, and endothelial cells (Fig. 1, D and E). Even among neurons, L5 ET showed a ∼3-fold greater total transcript counts than that observed in the neuronal cell types with the smallest nuclear size, the *Vip*-positive inhibitory neurons (Fig. 1, D and E). A similar trend in transcriptional activity changes across cell types was also observed in the human cortex (fig. S5) (*49, 59*). We also observed a strong correlation between the nuclear size and chromatin accessibility across different cell types (Fig. 1, G and H), using a published snATAC-seq dataset (*52*) to assess chromatin accessibility.

Next, we investigated whether transcriptional activity was also correlated with the physical sizes of chromosomal territories (fig. S6A). We observed that cell types with larger nuclear sizes generally had larger chromosomal territory sizes, except that most inhibitory neurons exhibited disproportionately larger chromosomal territory sizes compared to excitatory neurons with the same nucleus size (fig. S6, A and B). Consistent with this observation, inhibitory neurons displayed more intermixing of chromosome territories than excitatory neurons (fig. S6C). We also observed a positive correlation between the overall transcriptional activity of the chromosomes and the chromosomal territory sizes across different cell types (fig. S6D), although this correlation was weaker compared to the correlation with the nuclear sizes (Fig. 1F). A possible explanation for this weaker correlation is the stronger influence that each chromosome received from other chromosomes invading its territory in inhibitory neurons, which in turn contributes to transcription regulation. Consistent with this notion, excitatory neurons, which exhibited stronger segregation between chromosomal territories (fig. S6C), also showed higher correlation between transcriptional activity and chromosomal territory sizes (fig. S6E).

### Cell-type-dependent variations in higher-order chromosome structures and their relationship with transcriptional activity

The DNA-MERFISH data allowed us to further determine the higher-order chromosomal organization beyond the sizes of the cell nucleus and chromosomal territories. To this end, we computed the median pairwise distances between imaged loci within individual chromosomes (e.g., Chr1) in different cell types (Fig. 2A). We observed overall increased pairwise cis-chromosomal distances in cells with larger chromosomal territory sizes across different cell types (Fig. 2A; fig. S7). However, closer inspection of our data revealed a deviation from a simple decompaction model (Fig. 2B). Specifically, we observed that locus pairs with genomic distances greater than 10Mb showed increased pairwise spatial distances in cell types with larger chromosomal territory sizes (Fig. 2B), whereas locus pairs with genomic distances smaller than 2Mb often showed the opposite trend (Fig. 2B). We further computed the frequencies of spatial proximity (defined with a 3D distance threshold of 750 nm) between chromatin loci separated by different genomic distances, and observed divergent trends for proximity frequencies between locus pairs with large versus small genomic distances: while genomic loci separated by genomic distances greater than 10Mb showed increased proximity frequencies in cell types with smaller chromosomal territory sizes, the opposite trend was observed for genomic loci separated by genomic distances smaller than 5Mb (Fig. 2C).

**Fig. 2.**
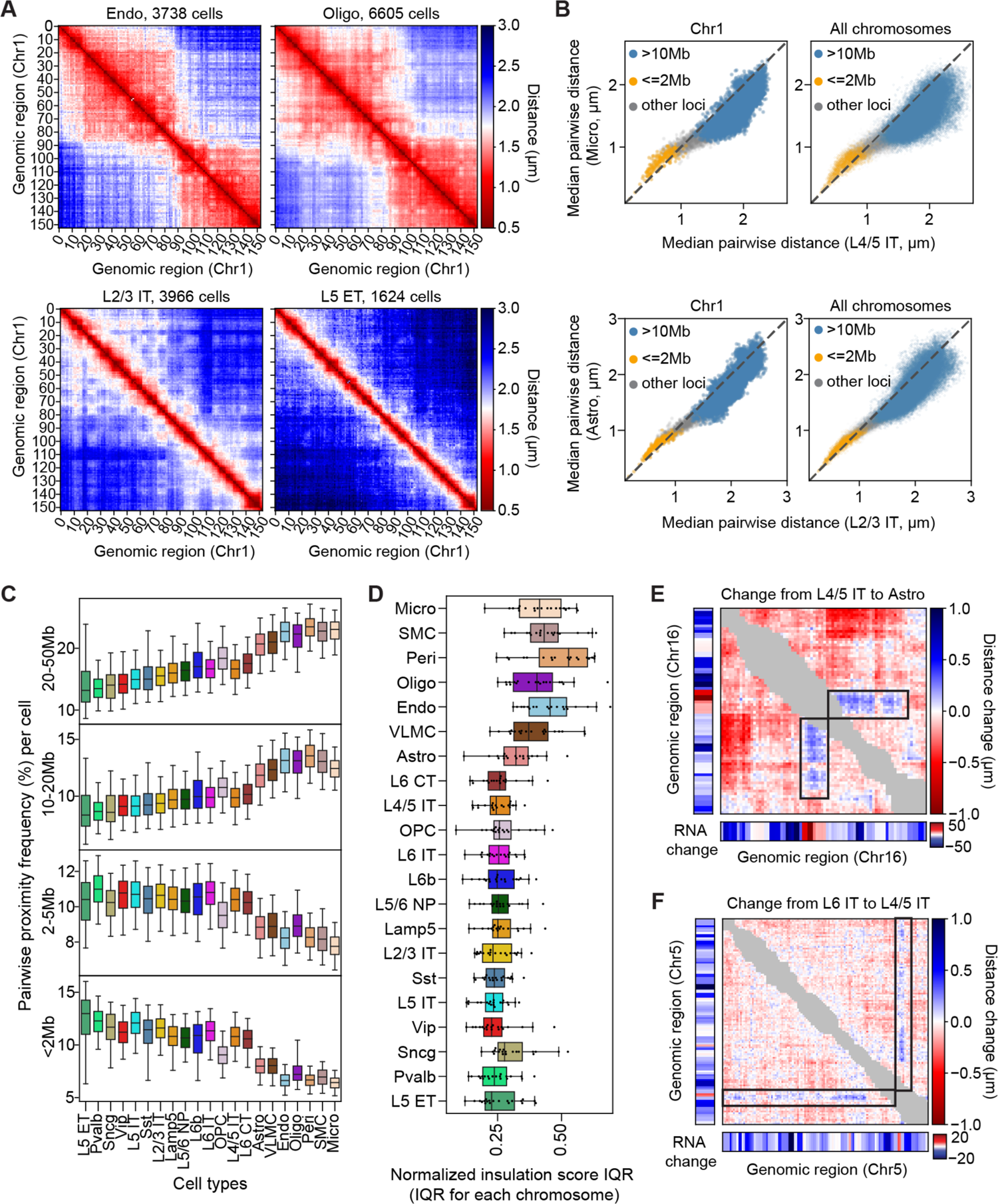
Cell-type-dependent variations in higher-order chromosome structures and transcriptional activity. (**A**) Median pairwise distance matrices for Chr1 (3.7Mb – 98.8Mb) in four example cell types. (**B**) Scatterplots of the cell-type medians of pairwise distances for cis-chromosomal locus pairs in one cell type versus another. Left: all locus pairs from Chr1. Right: all cis-chromosomal locus pairs from all chromosomes. Top: comparison between microglia and L4/5 IT neurons. Bottom: comparison between astrocytes and L2/3 IT neurons. Each dot in the scatterplot is colored according to the genomic distance of the indicated locus pair (orange: <2Mb; blue: >10Mb; light gray: all others). Dashed line indicates equality (y = x). (**C**) Boxplots showing the distributions of cis-chromosomal pairwise proximity frequency between genomic loci across all cells for each cell type, grouped by the genomic distances between individual locus pairs. Pairwise proximity frequency in each cell for the given genomic-distance group was calculated as the ratio of the number of proximal locus pairs (distance < 750nm) per cell over the number of all measured pairs per cell within the genomic-distance group. The center line, box, and whisker represent the median, 25th-75th percentile, and 5th-95th percentile, respectively. (**D**) Boxplots showing the distributions of normalized insulation score interquartile range (IQR) across all chromosomes for each cell type. Normalized insulation scores and the IQR were calculated as described in fig. S8B. (**E**) Comparison of higher-order chromosome organization and transcription in Chr16 (3.7Mb – 97.7Mb) between L4/5 IT and astrocytes. The matrix represents the change in median cis-chromosomal pairwise distance from L4/5 IT to astrocytes. Next to the matrix (left and below) are changes in transcript counts of the corresponding genomic loci from L4/5 IT to astrocytes (derived from snRNA-seq data (*52*)). Black boxes indicate genomic regions showing local changes between cell types that had an opposite trend to the overall changes of the whole chromosome. Gray elements near the diagonal represent locus-pairs whose genomic distances are <2Mb. (**F**) Similar to (E), but for comparison of higher-order chromosome organization and transcription in Chr5 (3.7Mb – 151.3Mb) between L6 IT and L4/5 IT neurons.

In addition to these differences in cis-chromosomal pairwise distances, we also observed large-scale structural differences between different cell types. In particular, when comparing non-neuronal cells and neurons, we observed that chromosomes in non-neuronal cells had a higher tendency to form large-size domains (Fig. 2A and fig. S8A) that resemble the “megadomains” in the transcriptionally inactive X chromosomes (*20, 22, 60–62*). We quantified the prevalence of these megadomain-like structures using the insulation score calculation (Materials and Methods), and observed more prominent insulation-score peaks, corresponding to the boundaries of megadomain structures, in chromosomes inside non-neuronal cells as compared to neurons (fig. S8B and Fig. 2D). Because non-neuronal cells generally have lower transcriptional activity than neurons (Fig. 1, E and F), we speculate that inactive chromatin may promote the formation of megadomain structures in non-neuronal cells, in a manner similar to how it occurs in inactive X chromosomes (*61, 62*).

To further explore the relationship between transcription and chromosomal organization, we also examined the differences in median pairwise distance matrices between cell types of different overall transcriptional activities. In most cases, these differential matrices showed unidirectional changes that were consistent with overall differences in chromosomal territory sizes and transcriptional activity: chromosomes in cell types with lower overall transcriptional activity showed more compaction (fig. S9). However, in some cases, we observed exceptions to this general trend (Fig. 2, E and F). For example, when comparing the median pairwise distance matrices of Chr16 between L4/5 IT neurons and astrocytes, we observed clear stripes running against the overall trend of the chromosome (Fig. 2E, indicated by black boxes). The chromatin region associated with these stripes showed notably higher transcriptional activity in astrocytes as compared to L4/5 IT neurons, despite that the chromosome-wise overall transcription activity level was lower in astrocytes (Fig. 2E). The physical distances between this region and the down-stream regions in the chromosome were also larger in astrocytes as compared to those observed in L4/5 IT neurons, even though astrocytes showed a general compaction of this chromosome as compared to L4/5 IT. We observed such “looping-out” of chromatin regions with relatively high transcriptional activity not only when comparing non-neuronal cell types with neurons, but also when comparing different neuronal cell types, for example, in Chr 5 between L4/5 IT and L6 IT (Fig. 2F). Mechanistically, it is possible that the increased physical separation between these regions and the other repressive regions in the chromosome helped increase the transcriptional activities of these regions.

### Cell-type-dependent variations in A/B compartmentalization and their relationship with transcriptional activity

We also observed notable differences in A/B compartment arrangements across different cell types, in particular, between neurons and non-neuronal cells. Specifically, we observed stronger plaid patterns in the median pairwise distance matrices of chromosomes in neurons (fig. S8A) and performed further analyses to study these features.

We applied standard Principal Component (PC) analysis to determine A/B compartments (*16*) in all major cell types. For each cell type, we calculated the normalized median pairwise distance matrix, computed the correlation matrix from this normalized distance matrix, performed PC analysis, and then selected the PC that showed the highest correlation with chromatin accessibility measured by snATAC-seq (*52*) to determine A/B compartments in that cell type (Materials and Methods). Interestingly, we found that the largest PC (PC1) did not always correspond to the A/B compartments in all cell types (Fig. 3A). For example, in excitatory neurons, the correlation matrix of Chr1 exhibited a pronounced plaid pattern, indicative of A/B compartment arrangements (Fig. 3A). Among all PCs, PC1 of the correlation matrix showed the highest correlation with chromatin accessibility (fig. S10). In contrast, in oligodendrocytes and oligodendrocyte progenitor cells, PC1 primarily corresponded to the megadomain structures described above, whereas PC2 showed the highest correlation with chromatin accessibility (Fig. 3A and fig. S10). Astrocytes displayed an intermediate behavior, where megadomain structures and A/B compartments were similarly prominent (Fig. 3A) and both PC1 and PC2 showed similar correlation coefficients with chromatin accessibility (fig. S10). These results indicate that the megadomain structures and A/B compartments, which represent different types of higher-order chromosomal structures, differ in strengths between different cell types, apparently counteracting each other. Given that neurons had higher overall transcriptional activities than non-neuronal cells (Fig. 1, E and F), the observation that neurons showed more prominent A/B-compartment chromatin segregation suggests a possibility that a stronger degree of A/B compartmentalization may cause higher transcriptional activity or vice versa, for example through the formation of transcriptional condensates (*6, 8*).

**Fig. 3.**
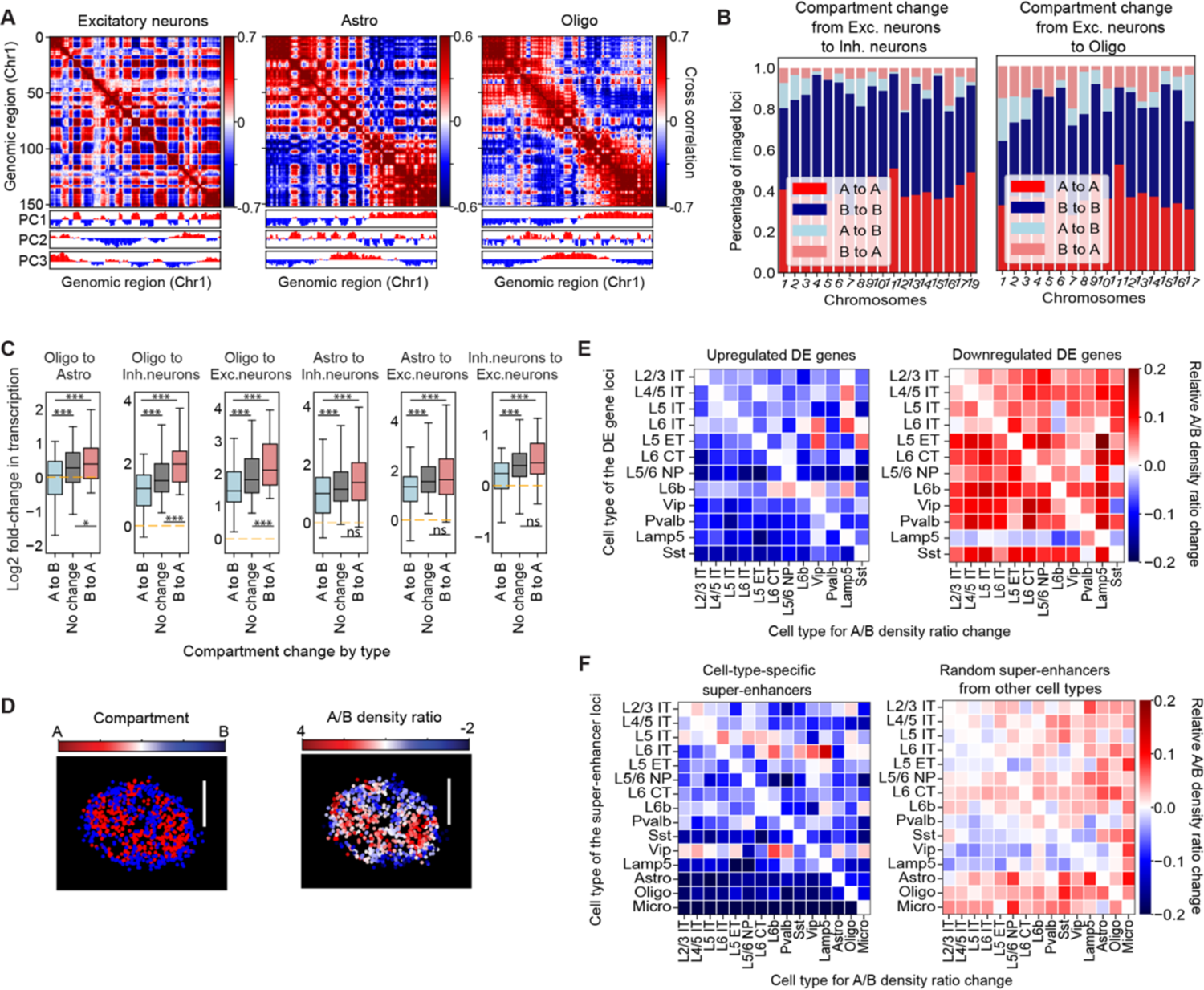
Cell-type-dependent variations in A/B compartment organization and transcriptional activity. (**A**) Cross-correlation matrices of Chr1 (3.7Mb – 98.8Mb) in several major cell types. Each element in the matrix represents the Pearson correlation coefficient between the indicated locus pair in terms of their normalized proximity frequency with all other loci from Chr1 (see Materials and Methods). The three rows below the matrices indicate the principal component (PC) values (positive values: red; negative values: blue). (**B**) Bar plots for the fraction of the A/B compartment conservations (e.g., A to A) and changes (e.g., A to B) from excitatory (Exc.) neurons to inhibitory (Inh.) neurons (left), or from excitatory neurons to oligodendrocytes and OPCs (right). Oligo here represents oligodendrocytes and OPCs together. (**C**) Boxplots for the distribution of the log2-fold change in transcription of individual genomic loci grouped by the A/B compartment conservations or changes from one cell type to another. The center line, box, and whisker represent the median, 25^th^-75^th^ percentile, and 5^th^-95^th^ percentile respectively. One-way ANOVA with Tukey post-hoc multiple comparison test was used for statistical analysis. ns: p>0.05; *: p<=0.05; ***: p<=0.001. (**D**) A single-cell image of compartment-A and B loci and their local A/B density ratio. Left: all decoded chromosome loci within a 2-µm z-plane in the nucleus, with compartment-A loci shown in red and compartment-B loci shown in blue. Right: same loci as in the left panel but colored by the local A/B density ratio. The local A/B density ratio is defined as the ratio of the local densities of trans-chromosomal A and B loci (i.e., A and B loci from other chromosomes, see Materials and Methods). Scale bar: 5 µm. (**E**) Changes in local A/B density ratio for cell-type-specific differentially expressed (DE) gene loci between the cell types where the DE genes were identified and other cell types. For each cell type, an imaged locus is considered a DE locus if there is a DE gene within 100kb of this locus. DE genes were identified for one cell type against all the other cell types using snRNA-seq data (*52*). Left: each pixel in the heatmap represents the median values of the fractional changes in the A/B density ratio of all upregulated DE gene loci for each cell type (x-axis) over the reference cell type (y-axis) where the DE genes were identified. Right: similar to the left panel, but for downregulated DE gene loci. (**F**) Similar to (E), but for changes in local A/B density ratio for cell-type-specific super-enhancer loci (left) or a random set of other super-enhancer loci selected regardless of cell-type identity (right). Cell-type-specific super-enhancers were selected by cell-type-specific long stretch snATAC-seq signals (*52*) (see Materials and Methods).

In addition to the different degree of overall A/B-compartment chromatin segregation in different cell types, we observed switching of genomic loci from one compartment to another when the cell type changed (Fig. 3B). Between excitatory and inhibitory neurons, ∼10-20% of genomic loci changed their compartment identities (A-to-B or B-to-A) (Fig. 3B, left). Likewise, between non-neuronal and neuronal cells, 10-30% of genomic loci changed their compartment identities (Fig. 3B, right). Interestingly, when changing from non-neuronal cells to neurons, or from inhibitory to excitatory neurons, transcriptional activity of genomic loci all tended to increase in general, regardless of whether the compartment identity of genomic loci switched from A to B, or from B to A, or did not switch (Fig. 3C). However, the genomic loci that switched from B to A compartment in general showed a greater increase in transcriptional activity than those that did not switch or switched from A to B compartments (Fig. 3C). These results suggest that A/B compartment rearrangement substantially influenced, but was not the sole determinant of, transcriptional activity change between cell types.

Next, we further explored the relationship between the local A/B chromatin environment and transcriptional activity of specific genes. To this end, we computed, for each imaged genomic locus, the local spatial densities of A and B chromatin loci near that locus and then determined the ratio of A-chromatin density over B-chromatin density as an indicator for the local A/B chromatin environment for that genomic locus in each cell (Fig. 3D; Materials and Methods) (*24*). This in turn allowed us to calculate, for each chromatin locus, the median of local A/B chromatin density ratio among all cells in any given cell type, which provided a quantitative proxy of the local A/B chromatin environment for each locus in each cell type.

We then investigated how this local A/B chromatin environment influenced differential expression of genes across different cell types. To this end, we first identified top differentially expressed (DE) genes (∼200 upregulated genes and ∼200 downregulated genes) in each cell type versus all other cell types using snRNA-seq data (*52*) (Materials and Methods). Among these top DE genes, we selected ones that were within 100kb of each of our imaged genomic loci, and then calculated the change in the median local A/B density ratio of this locus between the cell type where the DE gene was identified versus other cell types. Interestingly, we found that genes that were upregulated in a cell type in general tended to have higher local A/B density ratios in that cell type as compared to other cell types (Fig. 3E, left; fig. S11A; fig. S11B, left). Likewise, genes that were downregulated tended to have lower local A/B density ratios in that cell type (Fig. 3E, right; fig. S11A; fig. S11B, right).

We also examined the relationship between the local A/B chromatin environment and the activity of super-enhancers, which are known to contribute to cell-type-specific activation of gene expression (*55, 56*). For each imaged super-enhancer locus (which were all selected in a cell-type-specific manner), we examined the change in the local A/B density ratios between the cell type where the super-enhancer was identified and the other cell types. Similar to the trend observed for the DE genes, the super-enhancers also tended to have higher local A/B density ratios in their respective cell type as compared to other cell types (Fig. 3F, left; fig. S11C, left). In contrast, this trend was not observed in a randomization control where the super-enhancers were randomly selected regardless of cell-type identity (Fig. 3F, right; fig. S11C, right). Taken together, these results suggest that the local A/B-compartment chromatin environment plays a role in regulating cell-type-specific gene expression, not only for genes themselves, but also for gene regulatory elements.

### Cell-type-dependent variations in nuclear radial positioning of active and inactive chromatin

We calculated the normalized radial positions of our imaged loci (normalized by the radius of the nucleus in the same direction, Materials and Methods) and observed distinct radial positioning for different genomic loci in different cell types (Fig. 4A). In general, neurons and non-neuronal cells showed lower similarity in nuclear radial positioning profiles of genomic loci than the similarities observed among subtypes of neurons or among non-neuronal cell types (fig. S12).

**Fig. 4.**
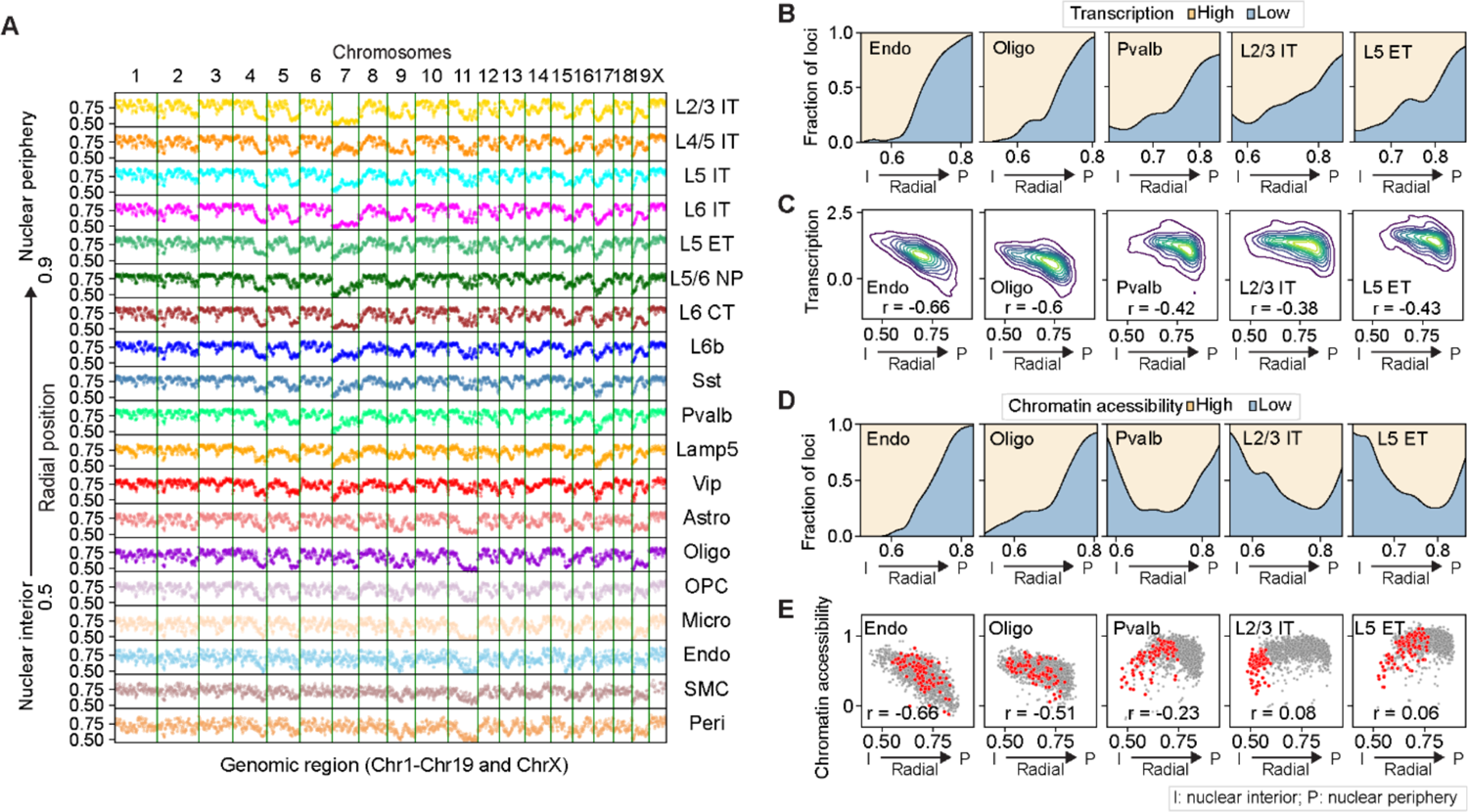
Cell-type-dependent variations in nuclear radial positioning of active and inactive chromatin. (**A**) Cell-type-median nuclear radial positions of all imaged loci (across Chr1-19 and ChrX, each column representing one chromosome) for individual cell types (rows). For each locus, nuclear radial position was calculated as the normalized distance of the locus to the centroid of the nucleus, with the maximum locus-to-centroid distance along the same direction normalized to 1. (**B**) The fractions of genomic loci with high and low transcriptional activity as a function of the normalized nuclear radial position for several cell types. Genomic loci were binned by their normalized radial positions. Within each bin, the fractions of high- and low-expression loci (top 25^th^ and bottom 25^th^ percentile, respectively, calculated from snRNA-seq data (*52*)) over the total counts of high- and low-expression loci were plotted. (**C**) 2D density plots of the cell-type medians of transcriptional activity of individual genomic loci versus the cell-type medians of normalized nuclear radial positions of the same loci. The density plot shows the kernel density estimation of the distribution of the data points. (**D**) Similar to (B) but for chromatin accessibility instead of transcriptional activity. The fractions of highly and lowly accessible loci were calculated from snATAC-seq data (*52*). (**E**) Scatterplots for the cell-type medians of chromatin accessibility of individual genomic loci versus normalized nuclear radial positions of the same loci. Gray dots represent loci from all chromosomes, and red dots represent loci from Chr7. The Spearman correlation coefficient r is indicated in (C) and (E).

Interestingly, we observed different degrees of correlation between transcriptional activity of genomic loci and their nuclear radial positions for different cell types. In non-neuronal cells, transcriptional activity of genomic loci was strongly correlated with their nuclear radial positions, with loci positioned closer to the nuclear periphery exhibiting lower transcriptional activity (Fig. 4, B and C). However, this correlation was weaker in neurons, accompanied by increased transcriptional activity for genomic loci near the nuclear periphery (Fig. 4, B and C). This difference between neurons and non-neuronal cells was even more pronounced for the correlation between chromatin accessibility and nuclear radial positioning (Fig. 4, D and E).

Moreover, the local A/B-chromatin environment of the imaged loci (as measured by their local A/B density ratio) were also less correlated with their nuclear radial positions in neurons as compared to non-neuronal cells (fig. S13). Interestingly, compared to excitatory neurons, inhibitory neurons showed consistently low levels of correlation between the nuclear radial positioning of genomic loci and their local A/B density ratios (fig. S13). Together, our results showed that active and inactive chromatin tended to be better segregated along the radial axis in non-neuronal cells than in neurons.

Interestingly, we also observed different degrees of correlation between transcriptional activity and nuclear radial positioning, and between chromatin accessibility and nuclear radial positioning, for different chromosomes (fig. S14, A to C). Notably, for some chromosomes, such as Chr7, the correlation between nuclear radial positioning and chromatin accessibility even showed opposite trends for non-neuronal cells and neurons (Fig. 4E; fig. S14, B and D). To understand the potential biochemical basis underlying these divergent behaviors between neurons and non-neuronal cells, we examined the histone modifications associated with our imaged loci, by examining single-cell joint histone modification and gene expression profiles obtained from the mouse cortex using Paired-Tag assays (*63*). We found that H3K9me3, a histone modification that marks heterochromatin (*64*), also showed opposite trends for radial-position dependence between neurons and non-neuronal cells (fig. S14, E and F). Likewise, such opposite trends were also observed for H3K27ac (fig. S14F), a histone modification that marks active chromatin (*64*). Therefore, compared to non-neuronal cells, neurons have more active chromatin in the nuclear periphery and more inactive chromatin near the nuclear interior, in part due to the inverted radial organization of a subset of chromosomes.

Along with the increased transcriptional activity of genes near the nuclear periphery in neurons, we observed that the majority of the transcriptionally active genomic loci in the nuclear periphery of neurons contained actively expressed long genes (>300kb) (fig. S15A), by combining published snRNA-seq data (*52*) with our spatial data from DNA-MERFISH (Materials and Methods). Many of these long genes are associated with gene ontology (GO) terms that are related to synapse functions (fig. S15, B and C). In the nuclear interior, although long genes were not preferentially activated over short genes in neurons, there were still more transcriptionally active loci containing actively expressed long genes in neurons as compared to non-neuronal cells (fig. S15A).

### Radial-position-dependent transcriptional regulation by MeCP2

Our observations showed that neurons and non-neuronal cells have different radial organization of active and inactive chromatin in the nucleus. To further understand the neuron-specific radial organization of chromatin in the nucleus, we compared the patterns of several active and repressive chromatin marks and chromatin-binding proteins across all imaged loci (Fig. 5, A and B). Interestingly, among all factors that we examined, the binding of the Methyl-CpG binding protein 2 (MeCP2) (*65*) consistently showed a high correlation with the nuclear radial positioning of chromatin in neurons, with a stronger tendency of MeCP2 binding to chromatin loci closer to the nuclear center (Fig. 5, A and B).

**Fig. 5.**
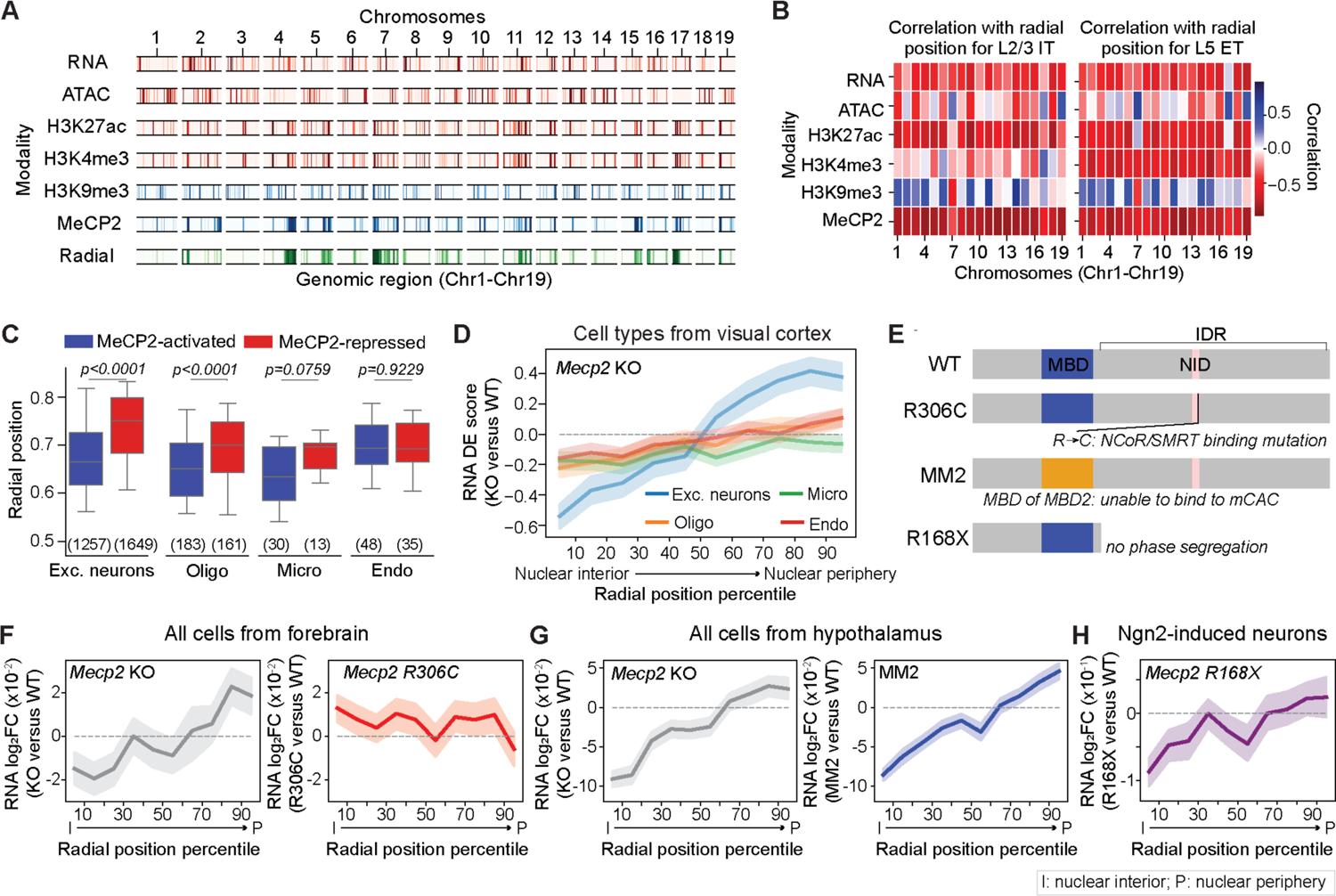
Nuclear-radial-position dependent transcriptional regulation by MeCP2. (**A**) Transcriptional activity (RNA), chromatin accessibility (ATAC), histone modifications (H3K27ac, H3K4me3, and H3K9me3), MeCP2-binding, and nuclear radial positioning for individual genomic loci on Chr1-19. For radial positioning, loci near the nuclear interior (35th percentile) are highlighted in color. For the other plots, loci enriched for the indicated signals (top 30th percentile) are highlighted in color. Transcriptional activity and chromatin accessibility were calculated from Ref. (*52*); H3K27ac, H3K4me3, and H3K9me3 levels were calculated from Ref. (*63*); MeCP2-binding level was calculated from Ref. (*65*). (**B**) Correlation of transcriptional activity, chromatin accessibility, histone modification levels, and MeCP2-binding level with nuclear radial positioning for L2/3 IT (left) and L5 ET neurons (right). The Spearman correlation coefficient between the radial positions and the indicated quantities of all loci on the indicated chromosome are shown. (**C**) Boxplots for nuclear radial positions of genes grouped by their transcriptional activity changes upon *Mecp2* deletion for various major cell types. Only genes that were significantly differentially expressed between male *Mecp2 +/y* (wild-type, WT) and *Mecp2-/y* (knock-out, KO) mice (adjusted *p*<0.05), determined using data from Ref.(*71*), are plotted. Numbers on the bottom indicate the number of significant DE genes. *P* values indicate the statistical significance calculated by Mann Whitney U test, corrected by the Benjamini-Hochberg procedure. (**D**) Transcriptional changes upon *Mecp2* deletion versus the normalized nuclear radial positioning of the genomic loci for various cell types. For each gene, the radial position is approximated by that of the closest imaged locus and all mapped genes are grouped into 10 equal bins by their normalized nuclear radial positions. The transcriptional level change (RNA DE score) is defined as the t-statistic score for differential expression when comparing the expression-level distributions of *Mecp2* KO and WT cells. The mean (line) and 95% confidence interval (shade) for the DE scores in each bin are plotted. (**E**) Schematics of Mecp2 mutations/transgenes analyzed in (F to H). MeCP2 contains a methyl-CpG (mCG) binding domain (MBD, blue) and a NCoR/SMRT-interacting domain (NID, pink). *Mecp2* R306C mutation in the NID (*68, 73*) abolishes its interaction with the NCoR/SMRT complex, which is a transcriptional repressor. MM2 transgene (*74*) replaces the MBD of MeCP2 with the mCG-binding domain of the MBD2 protein, which abolishes MeCP2’s ability to bind to methylated CAC (mCAC) but not mCG. *Mecp2* R168X mutation (*41*) truncates the intrinsically disordered region (IDR) in MeCP2, which abolishes its liquid-liquid phase segregation ability. (**F**) Similar to (D), but for the transcriptional changes (log_2_ fold change (FC) of RNA counts) upon *Mecp2* KO (left) or *Mecp2* R306C mutation (right) measured in the mouse forebrain (*73*). (**G**) Similar to (F), but for the transcriptional changes upon *Mecp2* KO (left) or upon introduction of the *Mecp2* MM2 transgene (right) in mouse hypothalamus (*74*). (**H**) Similar to (F), but for the transcriptional changes upon *Mecp2* R168X mutation in Ngn2-induced cultured neurons (*41*).

MeCP2 plays important roles in regulating gene transcription (*66–71*) and mutations of the *MECP2* gene cause the Rett syndrome, a progressive neurodevelopmental disorder (*39*). Considering that MeCP2 is highly expressed in neurons (*72*), we next asked whether the correlation between MeCP2 binding and nuclear radial positioning of chromatin is related to the transcriptional regulation by MeCP2 in neurons. As a first step, we analyzed published scRNA-seq data from wild-type (WT, *Mecp2 +/y)* and *Mecp2* knock-out (KO, *Mecp2-/y)* male mice (*71*). We focused on four major cell types from this dataset: excitatory neurons, oligodendrocytes, microglia, and endothelial cells because the number of inhibitory neurons and astrocytes measured in this dataset is relatively low. We found that genes downregulated upon *Mecp2* deletion (genes activated by MeCP2) preferentially localized closer to the nuclear center as compared to the genes upregulated upon *Mecp2* deletion (genes repressed by MeCP2) (Fig. 5C). Moreover, we systematically quantified the nuclear-radial-position dependence of transcription regulation by MeCP2 by calculating the mean differential-expression (DE) scores (t-statistics score) upon *Mecp2* deletion for all genes in these four cell types, using our imaging results to infer nuclear radial positions of the genes. Remarkably, we found that excitatory neurons showed a strong nuclear-radial-position dependence of transcriptional regulation by MeCP2: transcription of genes at the nuclear periphery were repressed by MeCP2 whereas genes near the nuclear interior were transcriptionally activated by MeCP2 (Fig. 5D). Interestingly, compared to excitatory neurons, the degree of transcriptional up- or down-regulation by MeCP2 was much smaller in non-neuronal cells (Fig. 5D).

To gain more insights into the molecular mechanisms underlying the radial-position-dependent effect of MeCP2, we further examined the radial-position-dependent effects of various *Mecp2* mutations by analyzing bulk RNA sequencing data (Fig. 5E). As an approximation, we mapped the genes in published bulk RNA sequencing data of *Mecp2* knockout and three *Mecp2* mutations (R306C, MM2 and R168X) (*41, 73, 74*) to the nuclear radial positions of the closest imaged genomic loci that we imaged. We observed a radial-position-dependent effect of transcriptional activity change upon *Mecp2* deletion in both forebrain and hypothalamus (Fig. 5, F and G, left), the general trend of which was similar to that observed for neurons in the scRNA-seq data (Fig. 5D). Next, we examined the effect of the R306C mutation in *Mecp2*, which disrupts MeCP2 binding to the transcriptional co-repressor NCoR/SMRT-complex (*68, 73*) (Fig. 5F, right). In contrast to *Mecp2* deletion, the R306C mutation caused a more moderate and largely radial-position-independent increase in transcriptional activity (Fig. 5F, right), indicating that the transcriptional upregulation by wild-type MeCP2 in the nuclear interior, as reflected by the decrease in transcriptional activity upon *Mecp2* deletion, does not require interaction with the NCoR/SMRT-complex. Likewise, the increase in transcriptional activity by the R306C mutation in the nuclear periphery was weaker than that by *Mecp2* deletion, suggesting that transcriptional repression by wild-type MeCP2 in the nuclear periphery is not fully accounted for by MeCP2’s ability to recruit the NCoR/SMRT co-repressor.

MeCP2 can bind to both methylated CG and methylated CA, with stronger affinity to mCAC than other forms of mCAH trinucleotides. To understanding which binding mode of MeCP2 is important for its radial-position-dependent transcriptional regulation effect, we analyzed bulk RNA sequencing data on the knock-in mice *Mecp2-MBD2* (MM2), which specifically abolishes MeCP2-binding to mCAC but not mCG by replacing the methylated DNA-binding domain of Mecp2 with that of the MBD2 protein (*74*) (Fig. 5E). Our analysis results showed that the MM2 transgene recapitulated the radial-position-dependent effect of *Mecp2* deletion on transcription (Fig. 5G, right).

MeCP2 is also a key component of the heterochromatin phase-separated condensates (*41, 42*). Our analysis of the RNA sequencing data on mice expressing the *Mecp2* truncation R168X (*41*), which lacks the intrinsically disordered region of MeCP2 required for liquid-liquid phase separation (Fig. 5E), showed that this mutation had a similar radial-position-dependent effect on transcription as seen in for *Mecp2* deletion (Fig. 5H).

Taken together, our analysis results suggest that MeCP2 regulates gene expression in a radial-position-dependent manner through its binding to mCAC and likely through the formation of phase-separated condensates. These molecular mechanisms promote transcriptional activation of genes in the nuclear interior and repress gene expression at the nuclear periphery. In addition, it has been shown previously that the ability of MeCP2 to recruit the repressive NCoR/SMRT-complex also contributes to MeCP2’s role in transcriptional repression (*68, 73*), and our data suggest that this latter effect occurs in a manner that is largely independent of the nuclear radial position. It is interesting to note that the R306C mutation confers milder symptomatic features in human patients than the R168X mutation or *MECP2* large DNA deletions (*75*). Hence, the radial-positioning-dependent transcriptional regulation by MeCP2 may be particularly important for the clinical severity in Rett syndrome.

### Effects of MeCP2 on nuclear radial positioning of chromatin and active/inactive chromatin segregation

Given our observations of the correlation between nuclear radial positioning of genomic loci and their local A/B chromatin environment and transcriptional activity (Fig. 4 and fig. S13), and the radial-position-dependent transcriptional regulation by MeCP2 (Fig. 5, C to H), we wondered whether MeCP2 influences the nuclear radial positioning of the chromatin loci and their local A/B-compartment chromatin environment. To address these questions, we applied integrated RNA- and DNA-MERFISH imaging to measure chromatin organization in the MOp of *Mecp2 +/-* heterozygous female mice. Since the *Mecp2* gene is on the X chromosome (*39, 76*) and one of two X chromosome homologs in female mice is randomly inactivated during development (*77*), we anticipate that roughly half of the cells in *Mecp2 +/-* mice would express *Mecp2* from the WT allele and the remainder cells would not express *Mecp2* due to the KO allele. Through immunostaining of MeCP2 conducted in conjunction with MERFISH measurements on the same samples, we identified WT cells and *Mecp2* KO cells across different cell types (Fig. 6, A and B; fig. S16A). *Mecp2* deletion did not significantly change the cell type composition in the MOp (fig. S16B), nor did it change the size of the nucleus substantially, with only a subset of cell types exhibiting statistically significant but subtle changes in the nuclear volume (fig. S17).

**Fig. 6.**
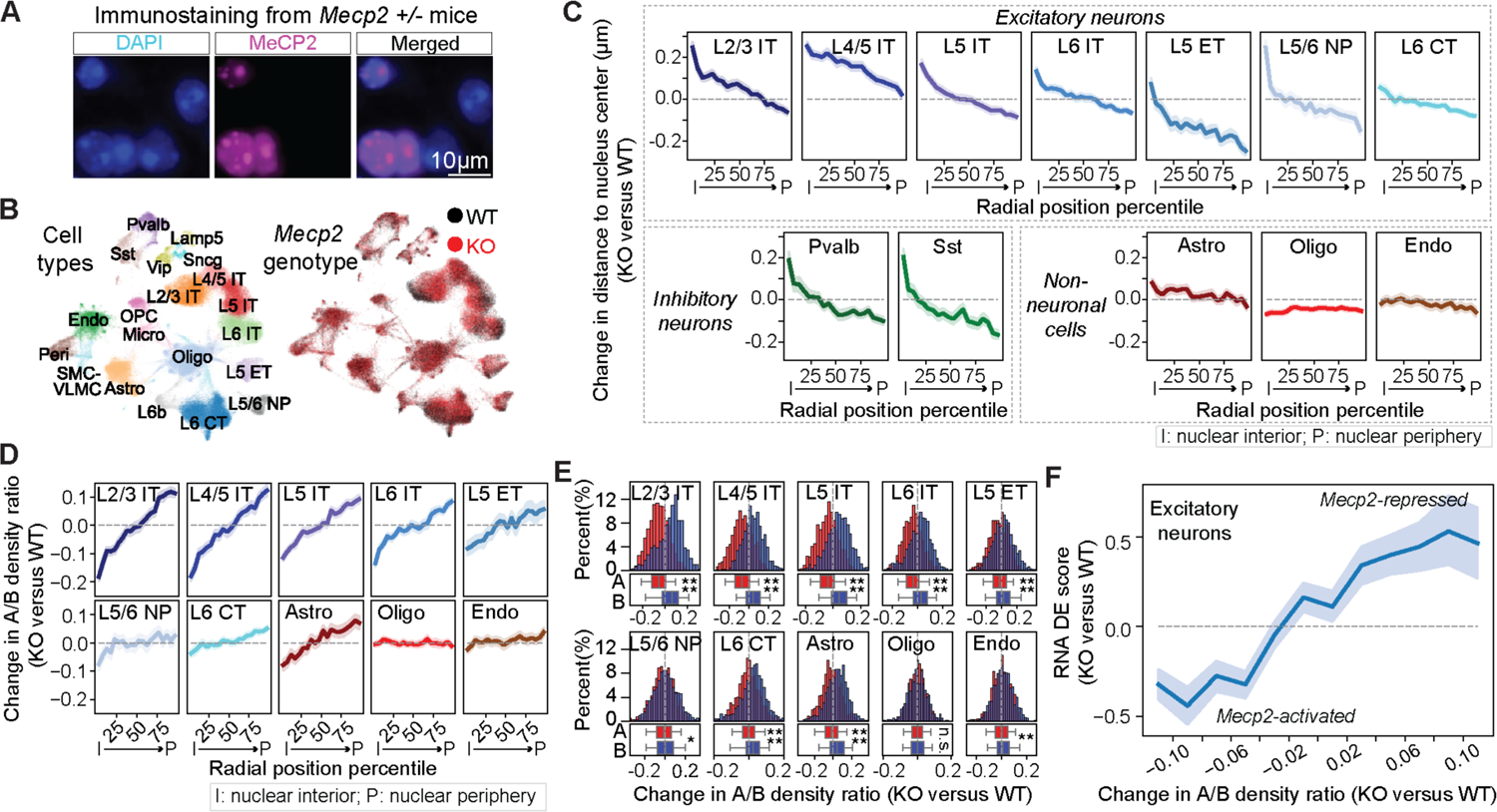
MeCP2-mediated A/B-compartment segregation regulates local chromatin environment in a radial-position and cell-type-dependent manner. (**A**) Representative images of DAPI (cyan) and MeCP2 (magenta) staining of a small region in the MOp in *Mecp2* +/- female mice. (**B**) UMAP representation of cell types (left) and genotypes (right) in the MOp of *Mecp2 +/-* female mice. (**C**) Nuclear radial position changes upon *Mecp2* deletion as a function of the normalized radial position of the genomic loci in various cell types. All imaged loci were grouped into 20 equal bins based on their cell-type median normalized radial positions in the WT cells. (**D**) Similar to (C), but for the changes in local A/B density ratio upon *Mecp2* deletion. (**E**) Histograms of local A/B density ratio changes upon *Mecp2* deletion for compartment-A (red) and compartment-B (blue) loci in various cell types. The corresponding boxplots for the distributions are shown at the bottom. (**F**) Correlation between transcriptional changes and local A/B density ratio changes upon *Mecp2* deletion in excitatory neurons. Genes are grouped based on their local A/B density ratio changes indicated in the x-axis. For each gene, the differential expression (DE) score for transcription change between KO and WT cells was calculated as in Fig. 5D, using scRNA-seq data (*71*). For the line plots (C, D, F), the line and shaded area represent the mean and the 95% confidence interval, respectively. For boxplots in (E), the center line, box, and whiskers represent the median, 25^th^-75^th^ percentile, and 5^th^-95^th^ percentile, respectively. Student’s t-tests with Bonferroni corrections were used for statistical significance evaluation in (E). *: p<=0.05; ***: p<=0.001; ****: p<=0.0001; n.s.: p>0.05.

Next, we systematically characterized the chromatin organization changes upon *Mecp2* deletion in different cell types. We focused our nuclear-radial-positioning analysis on several abundant cell types (fig. S18A). Notably, we observed a tendency for centrally-located chromatin loci to move towards the nuclear periphery and peripherally-located chromatin loci to move towards the nuclear center upon *Mecp2* deletion in most excitatory neurons and *Pvalb* and *Sst* inhibitory neurons (Fig. 6C; fig. S18B). Such movement was not observed in oligodendrocytes and endothelial cells and only to a small extent in astrocytes (Fig. 6C; fig. S18B).

Next, we examined the effect of *Mecp2* deletion on the local A/B-compartment chromatin environment of the imaged genomic loci. The cross-correlation matrices used to identify A/B compartments were highly similar between WT and *Mecp2* KO cells (fig. S19A), and the degree of similarity was particularly high for excitatory neurons, astrocytes, oligodendrocytes, and endothelial cells, but slightly lower for inhibitory neurons and microglia (fig. S19B). However, due to the low number of cells sampled for inhibitory neurons and microglia, we could not conclude whether these differences reflect real A/B compartment changes or noise. Therefore, we focused on the relatively abundant excitatory neurons, astrocytes, oligodendrocytes, and endothelial cells. Because of the high similarity between compartment boundaries observed between WT and *Mecp2* KO cells in these cell types, we used our A/B compartment assignment from the WT MOp datasets for further analysis.

Interestingly, similar to its effect on nuclear radial positioning, *Mecp2* deletion affected the local A/B chromatin environment of the imaged genomic loci most strongly in IT neurons, to a less extent in other excitatory neurons and astrocytes, but did not have any effect in oligodendrocytes and endothelial cells (Fig. 6D). In IT neurons, compartment-A loci showed a stronger tendency to move from the nucleus center outward to nuclear periphery compared to compartment-B loci, and this difference was reduced in other excitatory neurons and astrocytes, and further reduced or not observed in oligodendrocytes and endothelial cells (fig. S20). Consistent with these observations, the local A/B density ratio in excitatory neurons decreased for loci near the nuclear center and increased for loci at the nuclear periphery (Fig. 6D). This change was most pronounced in IT neurons, weaker in other excitatory neurons and astrocytes, and largely absent in oligodendrocytes and endothelial cells (Fig. 6D).

In line with the above observations, in excitatory neurons, compartment-A loci generally showed a decrease in local A/B density ratio and compartment-B loci generally showed an increase in local A/B density ratio (Fig. 6E), suggesting that A loci were relocated to a more heterochromatic environment and B loci were relocated to a more active chromatin environment upon *Mecp2* deletion. This trend was again the strongest in IT neurons, weaker in other excitatory neurons and astrocytes, but largely absent in oligodendrocytes and endothelial cells (Fig. 6E). Thus, *Mecp2* deletion led to an overall decrease in the degree of A/B-compartment chromatin segregation in excitatory neurons.

Finally, we observed a strong correlation between the changes in local A/B density ratio and transcriptional activity induced by *Mecp2* KO (Fig. 6F). Upon *Mecp2* deletion, the loci that showed an increase in local A/B density ratio also showed an increase in RNA expression level. Conversely, the loci that showed a decrease in local A/B density ratio showed a decrease in RNA expression level. Therefore, our data suggest that MeCP2 regulates the chromatin organization in neurons, in particular the nuclear radial positioning and A/B compartment segregation of chromatin, and through these effects, MeCP2 regulates transcription in excitatory neurons. Upon *Mecp2* deletion, the degree of A/B-compartment chromatin segregation was weakened, the nuclear interior became a less active environment, and the nuclear periphery became less repressive environment. This provides a uniform, mechanistic explanation for the apparently divergent effect of MeCP2 deletion on transcriptional activity, namely the decrease in transcription for genomic loci located in the nuclear interior and increase in transcription for genomic loci located in the nuclear periphery.

## Discussion

In this work, we developed an integrated RNA and DNA genome-scale imaging method to identify cell types and determine cell-type-specific 3D-genome organization in complex tissues. We applied this approach to study the primary motor cortex in the mouse brain and revealed a wide spectrum of nuclear-architecture differences among different brain cell types, including differences in the cell-nucleus sizes, chromosomal-territory sizes, higher-order chromatin structures, and nuclear radial positioning of chromatin. We further observed strong correlations of these cell-type-dependent variations in nuclear architecture and 3D-genome organization with transcriptional regulation, in terms of both the overall transcriptional activity of the cells and the differential expression of specific genes between cell types. Furthermore, our studies provided mechanistic insights into how MeCP2, a methylated-DNA binding protein, regulates the 3D-genome organization and transcription in neurons.

First, we observed that different cell types in the brain exhibit substantially different nuclear sizes and that the overall transcriptional activity of cells vary dramatically in a manner that is strongly correlated with the nuclear sizes (Fig. 1, D to F). In particular, both the nuclear size and the transcriptional activity increased from non-neuronal cells to inhibitory neurons, and then to excitatory neurons, and finer changes were also observed among subclasses of cells within these major cell classes. Our results suggest that the size of the cell nucleus could be a hallmark of the cell types and states. In disease contexts such as cancer, alterations in the nucleus size have been frequently reported and used as diagnostic markers (*78*). It is therefore possible that these disease-related changes in nuclear sizes also influence gene expression in a manner that is relevant to disease phenotypes.

Interestingly, as the nuclear size increased from non-neuronal cells to neurons, chromatin was not simply decompacted in a uniform manner. Instead, we observed a divergent trend in cis-chromosomal proximity for chromatin loci separated by different genomic intervals (Fig. 2, A to C), consistent with sequencing-based studies of the mouse and human brains reporting that non-neuronal cells and neurons exhibit distinct preferences for cis-chromosomal contacts at different genomic distances (*28, 29, 31*). In particular, despite their larger nucleus and chromosomal territory sizes as compared to non-neuronal cells, neurons exhibited a higher frequency of contact or spatial proximity for loci separated by short genomic intervals (<2Mb). This phenomenon may be attributed to the active transcriptional regulatory mechanisms in neurons such as the increased frequency in the formation of gene-body domain and enhancer-promoter loops (*26, 29, 79*).Interestingly, during the cell cycle as cells progress from G1 to G2/S phases, the transcriptional upregulation is also accompanied by an increase in the contact frequency of chromatin loci separated by short genomic intervals, whereas the contact frequency of long-genomic-interval locus pairs decreases (*80*). Thus, the increased interactions between loci separated by short-genomic intervals and decreased interactions between loci separated by long-genomic intervals maybe be a general signature associated with global transcription upregulation and nuclear volume increase in a variety of conditions, such as cell-cycle progress and cell-type differentiation (*31, 80*).

Notably, in non-neuronal cells, which exhibited relatively low overall transcriptional activity, chromosomes tended to form large-sized domain structures (Fig. 2, A and D), which resembled the megadomains previously observed in the transcriptionally inactive X chromosomes (*20, 22, 60–62*). Megadomain formation in inactive X chromosomes is mediated by the prevalent repressive epigenetic signatures, which compact heterochromatin into large domains separated by insulating elements (*61, 62*). Whether a similar mechanism causes the formation of megadomain structures in autosomes in non-neuronal cells remains an open question. By contrast, megadomain structures were diminished in neurons, which instead exhibited more prominent A/B compartments (Fig. 3A). The formation of megadomain structures and A/B compartments thus appear to counteract each other. Such an opposing effect has also been reported during X chromosome inactivation (*22, 61*). We hypothesize that the switching between megadomain and A/B-compartment structures are causally linked to transcriptional regulation, and that this switching is a general mechanism applicable to not only X chromosomes but also autosomes.

We also observed a correlation between the local A/B-compartment chromatin environment and transcriptional activity changes of specific genes between cell types (Fig. 3, E and F). Previous studies suggest that local A/B compartmentalization is associated with differential gene expression during brain development and aging (*28, 30*). Here, we showed that genes differentially expressed between different cell types were in general more strongly expressed in cell types where the local A/B chromatin density ratio of the genomic locus was higher. We observed the same trend for enhancer activity -- super-enhancers also tended to be more active in cell types where their local A/B density ratio was higher. The local enrichment for active chromatin could result in a higher local concentration of transcription machinery and activation factors, thereby enhancing the activity of these genes and enhancer elements. Conversely, it is also possible that active promoters and enhancers tend to colocalize with each other, for example, through the formation of transcriptional condensates (*6, 8*), thereby causing a higher local concentration of active chromatin.

Another major difference in 3D-genome organization that we observed between different cell types was the nuclear radial positioning of active and inactive chromatin. While non-neuronal cells exhibited a strong correlation between nuclear radial positioning of chromatin loci and their accessibility and transcriptional activity, with active and inactive chromatin respectively residing in the nuclear interior and periphery, neurons exhibited a more intermixed organization of active and inactive chromatin along the radial axis of the nucleus (Fig. 4, B to E). The widespread upregulation of gene expression in neurons may contribute to the enrichment of active chromatin in the nuclear periphery. The majority of active genomic loci near the nuclear periphery of neurons contain long genes (fig. S15A), which are likely more capable of concentrating transcriptional machinery (*81*), potentially overriding the repressive mechanisms in the nuclear periphery. In line with our observations, it has been shown recently that lamina-associated domains are less repressive in neurons than in cultured embryonic stem cells (*82*). Upregulation of transcription near the nuclear periphery may also weaken the tethering of inactive chromatin to the nuclear envelope (*83, 84*), causing its relocation to the nuclear interior.

Finally, we observed that MeCP2 regulates both chromatin organization and transcriptional activity in a nuclear-radial-position and cell-type dependent manner (Fig. 5 and Fig. 6). Notably, MeCP2 repressed transcription of genes residing near the nuclear periphery and activated transcription of genes near the nuclear center, and these effects were strongest in excitatory neurons and nearly absent in endothelial cells and oligodendrocytes (Fig. 5). Structurally, MeCP2 promoted local A/B-compartment chromatin segregation: upon *Mecp2* deletion, the originally active environment of nuclear interior underwent a decrease in local A/B chromatin density ratio, becoming less active, whereas the originally more repressive environment of nuclear periphery underwent an increase in the local A/B chromatin density ratio, becoming less repressive (Fig. 6). This role of MeCP2 in chromatin organization provides a natural explanation for the radial-position-dependent transcription regulation by MeCP2 and resolved a long-standing puzzle of why MeCP2 activates transcription of some gene and represses transcription of other genes (*43, 44, 67, 69–71*).

MeCP2 may enhance A/B-compartment chromatin segregation by promoting heterochromatin condensation. It has been shown previously that MeCP2 can cause liquid-liquid phase separation (*41, 42*) and that MeCP2-positive heterochromatin condensates do not coalesce with Brd4 and Med1 condensates that are enriched for active chromatin (*41*). This exclusion effect between MeCP2 condensates and transcription condensates could provide a mechanism to explain why loss of MeCP2 leads to increased intermixing between active chromatin and inactive chromatin, as we observed, and thereby causes transcription mis-regulation. Indeed, we observed that truncation of the intrinsically disordered domain in MeCP2, which abolishes its ability to form condensates (*41*), recapitulated the radial-position-dependent effect on transcription by *Mecp2* deletion (Fig. 5H). Since neurons exhibited a more intermixed organization of active and inactive chromatin along the radial axis (Fig. 4), the expression of MeCP2 may be particularly important for preventing unwanted crosstalk between active and inactive chromatin and thereby ensuring proper expression of genes in neurons. This cell-type-dependent effect of MeCP2 may thus reflect a functional adaptation to cell-type-specific 3D-genome organization.

In summary, our integrated RNA- and DNA-MERFISH studies provided rich insights into the connections between cell-type-specific 3D-genome organization and transcription regulation both in normal brain function and in disease conditions caused by *Mecp2* mutations. The ability of this technology to determine cell-type-specific 3D-genome organization in native tissues can be broadly applied to advancing our understanding of gene-expression regulation in health and in disease. We anticipate that future studies combining 3D-genome and transcriptome imaging with epigenome imaging (*85*) and protein imaging could further enhance our ability to understand the inter-relationship between epigenetic properties of chromatin, gene-regulation factors, chromatin structures, and transcription regulation. Moreover, combination of this approach with live-cell imaging, for example by CRISPR-based live-cell chromatin imaging (*86*) followed by fixed-cell genome-scale imaging of the same sample, and with spatial manipulations of genomic loci by CRISPR-based methods (*87*), could further advance our ability to study the causal relationship between 3D-genome organization and transcription regulation with high spatiotemporal resolution.

## Acknowledgements

We thank Xingjie Pan and Meng Zhang for their help with RNA-MERFISH analyses. We thank all members of the Zhuang lab for helpful discussions and suggestions.

## Funding

This work is in part supported by the National Institutes of Health (UM1HG011585 to B.R. and X.Z.). X.Z. is a Howard Hughes Medical Institute Investigator.

## Author contributions

S.L., P.Z., C.Y.W., B.R., and X.Z. designed the experiments. S.L., P.Z., and C.Y.W. performed the experiments. S.L., P.Z., C.Y.W. and B.B.J. analyzed the data. N.R.Z. helped with the selection of cell-type-specific super-enhancers. S.L., P.Z., C.Y.W. and X.Z. interpreted data and wrote the manuscript with input from B.B.J., N.R.Z. and B.R.

## Competing interests

X.Z. is an inventor of patents applied for by Harvard University related MERFISH and a co-founder and consultant of Vizgen, Inc. B.R. is a co-founder and consultant of Arima Genomics, and a co-founder of Epigenome Technologies, Inc.

## Data and materials availability

Oligonucleotide probe sequences used for this study can be found in Tables S1, S2, and S3, and they can be purchased from commercial sources. Data reported in this paper are available at 4DN Data Portal (https://data.4dnucleome.org/; accession numbers: 4DNESPE924IP and 4DNESMTNNB3N). Analysis code used for MERFISH probe library design, MERFISH image decoding, and data quantification is available at: https://github.com/ZhuangLab/Chromatin_Analysis_2023.

## Supplementary Materials for

### Materials and Methods

#### Animals

Three adult C57BL/6 male wild-type (WT) mice aged 57-63 days were used for investigating the cell-type-dependent chromatin organizations in the mouse primary motor cortex (MOp). Two adult Mecp2 heterozygous mutant female (*Mecp2 +/-*) mice aged 11 weeks, and three adult Mecp2 heterozygous mutant female (*Mecp2 +/-*) mice aged 27 weeks were used for RNA-MERFISH combined with Mecp2 immunostaining. Among these mice, brain slices from two 11-week-old and two 27-week-old mice were further used for DNA-MERFISH imaging to investigate the effect of Mecp2 on chromatin organizations. Animals were maintained on a 12 hour:12 hour light/dark cycle (2pm-2am dark period), at a temperature of 22 ±°C, a humidity of 30–70%, with ad libitum access to food and water. Animal care and experiments were carried out in accordance with NIH guidelines and were approved by the Harvard University Institutional Animal Care and Use Committee (IACUC).

#### Tissue preparation for integrated RNA-MERFISH and DNA-MERFISH

Mice were euthanized with CO2, and their brain was quickly harvested and frozen immediately in optimal cutting temperature compound (Tissue-Tek O.C.T.; VWR, 25608-930) in dry ice and stored at −80 °C until sectioning. Frozen brains were sectioned at −18 °C on a cryostat (Leica CM3050s). Slices were removed and discarded until the MOp region was reached. Specifically, a continuous set of 10-µm-thick serial coronal sections of the brain were cut from anterior to posterior to include the MOp region, according to Allen Institute reference map v3 (https://atlas.brain-map.org/atlas?atlas=602630314) (*88*), as previously described (*50*).

Typically, every 2 slices that are ∼100-µm apart along the anterior-to-posterior axis were collected onto one #1.5 round coverslip (Bioptechs, 0420-0323-2). Note that for all prepared WT tissues, only coverslips that were successfully imaged for both RNA-MERFISH and DNA-MERFISH were processed for downstream image analysis. In total, for WT tissues, 4 coverslips (containing 8 brain slices total, from 3 mice) were successfully imaged and analyzed. For *Mecp2 +/-* mutant tissues, 10 coverslips (containing 20 brain slices total, from 5 mice) were imaged with RNA-MERFISH and MeCP2 immunostaining, and 4 out of these 10 coverslips (containing 8 brain slices total, from 4 mice) were further completed for DNA-MERFISH imaging and analysis. In this study, we did not gel-embed and clear the brain tissues, and hence the coverslips were not silanized for gel sticking (*50, 89*), but we washed and cleaned the coverslips with 70% ethanol before collecting tissue slices.

After tissue slices were collected onto coverslips, tissue slices were fixed by treating with 4% paraformaldehyde (PFA) (Electron Microscopy Sciences, #15714) in 1×PBS (Corning, 21-031-CV) for 10-12 minutes at room temperature, washed three times with 1×PBS and then stored in 70% v/v ethanol in water at 4 °C for at least 18 hours to permeabilize cell membranes. The tissue slices were either subsequently hybridized with MERFISH probes and imaged or were stored in 70% v/v ethanol at 4 °C for no longer than 2 months before they were hybridized with MERFISH probes and imaged.

#### Gene selection for RNA-MERFISH

We have previously identified cell types and mapped their spatial organization in the mouse MOp by imaging a panel of 258 genes, among which 242 genes were imaged using MERFISH and the remaining 16 genes were imaged with eight sequential rounds of two-color FISH (*50*). In this work, in order to distinguish transcriptionally distinct cell types in the mouse MOp, we used the same panel of the 242 genes for the RNA-MERFISH run, as in our previous study (*50*). Even without the 16 remaining genes previously imaged by sequential rounds of FISH, we were able to identify all subclasses of cells, as identified in our previous study (*50*).

#### Codebook and encoding probe design for RNA-MERFISH

Binary barcodes for the 242 genes were designed as previously described (*50*), which were drawn from a 22-bit, Hamming-Distance-4, Hamming-Weight-4 encoding scheme. The encoding probes for RNA-MERFISH were designed and obtained as described previously (*50*). In brief, each encoding probe contains a targeting sequence, which is complementary to a target region on a target RNA, and multiple readout sequences. We designed 22 readout sequences in total, each corresponding to one of the 22 bits. The collection of encoding probes targeting each gene contains a total of 4 readout sequences, corresponding to the 4 bits that read “1” in the barcode for that gene. Each encoding probe contains two out of the four readout sequences encoding that gene. Unlike our previous study, where we detect these readout sequences directly with dye-labeled readout probes complementary to the readout sequences (*50*), in this study, we detected these readout sequences by adaptor probes followed by readout probes, as described in the “RNA-MERFISH imaging” section below.

#### Genomic locus selection for DNA-MERFISH

We designed three different encoding probe libraries for DNA-MERFISH (and also sequential DNA-FISH) as described in the main text, targeting three groups of genomic loci, respectively. (1) The first encoding probe library targets 988 genomic loci that are approximately evenly distributed across the mouse genome with ∼2.5Mb of spacing (except the Y chromosome). (2) The second library targets 28 genomic loci that are centered around transcription start sites (TSSs) for representative marker genes for different MOp cell types. These 28 marker genes are *Slc30a3*, *Slc17a7*, *Slc32a1*, *Gad1*, *Otof*, Rspo1, *Pvalb*, *Sst*, *Vip*, *Sncg*, *Lamp5*, *Lratd2*, *Tshz2*, *Syt6*, *Nxph4*, *Cux2*, *Rorb*, *Sulf2*, *Ptpru*, *Car3*, *Aqp4*, *Flt1*, *Igf2*, *Pdgfra*, *Sox10*, *Ctss*, *Vtn*, and Bgn. These 28 marker genes were previously shown as marker genes for distinct MOp cell types (*50*). (3) The third library targets 965 genomic loci that are candidate cell-type-specific super-enhancer loci for different MOp cell types. These candidate cell-type-specific super-enhancer loci are selected based on published snATAC-seq data of the mouse MOp (*52*). Specifically, for each MOp cell type identified previously (*50, 52*), cell-type-specific pseudo-bulk ATAC peaks were called using MACS2 (*90*) and peaks that are within 12.5kb of each other were stitched together and considered a super-enhancer locus, and peaks within 2.5kb of a TSS were excluded. Then, cell type-specific candidate super-enhancers were called from these stitched peaks using the Rank Ordering of Super-Enhancers (ROSE) method (*55, 91*) with a fitted cutoff score for each cell type. Because candidate super-enhancer peaks in different cell types may correspond to genomic regions that overlap with each other, we iteratively merged all cell-type-specific super-enhancer peaks if the peaks identified in different cell types are within a 100kb interval. We calculated the average overlapping ratio of each merged candidate super-enhancer peak (defined as the average single peak size divided by the merged peak size). We selected the merged candidate super-enhancer peaks whose average overlapping ratio is >= 0.8, which in turn are either single peaks that are specific to a single cell type, or single peaks that are on average 80% shared among multiple cell types. Among these selected candidate super-enhancer peaks, those larger than 15kb were kept.

For each genomic locus described above, we designed encoding probes against a 20kb segment near or within the locus. Segments whose sequence allows for designing of ∼50 encoding probes were kept as the final target genomic loci in our DNA-MERFISH measurements. Together, a total of 1981 genomic loci that were imaged in this study.

#### Codebook design for DNA-MERFISH

Binary barcodes for (1) genomic-locus panel 1: the 988 genomic loci that are approximately evenly distributed across the mouse genome, and (2) genomic-locus panel 2: the 965 genomic loci that are cell-type-specific candidate super-enhancers were designed separately but in the same fashion as described below. For genomic-locus panel 1 we first generated all possible 99-bit binary barcodes with a Hamming weight of 3 (i.e., each barcode containing three “1” bits and 96 “0” bits) based on the covering design (https://www.dmgordon.org/cover/). Then we randomly selected 988 barcodes from this list iteratively for each chromosome to maximize the balance of on-bits over all bits and over all used barcodes within the certain chromosome (so that for each bit, we kept the barcoded loci reading “1” from the same chromosome as far from each other as possible) and maintained a Hamming distance of 4. In other words, this resulted in an approximately equal number of loci imaged per bit for each chromosome and balanced over all bits. Due to the polymeric nature of DNA, locus pairs on a given chromosome tend to localize physically together if their genomic distance is short. Therefore, we allowed loci within the same chromosome to exchange barcodes so as to optimize (i.e., maximize) the minimal genomic distance between loci with barcodes reading “1” at the same code position. When comparing code assignments with identical minimal genomic distances between loci with barcodes reading “1” at the same code position, we selected the code assignment that minimized the coefficient of variation of genomic distances between loci with barcodes reading “1” at the same code position (so that genomic distances between these loci have both larger means and smaller standard deviations). Binary barcodes for the genomic-locus panel 2 (the 965 cell-type-specific candidate super-enhancer loci) were chosen iteratively and similarly as above, except that they were from a list generated using all possible 95-bit binary barcodes.

For the genomic loci containing the TSS of the 28 marker genes (genomic-locus panel 3), we used two different imaging strategies. In the first strategy, the 28 loci were probed independently with sequential rounds of multicolor DNA-FISH, and this strategy was used for the wild-type mouse brains and the 11-week-old *Mecp*2 +/- female mouse brains. In the second strategy, to reduce the total imaging time, we imaged genomic-locus panel 3 together with genomic-locus panel 2 using a single MERFISH run. Specifically, 28 barcodes were selected from the 95-bit binary barcodes that were not used for encoding genomic-locus panel 2; we selected these barcodes randomly and iteratively to maximize Hamming distances of neighboring genomic loci, following the same design principle as described above for genomic-locus panels 1 and 2. This strategy was used for imaging the 27-week-old *Mecp*2 +/- female mouse brains.

#### Encoding probe design for DNA-MERFISH

Encoding probes for DNA-MERFISH (and also sequential DNA-FISH) were synthesized from a pool of oligonucleotides purchased from Twist Biosciences whose sequences are listed in Table S1. Each oligo in this pool consisted of the following sub-sequences (from 5’ to 3’):

1. A 20-nucleotide (nt) or 19-nt forward priming region for PCR amplification and reverse transcription (RT).
2. A 20-nt readout sequence.
3. A 42-nt target sequence, designed to bind uniquely to a single targeted genomic locus without containing repetitive sequences.
4. Additional two 20-nt readout sequences.
5. A 20-nt or 19-nt reverse priming sequence for PCR amplification.

The forward and reverse priming sequences were chosen from a previously generated list of random 20-nt sequences optimized for PCR, as described previously (*24*).

We designed 225 readout sequences (99 for MERFISH imaging of genomic-locus panel 1; 95 for MERFISH imaging of genomic-locus panel 2, and 28 for sequential multicolor FISH imaging of genomic locus panel 3) that have been validated to have a good observed signal-to-noise ratio (SNR) in total. In the case of DNA-MERFISH imaging, we assigned 3 different readout sequences (as sub-sequences #2 and #4 described above) for each genomic locus, corresponding to the 3 bits that read “1” in the barcode for that locus. In case of sequential multicolor FISH, we assigned one readout sequence for each color channel in each round, targeting one genomic locus (so that sub-sequences #2 and #4 described above have the same sequence). These readout sequences were chosen from a list of 30-nt sequences with minimal homology to the mouse (and human) genome, as described previously (*24, 92*). We detected these readout sequences by adaptor probes followed by readout probes, as described in the “DNA-MERFISH imaging” section below.

The 42-nt target sequence was chosen similarly to a procedure described previously (*24*). Briefly, we repeated the following procedure for each genomic region of interest. First, we created a list of all 42-nt sequences complementary to the genomic region of interest (starting at each possible base in the targeted region). Then, sequences were filtered to ensure a GC content of 40-60% and a melting temperature of 57-67 degrees Celsius. These sequences were further filtered using BLAST (*93*) by limiting the allowed degree of homology to the mouse genome, the mouse transcriptome, and a database containing repetitive sequences using the same procedure as previously (*24*). Finally, target sequences were selected from the remaining sequences after the final filtering step such that no genomic overlap exists between any pair of target sequences.

#### Encoding probe synthesis

Encoding probes were amplified from the template library described above (see the “Codebook and encoding probe design for RNA-MERFISH” section and “Encoding probe design for DNA-MERFISH” section above). This was done using a previously described amplification protocol (*24, 47*) with additional modifications and involved the following steps. (1) The initial oligo pool was amplified using limited-cycle PCR for approximately 12 cycles. PCR primer sequences are listed in Table S3. The reverse primer used in this step also introduced a T7 promoter sequence via primer extension. (2) The resulting PCR product was purified via column purification (DNA Clean & Concentrator Kit, Zymo Research, D4033). (3) The purified PCR product underwent further amplification and conversion to RNA by a high-yield *in-vitro* T7-mediated transcription reaction (HiScribe T7 polymerase kit, NEB, E2050). (4) The resulting RNA product was purified via column purification (Monarch® RNA Cleanup Kit, NEB, T2050). (5) The purified RNA product was converted back to single-stranded DNA (ssDNA) by a reverse transcription reaction (Maxima H Minus Reverse Transcriptase, ThermoFisher, EP0753) using the forward primer. For the DNA-MERFISH encoding probe library sets, a single deoxyuracil residue (dU) was optionally introduced to replace a thymine (T) in between 6-14 base where applicable in the forward primer for subsequent USER enzyme cleavage (NEB, M5505) that will be described below. (6) The ssDNA product was subjected to alkaline hydrolysis to remove residual RNA and was subsequently column purified by ssDNA purification procedures (DNA Clean & Concentrator Kit with Oligo Binding Buffer, Zymo Research, D4032 and D4060). If dU was introduced in the PCR forward primer, the ssDNA product was additionally cleaved by USER enzyme (NEB, M5505) before alkaline hydrolysis. (7) The purified ssDNA product was dried in vacuum and resuspended in water to achieve the desired concentration of primary probe. All primers were purchased from Integrated DNA Technologies (IDT).

#### Readout and adaptor probe preparation

All readout and adaptor probes were ordered from IDT (see Tables S2 and S3) and were diluted directly from stock.

#### Overview of experimental system of integrated RNA-MERFISH and DNA-MERFISH

Experimental steps described below were conducted using a home-built imaging platform, including steps described in the “RNA-MERFISH imaging” section, “Sample positioning alignment preparation after RNA-MERFISH” section, “Sample positioning alignment for DNA-MERFISH” section, and “DNA-MERFISH imaging” section.

The physical setup of the home-built imaging platform consists of several components. A custom-built fluorescence microscope was used to acquire images, and a custom-built fluidics system was used to automatically perform buffer exchanges on the microscope stage. Custom software was used to synchronize and control the various microscope and fluidic components, and to automate many experimental steps (https://github.com/ZhuangLab/storm-control). Below is a detailed description of each of these components.

#### Microscope setup for Image acquisition

Image acquisition was performed using a custom-built microscope system, as previously described (*24*) but with some modifications. The system was built around a Nikon Ti-U microscope body with a Nikon CFI Plan Apo Lambda 60x oil immersion objective with 1.4 NA. The system also included a Nikon 10x objective for quick identification of the sample position under the microscope. Illumination of samples was through a Lumencor CELESTA light engine (a fiber-coupled solid-state laser-based illumination system) with the following wavelengths: 405 nm, 477 nm, 546 nm, 638 nm, and 749 nm. This system was used with a penta-bandpass dichroic (IDEX, FF421/491/567/659/776-Di01-25×36) and a penta-bandpass filter (IDEX, FF01-441/511/593/684/817-25). A scientific CMOS camera (Hamamatsu FLASH4.0 or Hamamatsu C13440 with factory calibration for single-molecule imaging) was used for image acquisition.

Each camera field of view (FOV) consisted of 2048 x 2048 pixels, with a camera pixel corresponding to 108 nm in the X and Y dimensions in the imaging plane for the 60x oil immersion objective with 1.4 NA. Sample position in three dimensions was controlled using a XYZ stage (Ludl Electronic Products). A custom-built auto-focus system (*24*) was used to maintain a constant focal plane over prolonged periods of time. This was achieved by comparing the relative position of two IR laser (Thorlabs, LP980-SF15) beams reflected from the glass-fluid interface and imaged on a separate CMOS camera (Thorlabs, uc480).

These different components were controlled using a National Instruments Data Acquisition card (NI PCIe-6321, X Series DAQ) and custom software (see the “Software for controlling experimental components” section below).

#### Fluidics system configuration

The fluidics system consisted of several main components: a pump, a set of valves connected in series, a flow chamber in which the sample was mounted, and tubing and connectors. We used a peristaltic pump (Gilson, MINIPLUS 3) to generate flow in the system. The pump was connected to an array of 12-way valves (IDEX, EZ1213-820-4), connected in series. In this study, we used three 12-way valves connected in the following manner. Each valve’s last connection was used as the input of the next valve in the series (except for the last one), while the rest were connected to a tube containing the buffer for a single round of hybridization. A fixed subset of the valves was used for imaging, bleaching and wash buffers. This valve system was used to flow the various buffers into the FCS2 flow chamber (Bioptechs, 060319-2), in which the sample was placed together with a 0.5-mm-thick flow gasket (Bioptechs, DIE# F18524). The chamber output was connected to a waste collection vessel, forming an open flow system. Components were connected using elastic plastic tubing, and connections were additionally sealed using a pressure adessive (Blu-tack). Put together, these components allow control of both the rate of fluid flow and of the type of fluid flowing at any given time.

The system was controlled using custom software (see the “Software for controlling experimental components” section below). Overall, this system allowed for 24 rounds of hybridization (depending on the number of valves and the number of spots reserved for special buffers). We constructed a bypassing flow circuit by a tubing connecting two 3-way adaptor sets (Bioptechs, 162003-1) that were added to each side of the FCS2 flow chamber (Bioptechs, 060319-2). Therefore, in experiments where the number of hybridization rounds exceeded the capacity of the flow system, we replaced the buffers via the following procedures: (1) We turned the manual valves on the 3-way adaptor sets to short-connect the sample-containing chamber. (2) All valves were washed using 30% v/v formamide (ThermoFisher, AM9342) in water and then with double-distilled water. (3) The new set of buffers was introduced, and the chamber was reconnected to the flow system to allow for the next round of hybridization.

#### Software for controlling experimental components

All system components were controlled using custom-built software as described previously (*24*), available at https://github.com/ZhuangLab/storm-control. This software package is composed of several main modules that work in concert:

“Hal” is the software package used to control and synchronize all illumination and microscope components. We note that in some cases it is necessary to write drivers for components which are not included in this package. Hal is also used to define imaging parameters, such as illumination light intensity, sequence of stage and illumination operations during imaging (e.g., during a z-scan), exposure time etc.

“Steve” is a module used to take mosaic images (i.e., a composite image that is made up of many individual fields of view (FOV)) of the sample and to select FOV for imaging in experiments.

“Kilroy” is the software used to control the fluidics components, and to define pre-programmed sequences of operations to be performed as sets (e.g., the set of operations that happens when a new round of hybridization is performed).

“Dave” can issue commands to both Hal and Kilroy and is used to automate the performance of data collection by defining in advance a complete set of fluidics system and microscope operations, the order and time-lag in which they are to be performed.

The general flow of an experiment is that, before the experiment starts, Hal and Kilroy are loaded with the parameters and specifications to be used. After the sample is loaded and the chamber is filled with the imaging buffer, a mosaic image of the DAPI channel is taken using Steve, and FOVs of interest are selected. A file is then generated to specify the sequence of operations throughout the entire experiment and is loaded to Dave, together with the coordinates of the selected regions of interest. The rest of the experiment is run automatically, without manual intervention. If the number of rounds in the experiment exceeds the capacity of the flow system, the automatic sequence specifies actions up to the capacity of the system. The buffers are then replaced, a new Dave file is created, and this is repeated until all rounds of imaging are completed.

#### Encoding-probe hybridization for RNA-MERFISH

After tissue preparation (as described in the “Tissue preparation for integrated RNA-MERFISH and DNA-MERFISH” section above), the prepared tissue slices were hybridized with the RNA-MERFISH encoding probe set as previously described (*50*), except for that tissue slices were not gel-embedded and cleared. Briefly, the samples were removed from the 70% v/v ethanol and washed with 2×saline sodium citrate (SSC) (ThermoFisher, AM9763) three times. Then, tissue slices were illuminated by multi-band light emitting diode arrays for three hours to reduce the autofluorescence background (*49*), which is critical in consideration of the omission of the tissue clearing step. Next, we equilibrated the samples with the wash buffer, containing 2×SSC and 30% v/v formamide) (ThermoFisher, AM9342), for 30 min at room temperature. The wash buffer was then aspirated from a coverslip, and the coverslip was inverted onto a 50-μl droplet of the encoding-probe mixture on a parafilm-coated Petri dish. The encoding-probe mixture comprised approximately 1 nM of each encoding probe for RNA-MERFISH, and 1 μM of a polyA-anchor probe (IDT) in 2×SSC with 30% v/v formamide, 0.1% wt/v yeast tRNA (ThermoFisher, 15401011) and 10% v/v dextran sulfate (Sigma, D8906). We then incubated the sample at 37 °C for 36–48 h. The polyA-anchor probe contained a mixture of DNA and LNA nucleotides (synthesized from IDT) (/5Acryd/TTGAGTGGATGGAGTGTAATT+TT+TT+TT+TT+TT+TT+TT+TT+TT+T, where T+ is locked nucleic acid, and /5Acryd/ is 5’ acrydite modification), which can hybridize to the polyA sequence on the polyadenylated mRNAs, as previously described (*50*). The polyA-anchor probe was mainly used for MERFISH detection of the cellular total RNAs. After hybridization, the samples were washed with the wash buffer, containing 2×SSC and 30% v/v formamide, for 30 min at 47 °C for a total of two times to remove excess encoding probes and polyA-anchor probes. The coverslips were then washed three times with 2×SSC and were post-fixed with 4% PFA in 2×SSC for 10 minutes at room temperature. The coverslips were then washed three times with 2×SSC and were incubated with yellow-green fiducial beads (FluoSpheres™ Carboxylate-Modified Microspheres, ThermoFisher, F8803) in 2×SSC for 10 minutes at room temperature. Lastly, the coverslips were washed once with 2×SSC and then stored at 4 °C in 2×SSC supplemented with 1:100 murine RNase inhibitor (NEB, M0314S) for no more than two days, before subsequent RNA-MERFISH imaging.

#### RNA-MERFISH imaging

After encoding-probe hybridization of samples, we assembled the sample into the FCS2 flow chamber (Bioptechs, 060319-2) and performed RNA-MERFISH imaging of the target genes using the home-built imaging platform, as described in the “Overview of experimental system of integrated RNA-MERFISH and DNA-MERFISH” section (*24*). All fluid exchanges involved in this part of the protocol were performed using a custom-built fluidics system, as described in the “Fluidics system configuration” section. For each coverslip, typically 100-300 FOVs from two coronal brain slices were selected for MERFISH imaging. Nucleus staining by DAPI was used to help select FOVs of interest.

During the MERFISH imaging course, we performed multiple rounds of hybridization and imaging, each round reading out two bits in the barcode. For each round, the samples were imaged as follows unless otherwise described. (1) The sample was hybridized with a set of oligonucleotide probes termed “adaptor probes” that converts bit-specific readout sequences on the encoding probes to common readout sequences. For example, for a MERFISH experiment with 22-bit Hamming-weight 4 barcodes, we designed 22 readout sequences, each of which corresponds to one of 22 bits. The collection of encoding probes targeting each gene would contain a total of 4 readout sequences, corresponding to the 4 bits that read “1” in the barcode for that gene. We then designed 22 adaptor probes, each comprising two parts: one part having a sequence that is complementary to one of the bit-specific readout sequences and the other part having two copies of a common readout sequence. We designed two common readout sequences in total, each corresponding to one imaging color channel. During each round, we added two different adaptor probes, corresponding to two bits, converting the two corresponding readout sequences on the encoding probes to the two common readout sequences. The adaptors probes were added at 100 nM in the adaptor hybridization buffer containing 2×SSC, 30% v/v formamide (ThermoFisher, AM9342), for 10 minutes at room temperature. (2) The sample was washed with the wash buffer containing 2×SSC and 30% v/v formamide (ThermoFisher, AM9342), for 3 minutes at room temperature. (3) The sample was then hybridized with a set of fluorescently labeled oligonucleotide probes termed “common readout probes” (synthesized by IDT), each common readout probe with a sequence that is complementary one of the common readout sequences on the adaptor probes. The fluorescence dyes were linked to the common readout probes via a disulfide bond. The common readout probes were added at 20 nM in the readout hybridization buffer containing 2×SSC, 30% v/v formamide (ThermoFisher, AM9342) and 0.005% v/v Triton-X (Sigma, T8787), for 15 minutes at room temperature. (4) The sample was washed again with the wash buffer containing 2×SSC and 30% v/v formamide (ThermoFisher, AM9342), for 3 minutes at room temperature. (5) The sample was then incubated and kept in imaging buffer comprising 5 mM 3,4-dihydroxybenzoic acid (Sigma, P5630), >50 µM trolox quinone (generated by UV radiation of a trolox solution; Sigma, 238813), 1:500 recombinant protocatechuate 3,4-dioxygenase (rPCO; OYC Americas, 46852904), 1:500 Murine RNase inhibitor (NEB, M0314L), 10 mM Tris-HCl (ThermoFisher, 15568025), and 5 mM NaOH (to adjust pH to 8.0; Sigma, S2770) in 2×SSC. (6) Finally, the samples were imaged using a 60x oil immersion objective, and signals for the two common readout probes and fiducial beads were collected in the 750-nm, 650-nm, and 488-nm channels, respectively at a rate of ∼5 Hz. All channels were imaged for 13 consecutive 1-µm-thick z-stacks for each FOV of interest as described above.

All rounds of imaging were carried out as described above except for the first and the last round. For the first round, before the imaging buffer step (5), the sample was additionally washed in 2×SSC once and then incubated in 2×SSC containing 1 µg/mL DAPI (ThermoFisher, D1306) for 10 minutes to stain nuclei. Signal of DAPI was additionally collected in 405-nm channel, and either 405-nm channel or all channels in this round were imaged for 50 consecutive 0.25-µm-thick z-stacks for each FOV of interest. For the last round, adaptor probes targeting the polyA-anchor probes were added at 10 nM to stain the cellular total RNAs. Signal of polyA-anchor probes was collected in 750-nm or 650-nm channel, and it was imaged for either 13 or 50 consecutive 1-µm-thick z-stacks for each FOV of interest.

Between each imaging round, the signal from the previous round was extinguished. This was achieved by incubating the sample in the cleavage buffer comprising 2×SSC, 30% formamide and 50 mM Tris (2-carboxyethyl) phosphine (TCEP; Sigma, 646547) for 10 minute at room temperature, which cleaved the disulfide bond connecting fluorophores to readout probes. The cleavage buffer also contained 167 nM unlabeled common readout probes, which blocked unoccupied common readout sequences on the adaptor probes from interfering with the next round of hybridization. After incubation with the cleavage buffer, the sample was washed with the hybridization wash buffer containing 2×SSC and 30% v/v formamide (ThermoFisher, AM9342), for 3 minutes at room temperature, before the next round of hybridization of adaptor probes. The same cleavage buffer incubation and wash were also employed after the last round of RNA-MERFISH imaging to remove any remaining readout signals from RNA-MERFISH.

#### Immunofluorescence imaging for MeCP2

For imaging tissue sections from *Mecp2 +/-* mutant female mice, the samples were additionally stained and imaged for antibodies against MeCP2 after RNA-MERFISH. The following procedures for MeCP2 antibody staining and imaging were performed using the home-built platform as described above for RNA-MERFISH. Specifically, (1) the sample was washed twice with 1×PBS (Corning, 21-031-CV) after RNA-MERFISH. (2) The sample was then incubated in the blocking buffer containing 1×PBS, 5% m/v bovine serum albumin (BSA; Jackson ImmunoResearch, 001-000-162), and 0.5% v/v Triton-X (Sigma, T8787), for 30 minutes at room temperature. (3) The sample was incubated with primary antibody against MeCP2 (rabbit anti-Mecp2, Cell Signaling, #3456), 1:300 diluted in the blocking buffer described above, for 3-4 hours at room temperature. (4) The sample was then washed with 1×PBS three times, for 5 minutes each. (5) The sample was incubated with a fluorescently tagged secondary antibody (goat anti-rabbit conjugated with Alexa-647, Invitrogen, #A-21245), 1:300 diluted in the blocking buffer described above, for 1 hour at room temperature. (6) The sample was then washed again with 1×PBS three times, for 5 minutes each. (7) The sample was incubated and kept in the imaging buffer as described above in the “RNA-MERFISH imaging” section above. (8) Finally, the samples were imaged using a 60x oil immersion objective, and signals for the secondary antibody and fiducial beads were collected in the 647-nm and 488-nm channels, respectively at a rate of ∼5 Hz. All channels were imaged for 13 consecutive 1-µm-thick z-stacks for each FOV of interest as described above.

#### Sample positioning alignment preparation after RNA-MERFISH

In order to align the sample on the home-built imaging platform for both RNA-MERFISH and DNA-MERFISH, we took DAPI images of 10 FOVs under the 10x objective, either before or after RNA-MERFISH imaging. These 10 FOVs were selected from two coronal slices for each imaged coverslip in an approximately evenly distributed manner, and they were used as landmarks for aligning the sample orientation and positioning between RNA-MERFISH and DNA-MERFISH, which will be described below in the “Sample positioning alignment for DNA-MERFISH**’’** section below.

#### Sample wash after RNA-MERFISH

After RNA-MERFISH (and immunofluorescence staining for *Mecp2* +/- mutant tissues when appropriate) and sample positioning alignment preparation, the sample was removed from the microscope. Next, the sample was washed with a harsh wash buffer containing 2×SSC and 60% v/v formamide (ThermoFisher, AM9342), for 30 minutes at 46 °C to remove bound probes from RNA-MERFISH. The sample was then washed with the hybridization wash buffer containing 2×SSC and 30% v/v formamide (ThermoFisher, AM9342) briefly. The sample was then washed three times with 2×SSC and was post-fixed with 4% PFA in 2×SSC for 10 minutes at room temperature. The sample was then washed three times with 2×SSC and then stored in 70% ethanol at 4 °C for at least 18 hours before subsequent DNA-MERFISH.

#### Encoding-probe hybridization for DNA-MERFISH

After RNA-MERFISH and sample wash, we typically stored the sample in 70% ethanol before DNA-MERFISH imaging. For DNA-MERFISH, the sample was removed from the 70% ethanol and washed with 2×SSC three times. Then, the sample was illuminated by the multi-band light emitting diode arrays for three hours to reduce the autofluorescence background, as what was done before RNA-MERFISH imaging (*49*). Next, the sample was then post-fixed with 4% PFA in 2×SSC for 10 minutes at room temperature. This post-fixation is important to preserve the nucleus structure during the following steps including heat denaturation.

After post-fixation, the sample washed with 1×PBS (Corning, 21-031-CV) three times and was incubated in a 1×PBS solution containing 0.1% of sodium borohydride (Sigma, 452882) to reduce autofluorescence background. Then, the sample was treated with 0.5% v/v Triton-X (Sigma, T8787) in 1×PBS for 10 minutes at room temperature, followed by three times of 1×PBS wash. Samples were treated with 0.1 M hydrochloric acid (HCl; Sigma, H9892) for 5 minutes at room temperature to increase the target DNA accessibility, presumably by depurinating DNA and denaturing histones, followed by washes in 1×PBS 2-3 times. Samples were then treated with a solution of 0.1 mg/mL RNase A (ThermoFisher, EN0531) dissolved in 1×PBS for 30-45 minutes at 37 ℃, to remove potential sources of off-target binding to RNA. Following this treatment, cells were incubated in pre-hybridization buffer, consisting of 2×SSC (ThermoFisher, AM9763) and 50% formamide (ThermoFisher, AM9342) for approximately 30 minutes. Next, the cell coverslip was inverted and placed on a drop of 50 μL of hybridization buffer (2×SSC, 50% v/v formamide, ThermoFisher, AM9342, 10% dextran sulfate, Sigma, D8906) containing a mixture of encoding probes at ∼1 nM of each encoding probe for the DNA-MERFISH run, with 5 µg mouse Cot-1 DNA (ThermoFisher, 18440016) in a 60-mm petri dish. The dish was partially submerged in a water bath at ∼86 ℃ for 3 minutes and incubated at 47 ℃ in a humidified chamber for 36-48 h. After incubation with encoding probes, the sample was washed in 2×SSC and 30% v/v formamide (ThermoFisher, AM9342), for 30 minutes at 47 °C for a total of two times to remove excess encoding probes. The coverslips were then washed three times with 2×SSC and were post-fixed with 4% PFA in 2×SSC for 10 minutes at room temperature. The coverslips were then washed three times with 2×SSC and were incubated with yellow-green fiducial beads (FluoSpheres™ Carboxylate-Modified Microspheres, ThermoFisher, F8803) in 2×SSC for 10 minutes at room temperature. Lastly, the coverslips were washed once with 2×SSC and then stored at 4 °C in 2×SSC for no more than two days, before subsequent DNA-MERFISH imaging.

#### Sample positioning alignment for DNA-MERFISH

After the encoding-probe hybridization step, we assembled the sample into the FCS2 flow chamber (Bioptechs, 060319-2) in a similar manner as for RNA-MERFISH so that the relative orientations of the sample to the microscope stage coordinate system were roughly the same for both RNA-MERFISH and DNA-MERFISH by visual comparison. Next, we incubated the sample with 2xSSC containing 1 µg/mL DAPI (ThermoFisher, D1306) for 10 minutes to stain the cell nuclei and we used the 10x objective to locate the 10 landmarks that we selected in the “Sample positioning alignment preparation after RNA-MERFISH” section. Based on these landmarks and also the general morphological features of brain slices, we further manually rotated the flow chamber to match the relative angle of the sample to the microscope stage coordinate system by visual comparison. Then, we calculated the rigid XY rotation and translation matrix from the original centroid coordinates of these 10 selected FOVs (recorded during the step described in the “Sample positioning alignment preparation after RNA-MERFISH” section) to their new centroid coordinates (recorded during this step) by singular-value decomposition (SVD), acknowledging the fact that the tissue underwent affine transformation between RNA-MERFISH and DNA-MERFISH imaging. We repeated the above flow chamber rotation step until the calculated rotation angle was as close as possible to 0 degree. We kept the last calculated rigid XY rotation and translation matrix and used it to locate the same FOVs where we acquired images for RNA-MERFISH. Lastly, we took DAPI images of 4-5 FOVs among these FOVs using the 60x objective. We calculated the average XY shift between the DAPI images from RNA-MERFISH and DAPI images for these 4-5 FOVs and offset the reference point (0, 0) in our microscope setting by the calculated average XY shift.

#### DNA-MERFISH imaging

After locating the FOVs of interest where we acquired images for RNA-MERFISH, we performed DNA-MERFISH imaging for all targeted genomic loci in a similar manner as described in the “RNA-MERFISH imaging” section and also described previously (*24*). All fluid exchanges involved in this part of the protocol were similarly performed using a custom-built fluidics system, as described in detail in the “Fluidics system configuration” section.

As in RNA-MERFISH imaging, we perform multiple rounds of hybridization and imaging for DNA-MERFISH, each round reading out three bits in the barcode. For each round, the samples were imaged in the same manner as described in the “RNA-MERFISH imaging” section except for the following differences. (1) During the adaptor-probe hybridization step, the adaptor hybridization buffer contained 35% v/v formamide (ThermoFisher, AM9342) for DNA-MERFISH instead of 30% v/v formamide for RNA-MERFISH. When the sample was incubating with adaptor probes, we also performed a photobleaching step by illuminating each FOV with the maximum available light power of the 560-nm, 647-nm, and 750-nm wavelengths together for 3-5 seconds. As a result, the incubation time duration varied from 15 to 30 minutes depending on the number of FOVs to be photobleached. (2) During the imaging step, the imaging buffer did not need to include Murine RNase inhibitor. (3) Instead of imaging two common readout probes in two color channels per round as in RNA-MERFISH, we imaged three common readout probes in three color channels per round in DNA-MERFISH. The sample was imaged using the 60x oil immersion objective, and signals for the 3 common readout probes and fiducial beads were collected in the 750-nm, 650-nm, 561-nm, and 488-nm channels, respectively, at a rate of ∼10 Hz. (Note that for the first round, where DAPI was additionally stained, signal of DAPI was collected in the 405-nm channel, as what was done for RNA-MERFISH imaging.) (4) All channels, including 405-nm if used, were imaged for 50 consecutive 0.25-µm-thick z-stacks for each FOV of interest.

In addition to these differences above regarding each sequential round, there were also additional modifications for DNA-MERFISH imaging due to the relatively large number of imaging rounds compared to RNA-MERFISH. Because our fluidic system allowed loading of a maximum of 24 different hybridization solutions, we washed our fluidic system and replaced hybridization solutions when the number of hybridization rounds exceeded this capacity of the fluidic system. The fluidic system wash and buffer replacement procedures were described in the “Fluidics system configuration” section. For long-duration sample imaging, we also performed a gentle post-fixation step with 2% PFA in 2×SSC for 5 minutes periodically between hybridization and imaging rounds (every ∼3-4 days) to maintain the structural integrity of our sample (*94*).

For each DNA-MERFISH experiment, we typically first imaged 988 genomic loci that are uniformed spaced along the genomic coordinate (genomic-locus panel 1) over ∼35 rounds of imaging, and then imaged 965 genomic loci of the super-enhancers (genomic-locus panel 2) plus, in some cases, the TSSs of the 28 marker genes (genomic-locus panel 3) over ∼35 rounds of imaging.

#### Sequential DNA-FISH imaging

The TSSs of the 28 marker genes (genomic-locus panel 3) were either imaged by DNA-MERFISH together with the 965 genomic loci of cell-type-specific super enhancers (genomic-locus panel 2) as described above or were imaged separately by sequential round of three-color FISH. In the latter case, the encoding probes for genomic-locus panel 3 were co-hybridized with the DNA-MERFISH probes for genomic-locus panels 1 and 2, also at ∼1 nM of each encoding probe. Sequential rounds of adaptor probe and common readout probes were then hybridized and imaged, in the same manner as described in the “DNA-MERFISH imaging” section, except that three genes, instead of three bits of the barcodes, were imaged in each round.

#### Overview of analysis pipeline for integrated RNA-MERFISH and DNA-MERFISH

The image analysis pipeline for processing and decoding MERFISH spots was implemented in Python, and the code is available at: https://github.com/ZhuangLab/Chromatin_Analysis_2023. The overall pipeline consists of the following steps:

1. Image corrections including bleed-through correction, chromatic-aberration correction, illumination-intensity uniformity correction.
2. Identify and segment all imaged cell nuclei from RNA-MERFISH imaging.
3. For RNA-MERFISH, follow standard pixel-based decoding to identify RNA molecules as described previously (*95*).
4. Translate the cell-nucleus segmentation from RNA-MERFISH images to DNA-MERFISH images.
5. Fit all detected DNA-MERFISH signals in imaging channels and determine corrected 3D coordinates based on drift correction.
6. Decode and assign identities to genomic loci and RNA molecules using custom algorithms and software.

#### Image corrections for color-channel bleed-through, chromatic aberration, and non-uniform illumination intensities

Our MERFISH microscope setup used a penta-band dichroic mirror and emission filter set, and we observed bleed-through between signals collected from different wavelength channels (i.e., for different fluorophores). To correct for this bleed-through, we devised a bleed-through correction approach by staining a set of samples with one single type of fluorophores whose emission corresponded to one of our microscope channels, and subsequently measuring the signal bleed-through in other channels. Given this bleed-through is approximately a fixed linear profile, we linearly decomposed the signal intensities from different channels at each pixel to correct for bleed-through between channels.

Because DNA-MERFISH aims to determine the precise XYZ coordinates (∼ 50-100 nm precision) of the genomic loci of interest, we also performed correction of chromatic aberration among different imaging channels. Bleed-through and chromatic aberration corrections for multi-color DNA-MERFISH imaging were performed by labeling the same set of genomic loci in each different imaging channel independently and by next comparing the location of signals of the same loci in the different color channels, respectively, as described previously (*24*). This shift in X and Y axes between each pair of channels is linearly correlated with X and Y positions in the FOV, and in Z axis is a constant (*96*). Therefore, we performed a linear regression of this shift on X, Y and Z axes for each pair of channels to obtain a shift-correction function. Next, we either translated signals of all pixels from an acquired image to generate a corrected image using this shift-correction function, or we kept this shift-correction function and applied it to the 3D fitted coordinates (generated by the “Spot fitting for DNA-MERFISH and DNA-FISH imaging” section below).

To better compare the signal intensities of foci across each whole FOV and facilitate better segmentation of cells on the edge of FOV, we also applied an illumination-intensity uniformity correction for each FOV. To do this, we generated the averaged laser intensity profile for each channel, and we next divided the acquired images by these fixed averaged laser intensity profiles to correct for the illumination differences within each FOV.

#### Cell and cell-nucleus segmentation for RNA-MERFISH

For the MOp datasets from wild-type mice, images for DAPI representing the cell nuclei and PolyT anchoring probe representing the cytoplasm from RNA-MERFISH were used to identify individual nuclei, estimate nucleus boundaries, and estimate the nuclear volume. Specifically, images were first downsampled on the x, y planes from 2048×2048 pixels to 1024×1024 pixels. 13 z-stacks with 1 μm-step were used for 3D segmentation as follows. We employed a deep-learning-based cell segmentation algorithm Cellpose 2.0 with a pre-trained model “TissueNet2” (*97*), which effectively identified the cell nuclei in 3D. We used anisotropy value of 1000/216 and diameter of 30 in Cellpose 2.0 model evaluation. We then used the segmentation masks determined using Cellpose 2.0 as the seed and performed watershed over the polyT image to determine cytoplasmic segmentation.

For the datasets from *Mecp2 +/-* mutant mice, images for DAPI from RNA-MERFISH were used for segmentation. Images were first downsampled on the x, y planes from 2048×2048 pixels to 1024×1024 pixels. 50 z-stacks with 250nm-step were used for 3D segmentation as follows. We employed a deep-learning-based cell segmentation algorithm Cellpose 2.0 with a pre-trained model “nuclei” (*97*). We used anisotropy value of 250/216 and diameter of 30 in Cellpose 2.0 model evaluation.

#### Decoding algorithm for RNA-MERFISH

RNA-MERFISH datasets were decoded by MERlin as previously described (*95*). Briefly, for each z plane, we aligned the images of different MERFISH imaging bits and generated overall normalization factors to obtain a normalized signal intensity vector for each pixel across all MERFISH bits. Based on this normalized intensity vector, we assigned the RNA identity for each pixel based on the MERFISH codebook we designed. Then, adjacent pixels that were assigned with the same barcodes were aggregated into putative RNA molecules, and then the list of putative RNA molecules was filtered to enrich for correctly identified transcripts as described previously (*50*), for a gross barcode misidentification rate at 5% using MERlin (*50*). All decoded RNA molecules were then assigned to each cell/nuclei segmentation based on their position relative to the segmentation mask (see the “Cell and cell-nucleus segmentation for RNA-MERFISH” section above), giving a cell-by-gene count matrix for the 242 genes measured by MERFISH. This cell-by-gene count matrix was next used for relevant analyses in the sections below.

#### Cell-nucleus segmentation transformation from RNA-MERFISH to DNA-MERFISH images

To obtain the correspondence of cell-nucleus segmentation between RNA-MERFISH and DNA-MERFISH images, we used the rigid XY rotation and translation matrix calculated in the “Sample positioning alignment for DNA-MERFISH” section to transform the cell-nucleus segmentations calculated from DNA-MERFISH images to align to those calculated from RNA-MERFISH images. We also used this rigid XY rotation and translation matrix to transform the cell-nucleus images from DNA-MERFISH and aligned their z-position using the fiducial beads. Then, we calculated the shift between the transformed DNA-MERFISH cell-nucleus image and RNA-MERFISH cell-nucleus image. Specifically, we split each 3D stack of 2D images into 8 equal 3D volumes and calculated the 2D image cross-correlation for each 3D volume. We removed the outlier correlation value and obtained the median correlation as the final correlation. By minimizing the sum of square difference between the transformed cell-nucleus images from DNA-MERFISH and the experimentally measured cell-nucleus images from RNA-MERFISH, we obtained the corresponding cell-nucleus segmentation for DNA-MERFISH and use it for downstream analyses.

#### Spot fitting for DNA-MERFISH and DNA-FISH imaging

The following analysis pipeline was applied to each imaged FOV in order to obtain the 3D positions of all genomic loci of interest. For each acquired DNA image, diffraction-limited spots within each identified nucleus were fitted to a 3D Gaussian function to identify their center of mass and brightness above local background (*24*). To make analysis more manageable, we fixed the number of fitted spots per image, which would be retained for decoding, to 20,000 or fewer in the DNA-MERFISH images (>3-fold greater than the number of distinct loci expected without noise). For sequential DNA-FISH images of the 28 marker gene TSS loci, we fixed the number of fitted spots per chromosome per image to 3 or fewer.

The fitted spots were then used for identifying genomic loci and for determining their positions, as described in the “Decoding of fitted DNA spots” section below.

#### Drift correction

Fluidic exchange can cause sample position drift during MERFISH hybridization and imaging. To correct for this drift between hybridization rounds, we aligned images of fiducial beads acquired from different hybridization rounds by calculating cross correlations (using Skimage package function, “skimage.registration.phase_cross_correlation”). In DNA-MERFISH, the bead images were subsampled (usually as ¼ of x and y dimension of 3D image) and the 3D drifts were calculated iteratively until drifts calculated from 3 subsampled image pairs were within 1 pixel difference. The mean drift of these 3 selected drift values was then used for the drift correction of the 3D coordinates of each DNA spot between hybridization rounds.

#### Decoding of fitted DNA spots

After DNA spot fitting and drift corrections, fitted DNA spots were decoded according to their barcode assignment, as designed in “Codebook design for DNA-MERFISH” section above. Specifically, a list was generated for the drift- and aberration-corrected locations of all identified DNA spots in each bit-image (corresponding to a specific color channel in a specific round of imaging). For each detected spot DNA in every bit-image, we found all spots from other bit-images that were within a set cutoff distance (300 nm in x, y, and z) from the location of the given spot in the given bit image to from spot pairs across bits. To increase the recovery efficiency of all detected spots, we retained all spot pairs for further analysis, regardless of whether such spot pairs had other spot(s) from other bit-images that were within the set cutoff distance. Moreover, each spot pair was given a score for validity based on their 3D distances, intensity levels, and intensity variations. A spot pair, if valid within our codebook design, will have a higher score for validity if they have closer 3D distances between them, higher intensities, and low intensity variations between them. We then iterated through all spot pairs in a descending order of the validity scores. For each spot pair, we iteratively searched all possible spot trios within the cut-off distance (300 nm) containing the examined pair. Like for spot pairs, we then calculated the validity score for each spot trio that corresponds to a genomic locus in our codebook. We selected the spot trio with the highest score and excluded this trio from further analysis. After the above steps, for the remaining spot pairs without an identified trio, we next assigned barcode identities to all such spot pairs (by allowing for 1-bit error correction), generating a table of all candidate DNA spots according to our barcode design scheme. All candidate DNA spots were then assigned to each segmented cell nucleus based on their 3D positions.

After barcode identity and cell-nucleus assignment, we used the previously published spatial genome aligner algorithm, “Jie” (*57*), to pick the spots for different chromosome homologs without assuming the number of chromosome homologs for each nucleus. This is because some cell nuclei may be incompletely included in the 10-µm-thick coronal sections and contain an uncertain number of homologs. Briefly, the “Jie” algorithm utilizes the Gaussian chain polymer model to estimate the probability of observing a physical distance between a locus pair given their genomic distance, and the DNA polymer fiber connecting decoded spots with the maximal probability is picked. We introduced several modifications to the “Jie” algorithm to best fit our library design and DNA imaging experiment. (1) We adjusted the threshold score calculation as follows. We considered a case where the physical distance between the pair of any two adjacent imaged loci was set as the σ value in the Gaussian chain polymer model.

Then, we calculated the probability of observing such a polymer fiber and pick only DNA fibers with higher probability. (2) Moreover, we adjusted the penalty scores in Jie and included a larger penalty for potential fiber path that is more than 1.5 μm because the original penalty scores sometimes assigned DNA spots from two distant fibers to one chromosome homolog. (3) We also explored several sets of parameters in Jie to pick more candidate spots while keeping the recovered chromosome fibers as continuous as possible. These parameters led to 0 – 2 recovered chromosome homologs for each chromosome per nucleus for most of the nuclei, as expected. (4) After the spot picking by Jie, we added an additional spot swapping algorithm to swap spots between the detected chromosome homologs within each nucleus as follows. We calculated the physical distance of each spot to their neighboring regions’ center of mass from each chromosome homolog and then progressively swapped all spots between the chromosome homolog groups to minimize their summed physical distance to their neighboring regions’ center of mass. The selected spots were then used to determine the 3D positions of the targeted genomic loci, which were used to trace the chromatin structure.

#### Cell clustering analysis of RNA-MERFISH data

After obtaining the cell-by-gene RNA count matrix from the “Decoding algorithm for RNA-MERFISH” section above, we preprocessed the matrix and performed cell clustering analysis to resolve distinct MOp cell types using the Scanpy pipeline (*98*), as described previously (*50*).

Preprocessing of the count matrix includes the following steps briefly described below. (1) Because the cell and cell-nucleus segmentation approach that we used can generate a small fraction of spurious artifacts, we removed the segmented cells that had a volume that was either less than 100 μm^3^ or larger than 5000 μm^3^. We also removed the segmented cells where less than 10 genes or 20 RNA counts were detected, which can arise either from segmentation artifacts or from poor imaging quality for a small number of FOVs due to the buffer flow issue during the RNA-MERFISH run. (2) We additionally removed potential doublets of cells using Scrublet (*99*): candidate cells with a doublet score higher than 0.3 were removed, which accounted for approximately 0.3% of the total cell number after step 1. (3) Next, we normalized the total RNA counts for each cell to the median total RNA counts of all cells. (4) After count normalization, the cell-by-gene count matrix was further log1p-transformed and converted to z-scores.

After preprocessing the cell-by-gene matrix above, we performed dimensionality reduction of the matrix using principal component analysis (PCA) and used the first 30 principal components.

We then performed graph-based community detection in the 30 principal components space, with nearest neighborhood size parameter k = 10, as previously described (*50*). We ran the Leiden clustering method to cluster the cells with the resolution parameter r = 0.5, which identified 23 clusters that corresponded well to the MOp cell types at the subclass level obtained in our previous study (*50*). We next manually curated these identified 23 clusters to match the consensus MOp subclass labels by merging a few small clusters to their adjacent large clusters and by splitting two clusters into six smaller clusters using a higher Leiden resolution parameter r = 0.1. The basis for merging and splitting was obtained from both inspecting the marker gene expression in these identified clusters, and from performing one round of cell type correspondence analysis (see the “Correspondence between clusters identified by different MERFISH protocols” section below). These curated clusters were then assigned into cell types at subclass level as described in the main text.

For presentation, Uniform Manifold Approximation and Projection (UMAP) was used to embed the cells in two dimensions using the same principal components that were used for clustering.

#### Correspondence between clusters identified by different MERFISH protocols

Correspondence between cell clusters identified by RNA-MERFISH in this study without tissue clearing (referred to as “uncleared RNA-MERFISH” below) and by RNA-MERFISH using tissue clearing in our previous study (*50*) (referred to “cleared RNA-MERFISH”) was assessed by running a neural-net classifier, as previously described (*50*). Briefly, the pre-normalized count matrices from the uncleared RNA-MERFISH in this study and the cleared RNA-MERFISH from the prior study were both log1p-transformed and z-scored. Then, the shared 242 genes measured in both studies were used to train a multi-layer perceptron (MLP) model, which was next used to predict cleared RNA-MERFISH cluster labels for each cell in the uncleared RNA-MERFISH dataset. As such, each cell in our uncleared RNA-MERFISH datasets had both a predicted cleared RNA-MERFISH cluster label and a cluster label determined from our *de novo* clustering above (see the “Cell clustering analysis of RNA-MERFISH data” section above). Cells were grouped based on their *de novo* cluster identity, and then the fraction of cells from a given *de novo* cluster identity that were predicted to have each cleared RNA-MERFISH cluster label was then determined to generate a confusion matrix, as shown in fig. S2D. Likewise, the same classifier approach was used for assessing our initial cell clustering result before any manual curation, as described in the “Cell clustering analysis of RNA-MERFISH data” section.

#### Bulk RNA sequencing data analysis of wild-type mouse brains

The bulk RNA sequencing data was obtained from Ref. (*100*). For each gene from the shared 238 genes between the bulk RNA sequencing and the uncleared RNA-MERFISH data, the FPKM per gene from the bulk RNA sequencing was calculated. 231 out of the 238 genes whose FPKM is larger than zero in the bulk RNA sequencing were kept and log10-transformed. The log10-transformed FPKMs of these shared genes were compared to their average log10-transformed RNA count per cell from the uncleared RNA-MERFISH in this study, as shown in fig. S2B.

#### 3D spatial distances between locus pairs from DNA-MERFISH data

For each decoded genomic locus that had been picked into a chromatin fiber, its 3D coordinate was calculated as the average of the fitted 3D Gaussian centers of the spot trio (or spot pair) decoded for that locus (see the “Decoding of fitted DNA spots” section above). The 3D spatial distance between any pair of genomic loci in each single cell was simply calculated as their Euclidean distance. The pairwise spatial distance matrix (hereafter referred to as pairwise distance matrix) can therefore be generated for all pairs of genomic loci of interest in each single cell, either at chromosome level (for all locus pairs from the given chromosome) or at the genome level (for all locus pairs across the mouse genome, including Chr1-19 and ChrX). To obtain the cell-type medians of the pairwise distance matrices, we aggregated pairwise distance matrices from single cells that belong to the same cell type. We then calculated the medians for each element in the pairwise distance matrices at chromosome level, as shown in Fig. 2A and fig. S3A.

#### snm3C-seq data analysis of wild-type mouse brains

Cell-type-specific snm3C data from the mouse neocortex was obtained from Ref. (*29*). The cells from the mouse neocortex subregion, as noted in the associated metadata from the paper, were used. To compare DNA-MERFISH data and snm3C data (as shown in fig. S3), we created bins for the snm3C data centered around the genomic regions that we imaged with DNA-MERFISH and procured the corresponding pairwise contact matrix for single cells by summing the pairwise contact counts per bin from the snm3C data. The bin size used was 500kb upstream to the start plus 500kb downstream to the end of each targeted imaged locus (which is about 15kb in length). We used a 500kb bin size in this case because the detected snm3C contacts are relatively sparse for long-range interactions. To obtain average pairwise contact matrices for selected MOp cell types, we aggregated pairwise contact matrices from single cells that belong to the cell type of interest and calculated the cell-type means for each element of the pairwise contact matrices, as shown in fig. S3A. The corresponding pairwise distances derived from DNA-MERFISH, and the number of contacts derived from snm3C were next compared by the Pearson correlation analysis and were plotted on the log10 scale for each chromosome for the cell types selected, as shown in fig. S3B.

#### Estimation of nuclear volumes of cells from DNA-MERFISH data

The volume of the cell nucleus was estimated by generating the minimal 3D convex hull surface (using Python’s SciPy package) from the 3D locations of all decoded chromosome loci in a given cell. Because a fraction of cells did not have the whole nucleus included in a 10-μm-thick tissue slice, we only included cells that had a number of decoded chromatin loci that was larger than a set cutoff number of 1250. We picked this number by varying the cutoff number from 400 to 2500 (which were ∼10% to 60% of the recovery efficiency of all 1981 targeted genomic loci assuming two copies per loci) and by inspecting the changes in the median of nucleus size for all L2/3 IT neurons. After fitting a polynomial function, we found that the median of the cell-nucleus size reached a turning point when increasing the cutoff number to 1250. A higher cutoff number than 1250 dramatically decreased the number of qualified cells and the remaining qualified cells started to contain more merged cell-nucleus objects (“doublets” as assessed by inspection of the ploidy number of chromosome homologs). Therefore, we used 1250 as the cutoff number when calculating the cell-type medians of the nuclear volume per cell for each MOp cell type, as shown in Fig. 1, D, F to H; fig. S6B.

#### snRNA-seq data analysis of wild-type mouse brains

The snRNA-seq data of MOp cell types from wild-type mouse brains was obtained from Ref. (*52*). The snRNA-seq 10x v3 data from this paper was used for our analyses because it was reported to have a higher number of genes recovered than other snRNA-seq methods in this paper (*52*). The cell-by-gene RNA count matrix from the snRNA-seq v3 data was preprocessed and analyzed for cell clustering similarly as described in the “Cell clustering analysis of RNA-MERFISH data” section above. The identified cell clusters showed a high correspondence to their original annotation in the paper (*52*). Thus, we directly used the original cell cluster annotation for procuring cell-type-specific RNA expression profiles because (1) this was derived from the consensus clustering results for all RNA-seq datasets in the prior paper (*52*) and (2) this was cross-validated from the cleared RNA-MERFISH from the prior study (*50*).

To calculate cell-type-specific RNA expressions for mouse MOp cell types, we used two different cell-by-gene RNA count matrices after basic filtering: which are (1) normalized, log-transformed, and z-scored RNA expression matrices, and (2) unnormalized RNA expression matrices. The normalized expression matrices were primarily used for standardized cell clustering as described above, and for DE gene analyses used in Fig. 3, E and F; fig. S11. Because we found that many aspects of chromatin organization changes were related to total transcription activity differences among cell types, the unnormalized expression matrices were used for most analyses in this study unless otherwise stated.

#### snATAC-seq data analysis of wild-type mouse brains

The snATAC-seq data of MOp cell types from wild-type mouse brains was obtained from Ref. (*52*). The original consensus cell cluster annotation derived from this paper was also used for procuring cell-type-specific ATAC expression profiles. The unnormalized cell-by-region ATAC count matrices were used for most analyses in this study unless otherwise stated.

#### Estimation of total transcriptional activity and chromatin accessibility per cell from wild-type mouse sequencing data

To estimate total transcriptional activity and chromatin accessibility per cell for MOp cell types, we used the unnormalized expression matrices from the “snRNA-seq data analysis of wild-type mouse brains” section and “snATAC-seq data analysis of wild-type mouse brains” section above, respectively. The number of unique molecular identifiers (UMIs) per cell captured from the corresponding sequencing analyses, which are typically also the total counts per cell of the unnormalized expression matrices, were used as total transcriptional activity and chromatin accessibility per cell. The cell-type medians of total transcription activity and chromatin accessibility per cell were then calculated for each cell type and shown in Fig. 1, E to G. To estimate the total number of accessible regions pe cell, the number of regions that had an ATAC read for each single cell were calculated and the corresponding cell-type medians were derived and shown in Fig. 1H.

#### Estimation of total transcriptional activity per cell in human brains

To estimate total transcriptional activity per cell for human brain cell types, we used two prior datasets as follows. The first dataset was the snRNA-seq from the human M1 brain region, which was obtained from Ref. (*59*). The original cell cluster annotation derived from this paper was used and adjusted to match the corresponding cell subclasses between human and mouse. The number of UMIs per cell captured were used to estimate the cell-type medians of total transcription activity per cell as described above for mouse brains, which are shown in fig. S5A.

The second dataset was the RNA-MERFISH data of 4000 genes in the human MTG brain region, which was obtained from Ref. (*49*). The original cell cluster annotation derived from this prior paper was used and adjusted to match the corresponding cell subclasses between human and mouse. The total RNA counts per cell detected from RNA-MERFISH were used to estimate the cell-type medians of total transcription activity per cell, as shown in fig. S5B.

#### Estimation of chromosomal territory size from DNA-MERFISH data

Chromosomal territory sizes in figs. S6 and S7 were estimated using the radius of gyration function of all decoded chromosome loci from any given chromosome. For each MOp cell type, the cell-type medians of chromosomal territory size were then calculated by aggregating all measurements of single chromosomes. For fig. S6, median chromosomal territory size for each chromosome type (e.g., Chr1-19 and ChrX) were calculated separately by aggregating chromosomes across individual cells in the given cell type that belonged to that chromosome type, whereas for fig. S7, median chromosomal territory sizes were calculated without distinguishing the chromosome type.

#### Chromosome territory segregation score from DNA-MERFISH data

Chromosomal territory segregation score in fig. S6C was used to quantify the degree of spatial separation between different chromosomal territories. To calculate this quantity, we adopted a previously developed method (*24*), which computes the spatial separation of chromosomal domains. Briefly, we treated two different chromosomes as two domains. Between two domains, we used median pairwise distances to first calculate an intra-domain distance distribution by considering all spatial distances between each pair of chromatin loci within the first domain and all distances between each pair of chromatin loci within the second domain. We then similarly calculated an inter-domain distance distribution by considering all distances between pairs of chromatin loci that reside in different domains. We then defined the segregation score as the median of all inter-domain distances divided by the median of intra-domain distances. Two highly intermixed domains (which are chromosome territories in this case) would have segregation score close to 1, while domains that are just contacting will have a segregation score of substantially larger than 2.

#### Estimation of chromosomal transcription per cell from wild-type mouse sequencing data

To estimate of chromosomal transcription (total transcript counts per chromosome) per cell, we used the unnormalized expression matrices as described in the “Estimation of total transcription activity and chromatin accessibility per cell from wild-type mouse sequencing data” section above, except for that genes were grouped by their genomic location on chromosomes. The corresponding total counts (UMIs) from genes for each chromosome per cell were then used to calculate the cell type medians of chromosomal transcription per cell, as shown in fig. S6, D and E.

#### Proximity frequency for genomic-locus pairs from DNA-MERFISH data

To calculate the proximity frequency for genomic-locus pairs, we first counted the number of locus pairs whose measured spatial distance in 3D was smaller than a set cutoff distance. The proximity frequency for a genomic-locus pair was determined by dividing the number of such qualified locus pairs with spatial distance smaller than the cutoff value by the total number of detected locus pairs for the same pair of target genomic loci. We then grouped the target genomic-locus pairs by their genomic distances, and determined distributions of the proximity frequency overall genomic-locus pairs included in each genomic-distance range for each cell type, as shown in Fig. 2C. We selected 750 nm as the set cutoff distance to match the cutoff distance used for other analyses (see the “A/B compartment analysis from DNA-MERFISH” section below). In addition, we also tested 500 nm as the cutoff distance, as in Ref (*24*), and we observed a similar trend as that shown in Fig. 2C.

#### Normalized insulation scores of megadomain-like structures from DNA-MERFISH data

Insulation score has been previously defined for megadomains in inactive X chromosomes in ensemble Hi-C (*61, 101*). We therefore used a similar method for megadomain analysis for our DNA-MERFISH data. Briefly, for each chromosome locus in a given chromosome, we selected upstream and downstream loci with a fixed window size (8 loci and ∼10Mb). We treated these two chromatin regions, up- and down-stream of the selected locus, as two “domains’’. We then computed the insulation score between these two regions as described in the “Chromosomal territory segregation score from DNA-MERFISH” section above. We then defined the normalized insulation score as the difference between the median of inter-region distances and the median of intra-region distances normalized by the sum of these two median values.

Therefore, a normalized insulation score will always be between 0 and 1, with the value 1 indicating strong insulation between the two regions. With this definition of normalized insulation score, we applied a sliding window along the chromosome to calculate the normalized insulation scores for each genomic loci in a chromosome. The local maxima then represent the boundaries between potential megadomain-like structures in the given chromosome, as shown in fig. S8B. Additionally, the interquartile range (IQR) of the normalized insulation scores for each chromosome was determined and the distribution of the IQR values across different chromosomes was used to estimate the prevalence of megadomain-like structures for each cell type, as shown in Fig. 2D.

#### Estimation of transcriptional activity and chromatin accessibility for each imaged DNA locus from sequencing data

To approximate the cell-type medians of transcriptional activity and chromatin accessibility for each imaged DNA locus, we used the unnormalized expression matrices from the “snRNA-seq data analysis of wild-type mouse brains” section and “snATAC-seq data analysis of wild-type mouse brains” section above, respectively. Specifically, we created bins centered around each imaged locus and summed the number of all RNA or ATAC counts for each bin in single cells. Next, for each cell type, we aggregated single-cell measurements to calculate the cell-type median value of each locus. We used 4Mb as the bin size because we found that it generated the highest correlation with chromosome compartment principal component (PC) values derived from our DNA-MERFISH (e.g., fig. S10, see the “A/B compartment analysis from DNA-MERFISH” section below for details). We then analyzed the cell type medians of transcriptional activity or chromatin accessibility for each imaged locus of interest with other chromatin features that were measured by DNA-MERFISH, whose results were used in Fig. 2, E and F; Fig. 3C; Fig. 4, B to E; fig. S9; fig. S10; fig. S14 for example.

After obtaining transcriptional activity and chromatin accessibility for each imaged locus, we additionally grouped each locus into different categories (e.g., “high”, and “low”) based on the percentile thresholds (<25th, and >75th percentiles respectively) of the corresponding transcription activity or chromatin accessibility distribution of all imaged loci, as described and shown in the Fig. 4, B to E; fig. S14, C and D.

#### Normalized pairwise distance matrices in selected MOp cell types from DNA-MERFISH data

To obtain normalized pairwise distance matrices as shown in fig. S8A, we first calculated the cell-type-specific median pairwise distance matrices as in the “3D spatial distances between locus pairs from DNA-MERFISH data” section. Then, we calculated the median value of all elements from each cell-type-specific median pairwise distance matrix for each MOp cell type and normalized the matrix by this median value such that this median value for every cell type was equal to that of the reference cell type (L2/3 IT).

#### A/B compartment analysis from DNA-MERFISH data

We performed A/B compartment analysis at major cell class level for neuronal cells (excitatory neurons and inhibitory neurons) given that the cell number of some neuronal subclasses were small. We also combined oligodendrocytes and oligodendrocyte progenitor cells (OPCs) as one major cell type (“Oligo” in Fig. 3, A to C; fig. S10) for this analysis. The A/B compartment analysis includes the following steps.

First, we calculated the pairwise proximity frequency matrices for each major cell type. To do this, we counted the total number of measured distances between any given pair of target genomic loci from all single cells belonging to the given cell type. The proximity frequency for any given genomic-locus pair was then determined as described in the “Proximity frequency for genomic-locus pairs from DNA-MERFISH data” section, using 750 nm as the cutoff distance.

We selected 750 nm as the cutoff distance because (1) the Pearson correlation coefficient between the calculated proximity frequency matrices and snm3C-derived contact matrices (*29*) remained high for cutoff distances between 200 nm to 800 nm, and (2) the cutoff distance of 750 nm retained a higher baseline level of pairwise proximity frequency and produced a less noisy Pearson cross-correlation matrices for compartment analysis as described below.

After we obtained the pairwise proximity frequency matrices for each major cell type as described above, we normalized these matrices by dividing the observed proximity frequency matrices over the expected proximity frequency matrices. The expected proximity frequency matrices were derived by calculating the expected frequency of observing a locus pair within 750nm distance as a function of their genomic distance with a DNA polymer model (*57*). This normalization is meant to remove the effect of genomic distances between target genomic loci such that the chromatin locus pairs that are closer to or farther from each other than expected from the polymer model can be identified.

Next, the Pearson cross-correlation matrices were calculated from the normalized proximity frequency matrices above, as previously described (*24*), as shown in Fig. 3A. The Pearson cross-correlation matrices were then subject to Principal Component Analysis (PCA), and from different principal component (PC) values were obtained for each imaged chromosome locus. To select the PC that corresponds to A/B compartments for each cell type, we calculated the Spearman correlation coefficients between the corresponding cell-type-specific median locus chromatin accessibility (calculated as in the “Estimation of transcriptional activity and chromatin accessibility for each imaged DNA locus from sequencing data” section above, using snATAC-seq from Ref. (*52*)) and PC values for the first three PCs. The signs of PC values for each chromosome were then assigned based on their signs of correlation with the corresponding chromatin accessibility. We picked the PC whose values had the highest correlation as the PC that corresponds to A/B compartments. Imaged loci whose selected PC values that were greater than 0 were assigned as compartment-A loci, whereas loci whose selected PC values that were equal to or less than 0 were assigned as compartment-B loci.

#### Differentially expressed (DE) gene locus selection for local A/B environment analysis

To select DE gene loci for the indicated cell type as shown in Fig. 3E; fig. S11, A and B, we used the normalized expression matrices from the “snRNA-seq data analysis of wild-type mouse brains” section above to find DE genes. For each given cell type (termed “reference cell type”), we performed “rank_genes_groups” analysis using the Scanpy package (*98*) to select the top 200 upregulated and top 200 downregulated DE genes relative to the rest of the cell using the Wilcoxon test. Next, we picked DE genes whose TSS are within 100kb of any of our imaged loci, and such loci that had at least one nearby DE gene were identified as DE gene loci.

#### Super-enhancer locus selection for local A/B environment analysis

The selection of super-enhancer loci for indicated cell types, as shown in Fig. 3F; fig. S11C, was based on super-enhancer locus panel design as described in the “Genomic locus selection for DNA-MERFISH” section above. For each given cell type (termed “reference cell type”), we selected the candidate super-enhancer loci that were specific to the given reference cell type as the super-enhancer loci for this analysis. In the meantime, we randomly selected 100 loci from our super-enhancer panel that were specific to the other cell types but not to the reference cell type as the randomized super-enhancer control loci.

#### Local A/B density ratio analysis

To calculate the local density ratio of compartment-A and compartment-B loci for each imaged locus as in Fig. 3, D to F; fig. S11, we first calculated the local density score for each locus in single cells through similar procedures as follows. The local trans-A density score for any given locus was defined as the sum of the Gaussian probability density function values from all compartment-A loci from other chromosomes. The local trans-B density scores were defined similarly.

Local A/B density ratio for any given locus was defined as the ratio between local trans-A density score and local trans-B density score and we then log2-transformed this ratio. After we obtained these local A/B density ratios in single cells, we then aggregated ratios from single cells that had more than 600 decoded loci (to filter out single cell measurements that were less accurate) for each cell type. The cell-type median values for each locus were then calculated and used for the following analyses. In this study, we defined local A/B density based on trans-A and trans-B density scores (i.e., contributions from other chromosomes) because we found that they better correlated with transcriptional activity across all of our imaged loci.

To study the change in A/B density ratio between cell types as shown in Fig. 3, E and F; fig. S11, we additionally performed linear regression between the density ratios from all targeted genomic loci for any given cell type and those for a common reference cell type (L2/3 IT in this study). Based on the linear regression results, for any given cell type, we then normalized its A/B density ratios by this linear regression results to correct for the intrinsic differences in A/B density ratios between cell types (i.e., some cell types had consistently higher A/B density ratios than other cell types for all imaged loci). Another reason for this normalization is because the DE genes and super-enhancers were also derived from normalized snRNA-seq and snATAC-seq count matrices, where cell-type-intrinsic differences in total transcription and chromatin accessibility had been normalized.

Finally, for each given cell-type-specific DE gene or super-enhancer locus, we subtracted its local A/B density ratios in other cell types from that in the cell type where the given locus was identified (see the “DE gene locus selection for local A/B environment analysis” section and “Super-enhancer locus selection for local A/B environment analysis” section above for details on how these loci were identified). We then divided the local A/B density ratio difference (calculated from the above-mentioned subtraction) by the local A/B density ratio in the cell type where the given locus was identified. This generated the percentage changes in the local A/B density ratio as shown in the heatmaps in fig. S11, B and C. For heatmaps shown in Fig. 3, E and F, for each reference cell type, the median of all A/B compartment density ratio percentage changes from all of its cell-type-specific DE gene (or super-enhancer) loci were shown.

#### Radial positioning analysis for imaged genomic loci from DNA-MERFISH data

To calculate the nuclear radial positions for individual imaged genomic loci in single cells, we first generated the minimal 3D convex hull surface surrounding all decoded chromosome loci in a given cell. Then for each DNA locus, we calculated its spatial distance to the centroid of the 3D convex hull and normalized this distance to a corresponding radius of the 3D convex hull to obtain the normalized radial position of the locus. Specifically, the line connecting the centroid and the locus of interest was drawn and its intersection with the convex hull surface was determined. The normalized nuclear radial position was calculated by normalizing the locus’ distance to the nuclear centroid over the distance between the nuclear centroid and the intersection point on the convex hull. Next, for each locus in each cell type, the cell-type median of its normalized nuclear radial positions was calculated from aggregated single-cell measurements, which were shown or used in Fig. 4; figs. S12 to S15.

After obtaining the cell-type median of nuclear radial position for each target genomic locus, we examined the relationship between transcription activity, chromatin accessibility, and local A/B density ratio with the nuclear radial position of the genomic loci as shown in Fig. 4B to E; figs. S13 and S14. We additionally grouped each locus into 5 different bins (every 20^th^ percentile of the distribution of radial position across all imaged loci) for each cell type. The top and bottom 20^th^ percentile were defined as the loci near the nuclear periphery and loci near the nuclear interior/center, respectively, for each cell type. Features such as the status of long gene expression (see the “Long gene expression analysis” section below for details) were assessed for loci near the nuclear periphery or near the nuclear interior/center as shown in fig. S15.

#### Long gene expression analysis

To examine whether a given locus contained a long gene (>300kb) that was highly expressed, as in fig. S15, we defined an active transcription cutoff value as the 90^th^ percentile of the distribution of mean transcription level per cell across all genes from snRNA-seq data (*52*). Genes whose mean transcription level per cell is above this cutoff value were considered as highly active.

To characterize the associated function of long genes that are highly expressed in fig. S15, we further identify highly active long genes that were also located near the nuclear periphery in neurons. Next, we performed Gene Ontology analysis (g:GOSt analysis) using the online g:Profiler tool (https://biit.cs.ut.ee/gprofiler/gost) of these genes for each sub-ontologies (*102*). Results of g:GOSt analysis were downloaded from the online g:Profiler tool and re-plotted in fig. S15.

#### Estimation of histone mark levels for each imaged DNA locus from sequencing data

To approximate cell-type-specific histone mark levels for each imaged DNA locus as used in Fig. 5, A and B; fig. S14, E and F, we similarly created bins centered around each imaged loci and summed the reads of the corresponding histone mark for each bin as described in the “Estimation of transcriptional activity and chromatin accessibility for each imaged DNA locus from sequencing data” section above. The cell-type-resolved single cell histone profiles (H3K9me3, H3K27ac, H3K4me3) were obtained from the Paired-Tag data from Ref. (*63*).

#### Analysis of transcriptional regulation by MeCP2 from single-cell RNA sequencing data

For analysis of the relationship between radial positioning and transcriptional regulation by MeCP2 for cell types in the cortex, single-cell RNA sequencing results from the visual cortices of WT *Mecp2 +/y* and mutant *Mecp2-/y* mice were downloaded from GEO (GSE113673) (*71*). We used similar normalization, dimensionality reduction, and clustering analysis as described in the “Cell clustering analysis of RNA-MERFISH data” section. We used marker genes *Camk2a*, *Olig1*, *Cx3cr1* and *Cldn5* to identify clusters corresponding to excitatory neurons, oligodendrocytes, microglia, and endothelial cells, respectively.

For DE gene analysis, raw count matrix was used, followed by per-cell RNA count normalization and log transformation. DE genes between WT and *Mecp2* KO cells were identified within each cell type by the “rank_genes_groups” function in Scanpy (*98*) using the Welch’s t test. Cell-type-specific DE genes between WT and *Mecp2* KO cells (adjusted *p* value <=0.05) were then grouped based on the direction of gene expression changes, as MeCP2-activated (transcription-downregulated upon *Mecp2* deletion) and MeCP2-repressed (transcription-upregulated upon *Mecp2* deletion) (Fig. 5C). Differential expression (DE) scores calculated from “scanpy.tl.rank_genes_groups” were used for analysis in Fig. 5D to circumvent the effect from lowly expressed genes. Specifically, the DE score of a gene is the t-statistics score when comparing the expression-level distributions of WT and KO cells using Welch’s t test.

The nuclear radial positioning for each gene was estimated by using the normalized nuclear radial position of its closest imaged locus, in a cell-type-specific manner (obtained from the WT dataset, see the “Radial position analysis for imaged genomic loci from DNA-MERFISH data” section above). However, if the genomic distance between the closest imaged locus and the gene is greater than 3Mb, we excluded the gene from further analysis. For the analysis of the relationship between nuclear radial positioning and transcriptional regulation by MeCP2, we divided the genes into ten equal bins based on their normalized nuclear radial position. The average and 95% confidence interval of transcription level changes from DE gene analysis as a function of the normalized nuclear radial position were then plotted for each bin in Fig. 5D.

#### Analysis of transcriptional regulation by MeCP2 from bulk RNA sequencing data

Published bulk RNA sequencing data from *Mecp2* deletion or mutations were downloaded from GEO (GSE128186 (*73*), GSE152800 (*74*) and GSE139033 (*41*)). For all three datasets, genes that have at least one count per million reads across all samples in a given comparison were used for further DE gene analysis. For the dataset from Ref. (*74*), the WT and MM2 mutations from two age groups were combined for analysis. Pydeseq2 (*103*) with default settings was used to perform DE gene analysis. For datasets from Ref. (*73*) and Ref. (*74*), because they contain bulk RNA sequencing from forebrain and hypothalamus tissues, respectively, the nuclear radial position of each gene was estimated by the median of normalized nuclear radial positions of the gene’s closest imaged locus (with a 3Mb threshold) for all cell types (Fig. 5, F and G) (obtained from the WT dataset, see the “Radial positioning for imaged genomic loci from DNA-MERFISH” section above). For the dataset from GSE139033 (*41*), because Ngn2-induced neurons are excitatory neurons (*104*), the nuclear radial position of each gene was estimated by the median of normalized nuclear radial positions of the gene’s closest imaged locus (with a 3Mb threshold) for excitatory neurons only (Fig. 5H). Genes were binned into ten equal bins based on their normalized nuclear radial positions similar to the “Analysis of transcriptional regulation by MeCP2 from single-cell RNA sequencing data” section above.

#### Identification of WT and *Mecp2* KO cells from *Mecp2 +/-* female mice

The mean intensity of MeCP2 immunofluorescence signals within each segmented cell was first calculated and log_10_ transformed. Then, the mean MeCP2 intensity of a given cell was normalized by the mean intensity across all cells belonging to the same cell type and imaged during the same experiment. Such a normalized mean MeCP2 intensity helped to adjust the differential MeCP2 levels among different cell types (*72*) and also adjust the batch effects. Local relative MeCP2 signal was calculated as the log_2_ ratio of the mean MeCP2 intensity within each segmented cell divided by the mean MeCP2 intensity of the background surrounding the cells. The background region of interest was defined as a ring area with 10-pixel thickness around the segmented cell. Cells with normalized MeCP2 mean intensity less than 0.92 and local MeCP2 signal less than 0.2 were defined as KO cells and cells with normalized MeCP2 mean intensity greater than 0.98 and local MeCP2 signal greater than 0.5 were defined as WT cells (see fig. S16A). The rest of the cells (∼20%) remained un-determined and were not used in further analyses.

#### Analysis of MeCP2’s effect on chromatin organization

The nuclear volume of cells in *Mecp*2+/- mice was estimated as described in the “Estimation of the nuclear volume from DNA-MERFISH” section above (fig. S17). For unnormalized distance to nucleus center, we calculated the distance of each genomic locus to the nuclear center by generating the minimal 3D convex hull surface of all decoded loci in each cell and calculating the Euclidean distance between the locus of interest and the centroid of the 3D convex hull (Fig. 6C and fig. S18B). The normalized nuclear radial position was calculated in the same manner as described in the “Radial position analysis for imaged genomic loci from DNA-MERFISH data” section above (Fig. 6C and fig. S18B). Local A/B density ratio changes upon *Mecp2* deletion were calculated as described in the “Local A/B density ratio analysis” section above (Fig. 6, D to F). For analysis of the positional dependent effect of *Mecp2* deletion, the genes or loci were grouped into 20 equal bins based on its cell-type-specific normalized radial positioning calculated from wild-type cells (Fig. 6 and fig. S18B).

#### Analysis of MeCP2’s effect on A/B compartment

Cross-correlation matrices for each chromosome in each major cell type (excitatory neurons, inhibitory neurons, astrocytes, oligo cells, microglia, and endothelial cells) were calculated as described in the “A/B compartment analysis from DNA-MERFISH” section (fig. S19A). The elements in the upper triangle of the cross-correlation matrices were then vectorized, and the Pearson correlation coefficient between the WT and *Mecp2* KO vector representation of the cross-correlation matrices for each chromosome in each major type was then calculated (fig. S19B).

## Supplementary Figures

**Fig. S1.**
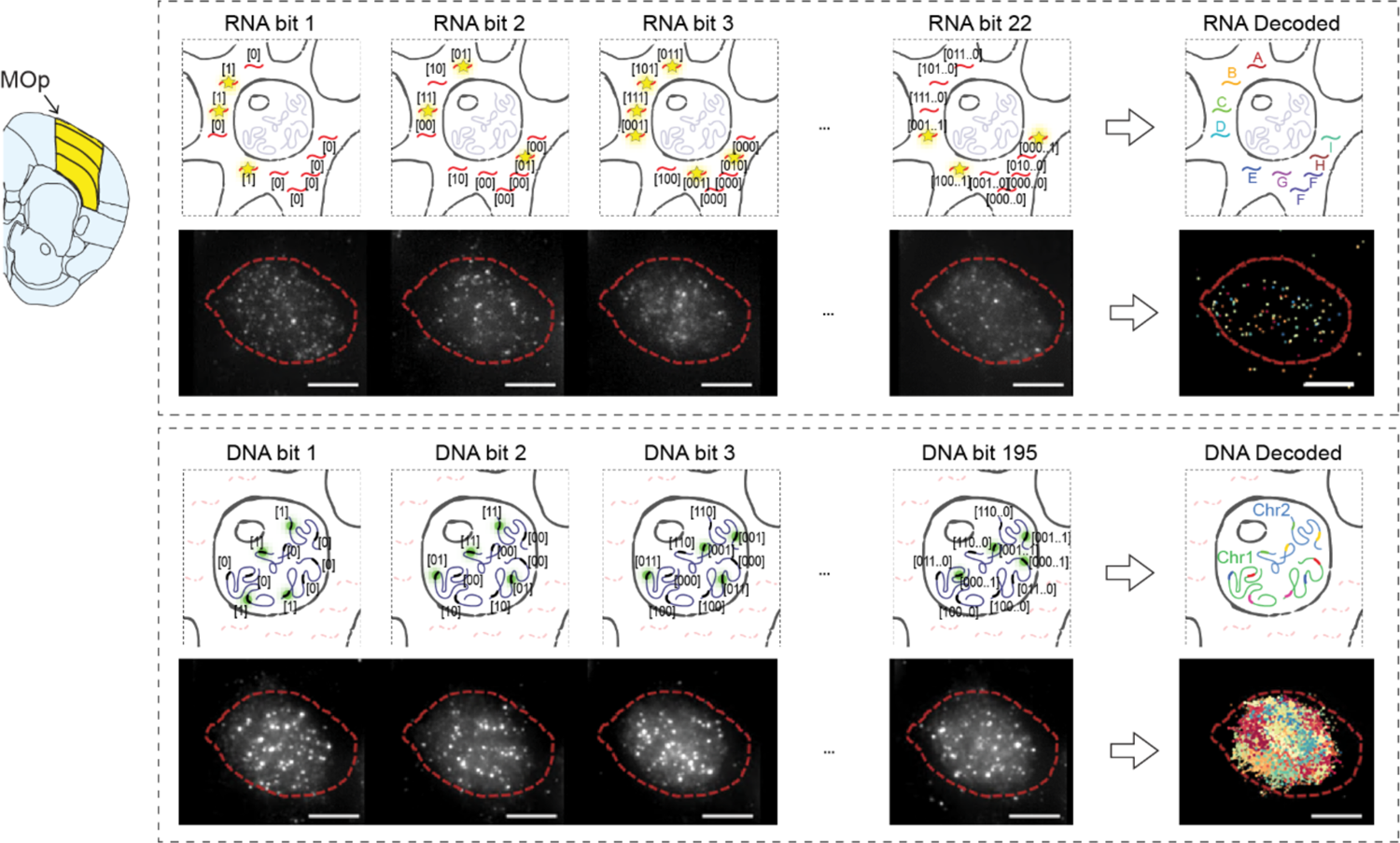
Experimental scheme of integrated RNA- and DNA-MERFISH. Schematics of integrated RNA-MERFISH and DNA-MERFISH. Left: Schematic annotation of the mouse MOp according to the Allen Common Coordinate Framework version 3 (http://atlas.brain-map.org/) (*88*). Right, Top: Schematic of RNA-MERFISH (top) and per-bit RNA-MERFISH images of an example cell (bottom). Right, Bottom: Schematic of DNA-MERFISH (top) and per-pit DNA-MERFISH images of the same cell (bottom). Red dashed lines in the images indicate the area where RNA and DNA foci were visualized and decoded.

**Fig. S2.**
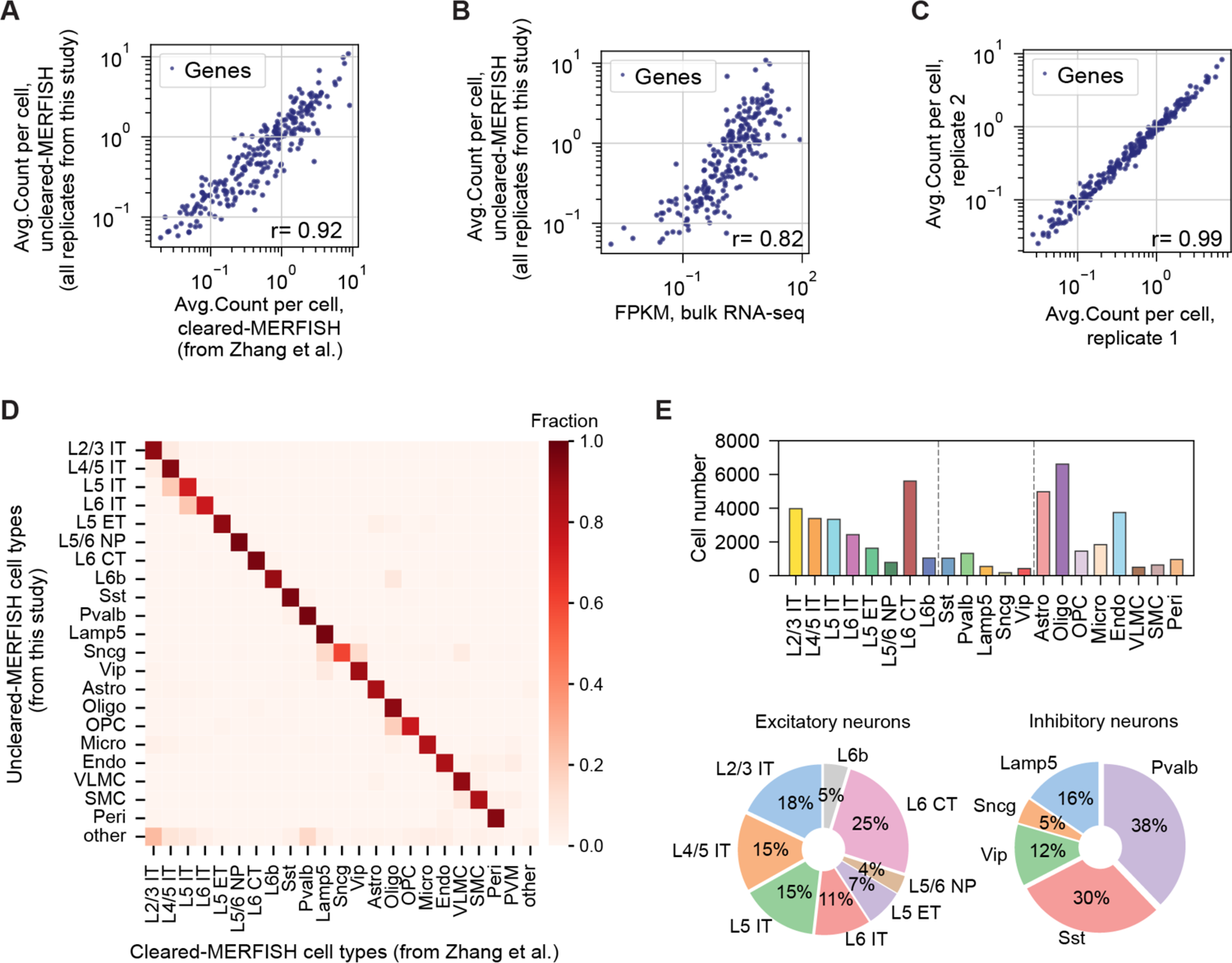
Performance of the modified RNA-MERFISH protocol in resolving molecularly defined cell types in the mouse MOp. (**A**) Scatterplot of the average copy number per cell of individual genes determined by RNA-MERFISH from our modified protocol without tissue clearing versus copy number per cell of individual genes determined by RNA-MERFISH with tissue clearing (*50*). (**B**) Scatterplot of the average copy number per cell of individual genes determined by RNA-MERFISH from our modified protocol versus expression level of the same genes determined by bulk RNA-seq (*100*). (**C**) Scatterplot of the average copy number per cell of individual genes determined by RNA-MERFISH from our modified protocol between two biological replicates. In (A to C), the Pearson correlation coefficients r are indicated in the plot. (**D**) Confusion matrix for correspondence between cell types determined by RNA-MERFISH in this study and cell types determined by a multi-layer perceptron (MLP) classifier trained by data from the previous RNA-MERFISH study with tissue clearing (*50*). Each element in the matrix represents the fraction of cells from the given cell type (x-axis) determined by the MLP classifier (see Materials and Methods) that was assigned to the individual cell type determined by *de novo* clustering in this study (y-axis). (**E**) Cell-type composition of the MOp. Top: Bar plots show the cell numbers for each MERFISH-identified cell type, combined from 4 biological replicates in this study. Bottom: Pie charts show the fractions of excitatory neurons (left) and inhibitory neurons (right) belonging to the indicated neuronal cell types, combined from 4 biological replicates. Our current analysis includes a larger area of the white matter and hence has a slightly different cell-type composition compared to our previous MERFISH analysis of MOp, which is focused on the grey matter (*50*).

**Fig. S3.**
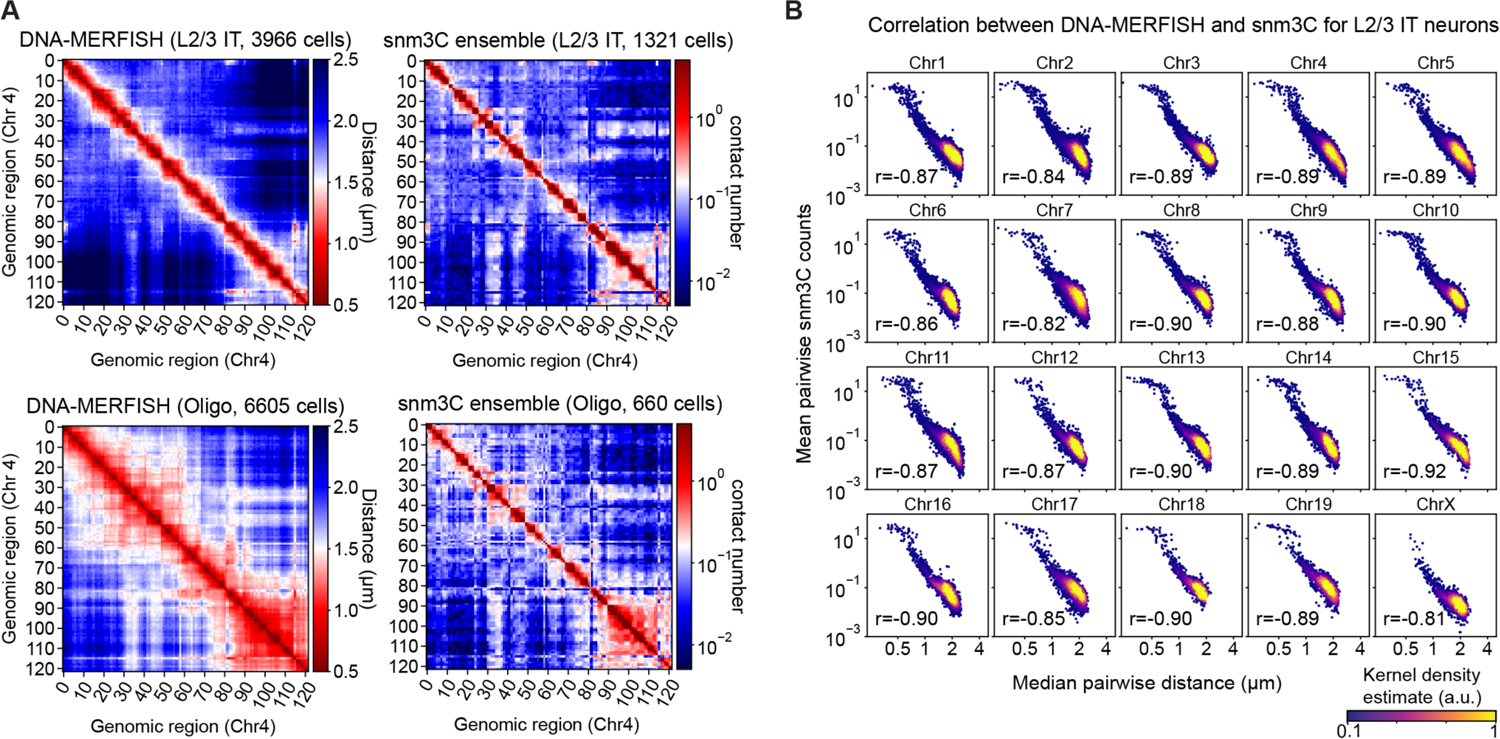
Comparison of 3D-genome organizations determined by DNA-MERFISH and snm3C. (**A**) 3D chromatin organization of an example chromosome (Chr4, 3.0Mb – 156.3Mb) revealed by DNA-MERFISH and snm3C (*29*), respectively. Left matrices: Median cis-chromosomal pairwise distance matrices of Chr4 for L2/3 IT neurons (left, top) and oligodendrocytes (left, bottom), derived from DNA-MERFISH data. Right matrices: Ensemble cis-chromosomal pairwise contact matrices of Chr4 for L2/3 IT neurons (right, top) and oligodendrocytes (right, bottom), derived from snm3C. (**B**) Scatterplots of the median cis-chromosomal pairwise distances derived from DNA-MERFISH data versus the mean contact counts from snm3C data (*29*) for all imaged locus pairs in the indicated chromosomes in L2/3 IT neurons. The kernel density in the scatterplots estimates the two-dimensional distribution. The Pearson correlation coefficients r are indicated.

**Fig. S4.**
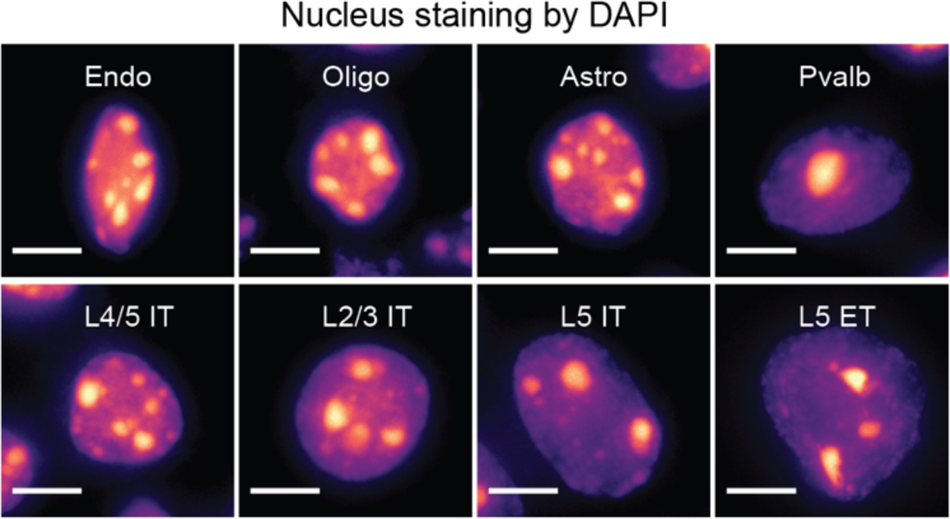
Cell-type-dependent variations in the cell-nucleus sizes visualized by DAPI staining in mouse brains. Representative maximum-projection images of DAPI staining are shown for several cell types. Scale bar: 5μm.

**Fig. S5.**
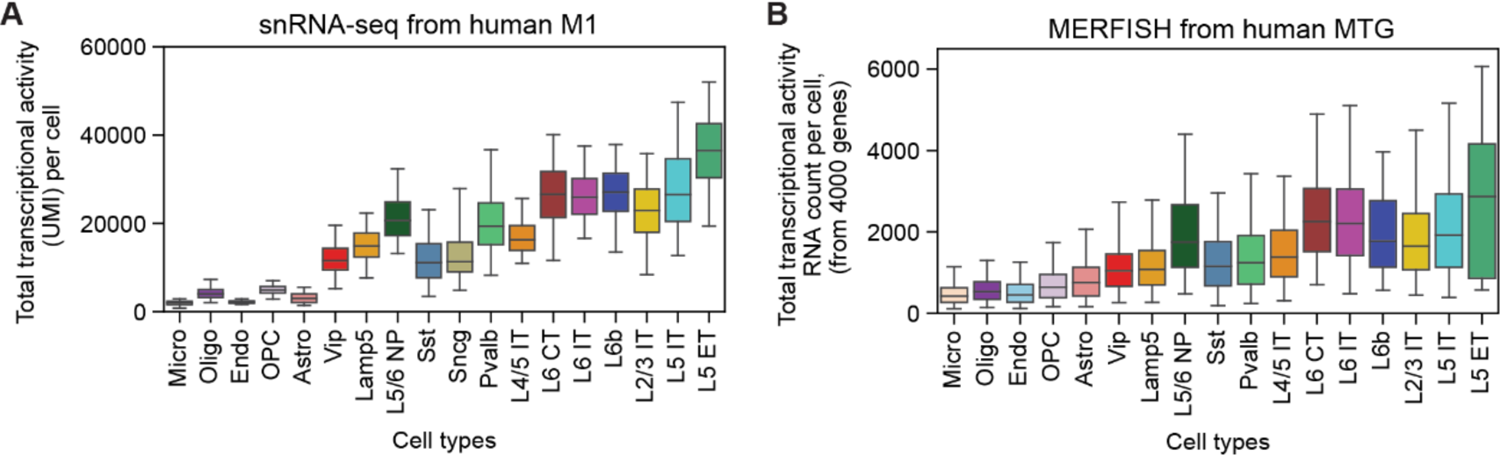
Cell-type-dependent variations in transcriptional activity in the human brain. (**A**) Boxplot for the distribution of total transcriptional activity across individual cells in each cell type in the human primary motor (M1) cortex. Total transcriptional activity per cell was calculated using the sum of UMIs from snRNA-seq data of the human M1 (*59*). (**B**) Boxplot for the distribution of total transcriptional activity across individual cells in each cell type in the human middle superior temporal gyrus (MTG). Total transcriptional activity per cell was estimated using the total RNA count per cell from 4000 genes measured by RNA-MERFISH (*49*). Cell types were ordered as in Fig. 1, D and E. The center line, box, and whisker represent the median, the 25th-75th percentile, and the 5th-95th percentile, respectively.

**Fig. S6.**
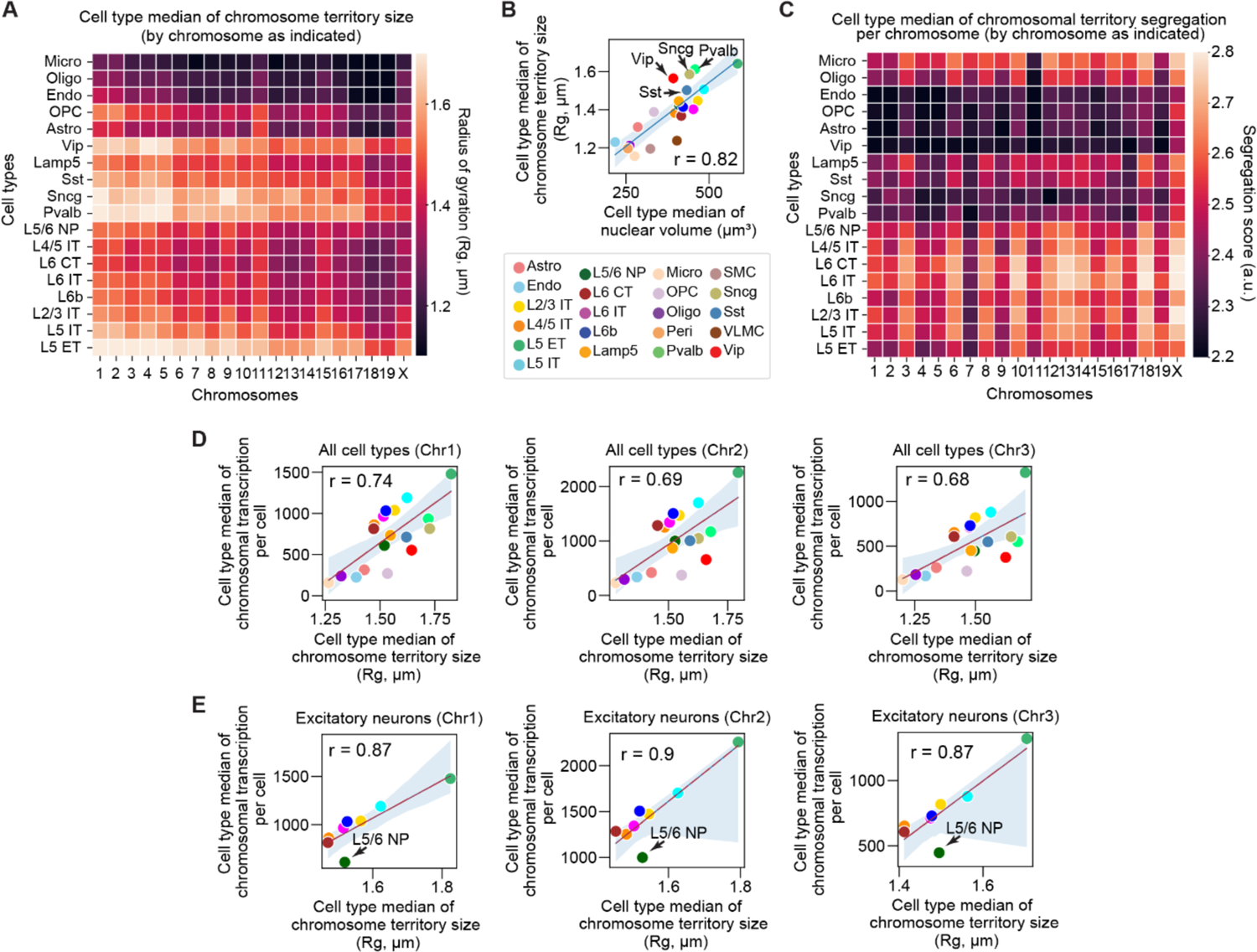
Cell-type-dependent variations in chromosomal territory sizes and transcriptional activity. (**A**) Heatmap of the median chromosomal territory sizes across individual cells in each cell type. Each pixel in the heatmap represents the median radius of gyrations (Rg) for the indicated chromosome (columns) within the indicated cell type (rows). (**B**) Scatterplot of the cell-type medians of chromosomal territory sizes versus nuclear volume. The chromosomal territory size per cell is calculated by averaging over all chromosomes (Chr1-19 and ChrX). Arrowheads indicate several inhibitory neuronal types that have larger chromosomal territory sizes on average compared to excitatory neuronal types with similar nuclear volumes. (**C**) Heatmap of the mean segregation scores of chromosomal territories across individual cells in all cell types. Each pixel in the heatmap represents the segregation score between the corresponding chromosome (columns) and all other chromosomes averaged over all cells in the indicated cell type (rows). Chromosomal territory segregation score was calculated based on the intermixing level between loci from the given chromosome and loci from other chromosomes (within the same cell) in 3D space (see Materials and Methods). (**D**) Scatterplot of cell-type medians of total transcriptional activity for Chr1 (left), Chr2 (middle), and Chr3 (right) versus the cell-type medians of the radius of gyration of the same chromosome. Total transcriptional activity of the indicated chromosome for each cell was calculated using the sum of UMIs of genes in this chromosome from snRNA-seq data (*52*). (**E**) Similar to (D), but for excitatory neurons only. Arrowheads indicate the L5/6 NP neuron that deviates from the correlation, which may be due to either the fewer cells of L5/6 NP in this analysis or a real cell-type difference. For scatterplots in (B), (D), and (E), the line and the shade represent the fitted linear regression line and the 95% confidence interval, respectively. The Pearson correlation coefficients r are indicated.

**Fig. S7.**
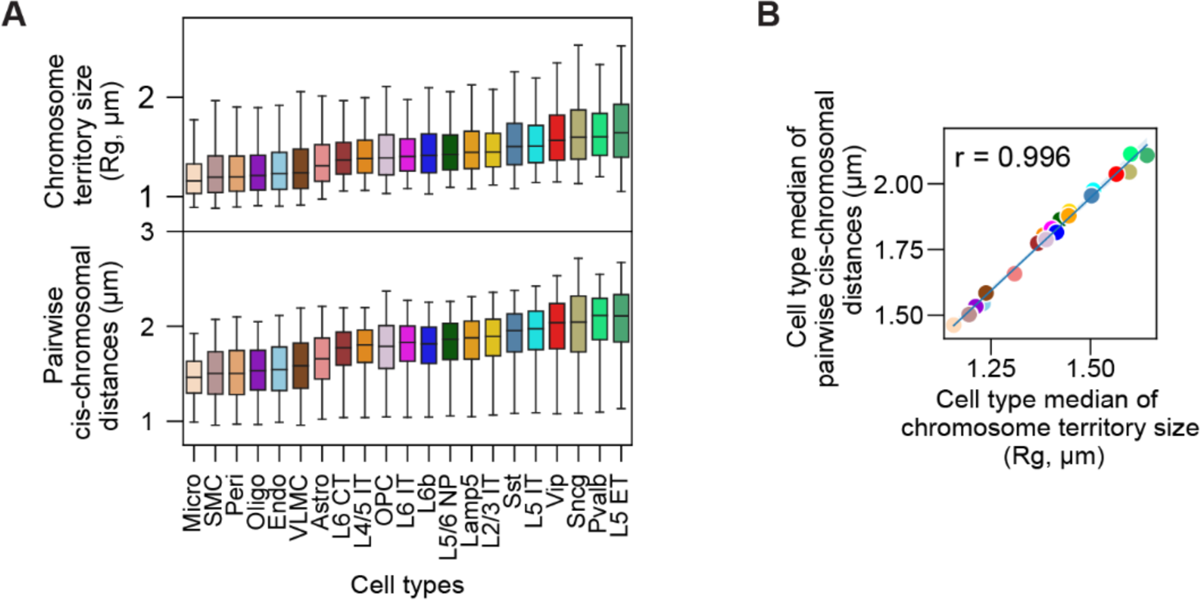
Relationship between the chromosomal territory sizes and cis-chromosomal pairwise distances. (**A**) Boxplots for the distributions of chromosomal territory sizes across individual chromosomes measured in all individual cells (top) and for the distributions of median cis-chromosomal pairwise distances across individual locus pairs (bottom) in each cell type. Each data point in the boxplot in the top panel represents a chromosomal territory size measurement from one chromosome in one cell, calculated by the radius of gyration (Rg). Each data point in the boxplot in the bottom panel corresponds to a median cis-chromosomal pairwise distance between a unique genomic locus pair across all individual cells for the indicated cell type. Cell types were ordered by their median chromosomal territory sizes for both top and bottom boxplots. The center line, box, and whiskers in the boxplot represent the median, 25^th^-75^th^ percentiles, and 5^th^-95^th^ percentiles respectively. (**B**) Correlation between cell-type medians of all unique cis-chromosomal pairwise median distance and individual chromosomal territory sizes. Each data point corresponds to one cell type.

**Fig. S8.**
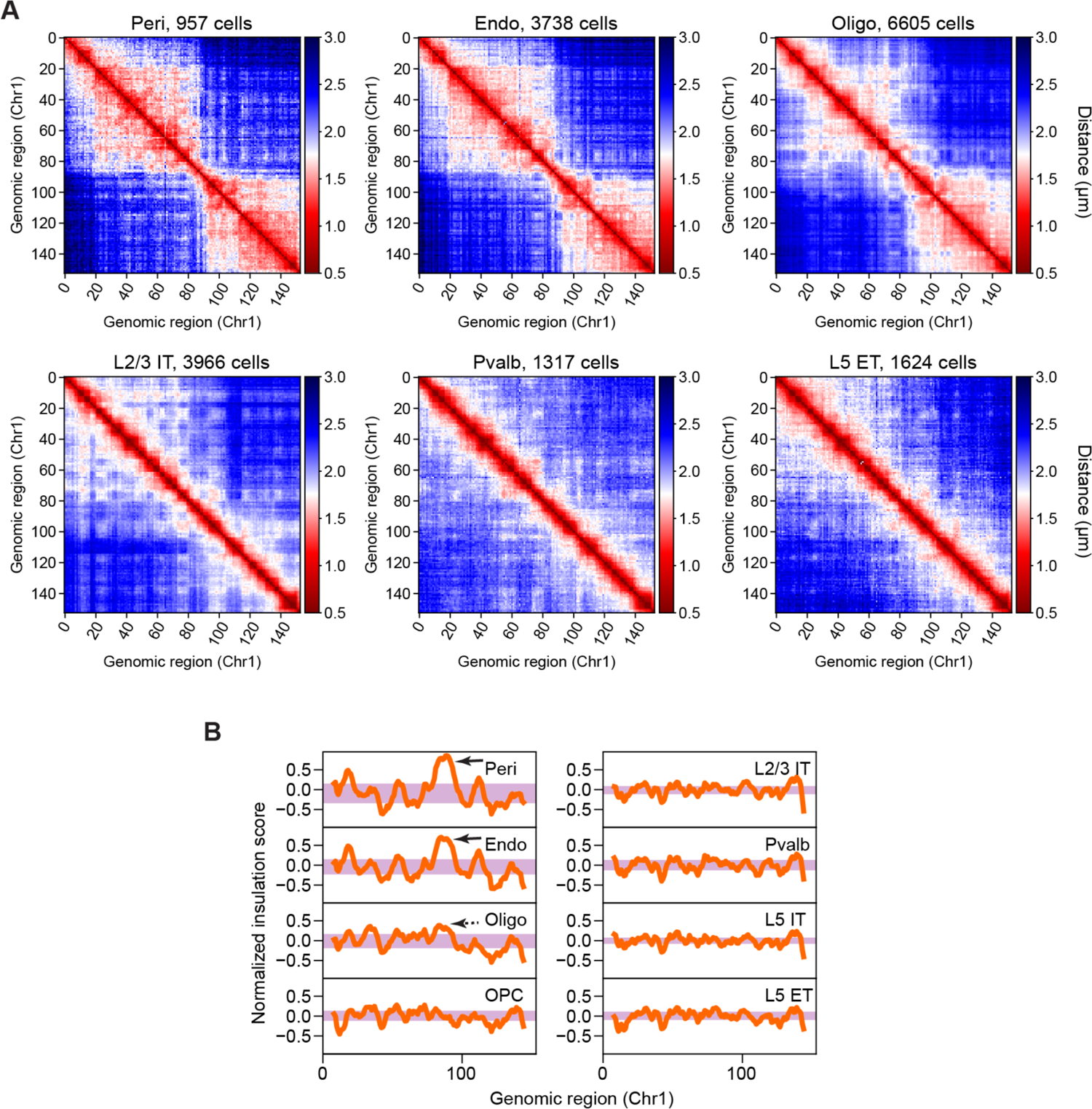
Different prominence of megadomain structures among different cell types. (**A**) Similar to Fig. 2A, but for normalized median cis-chromosomal pairwise distance matrices of Chr1 (3.7Mb – 98.8Mb), which were normalized to remove the difference in chromosomal territory sizes (see Materials and Methods). (**B**) Normalized insulation scores (see Materials and Methods) along the genomic coordinate of Chr1 for the indicated cell types. Purple shades represent the interquartile ranges (IQRs, 25th-75th percentiles) of insulation scores. Arrows indicate the putative domain boundary with high insulation scores, separating the two megadomains on Chr1 in non-neuronal cells.

**Fig. S9.**
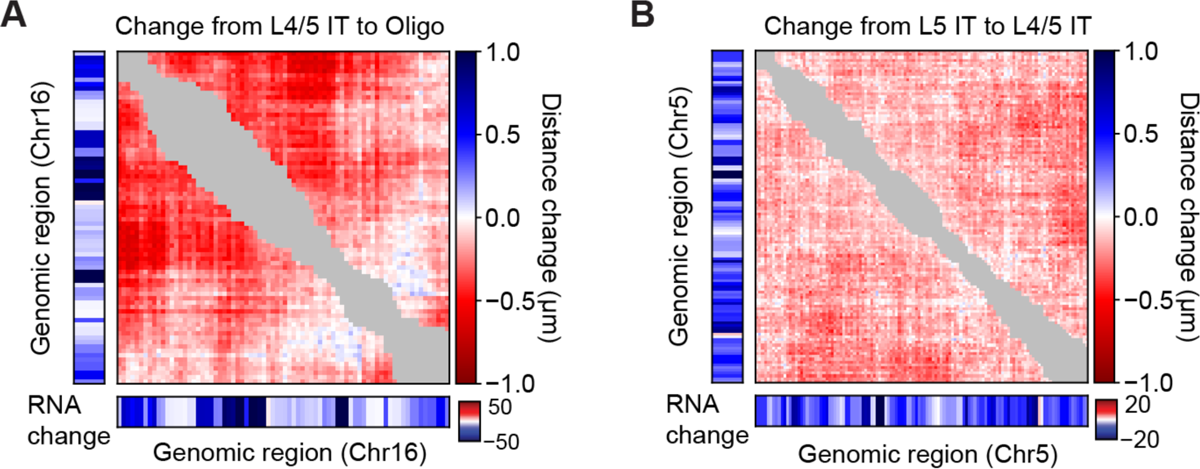
Consistent changes in higher-order chromosome organization and transcription between cell types. (**A**) Comparison of higher-order chromosome organization and transcription in Chr16 (3.7Mb – 97.7Mb) between L4/5 IT and oligodendrocytes. The matrix represents the change in median cis-chromosomal pairwise distance from L4/5 IT to oligodendrocytes. Next to the matrix (left and below) are changes in transcript counts of the corresponding genomic loci from L4/5 IT to oligodendrocytes (derived from snRNA-seq data in Ref. (*52*), see Materials and Methods). Gray elements near the diagonal represent locus-pairs whose genomic distances are <2Mb. (**B**) Similar to (A), but for comparison of higher-order chromosome organization and transcription in Chr5 (3.7Mb – 151.3Mb) between L5 IT and L4/5 IT.

**Fig. S10.**
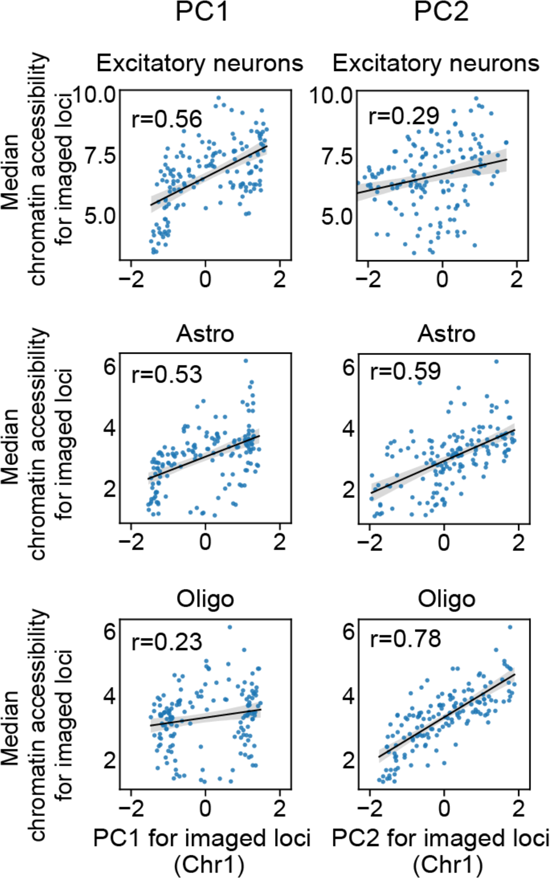
Scatterplots for the PC values of the cross-correlation matrix versus the cell-type median of chromatin accessibility of Chr1 for several major cell types. For each locus, the cell-type median of chromatin accessibility was calculated using snATAC-seq data (*52*) (see Materials and Methods) and the PC1 or PC2 values were calculated from the cross-correlation matrices as shown on Fig. 3A. The Spearman correlation coefficients r are indicated in each plot.

**Fig. S11.**
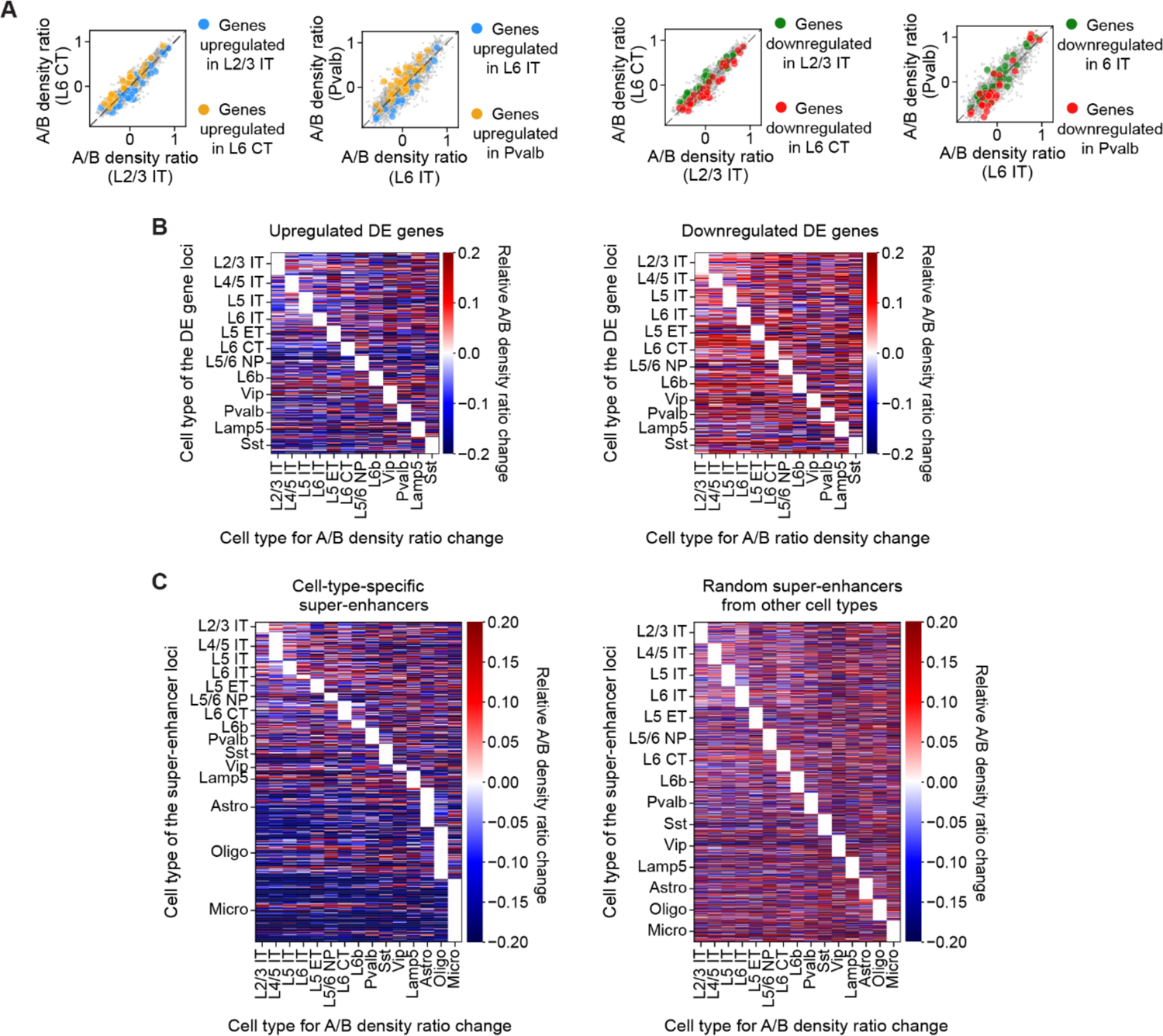
Correlation between local A/B-compartment chromatin environment and transcriptional activity of genes differentially expressed between different cell types, as well as activities of cell-type-specific super-enhancers. (**A**) Scatterplots of local A/B density ratio in one cell type versus another. DE (differentially expressed) genes are colored according to the legend. All other loci are colored in gray. Dashed line represents equality (y = x). For each cell type, DE genes and DE gene loci were identified as described in Fig. 3E. (**B**) A/B density ratio changes for cell-type-specific DE gene loci between the cell types where the DE genes were identified and other cell types. Left: each pixel in the heatmap represents the median value of the fractional change in the local A/B density ratio of an upregulated DE gene locus for each cell type (x-axis) over the reference cell type (y-axis) where the DE gene was identified. Right: similar to the left heatmap, but for downregulated DE gene loci. (**C**) Similar to (B), but for local A/B density ratio changes for cell-type-specific super-enhancer loci (left) or a random set of other super-enhancer loci selected regardless of the cell-type identity (right). Each pixel represents the changes of A/B density ratio for one super-enhancer locus. Super-enhancers were selected as described in Fig. 3F.

**Fig. S12.**
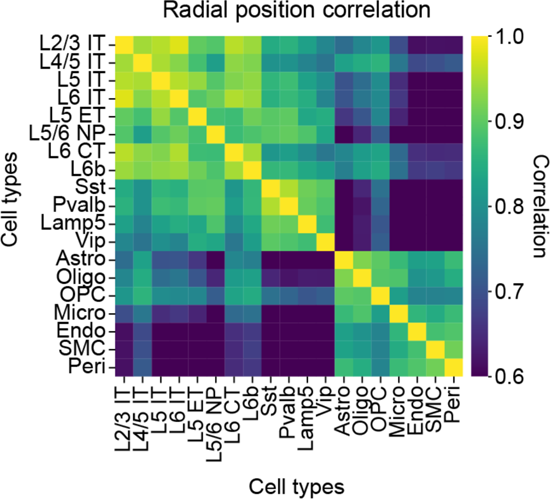
Correlation matrix for nuclear radial positions of genomic loci between different cell-type pairs. Normalized nuclear radial positions were calculated as in Fig. 4A. Each pixel in the heatmap represents the Pearson correlation coefficient of the normalized radial positions of all imaged chromosome loci between an indicated cell-type pair.

**Fig. S13.**
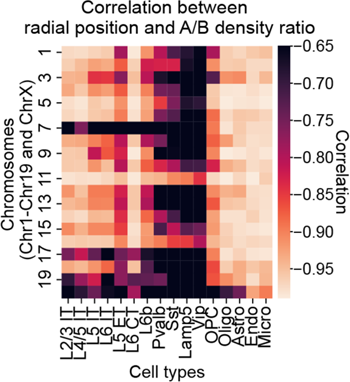
Correlation between cell-type medians of normalized radial positions and local A/B density ratios of individual genomic loci within each chromosome for each cell type.

**Fig. S14.**
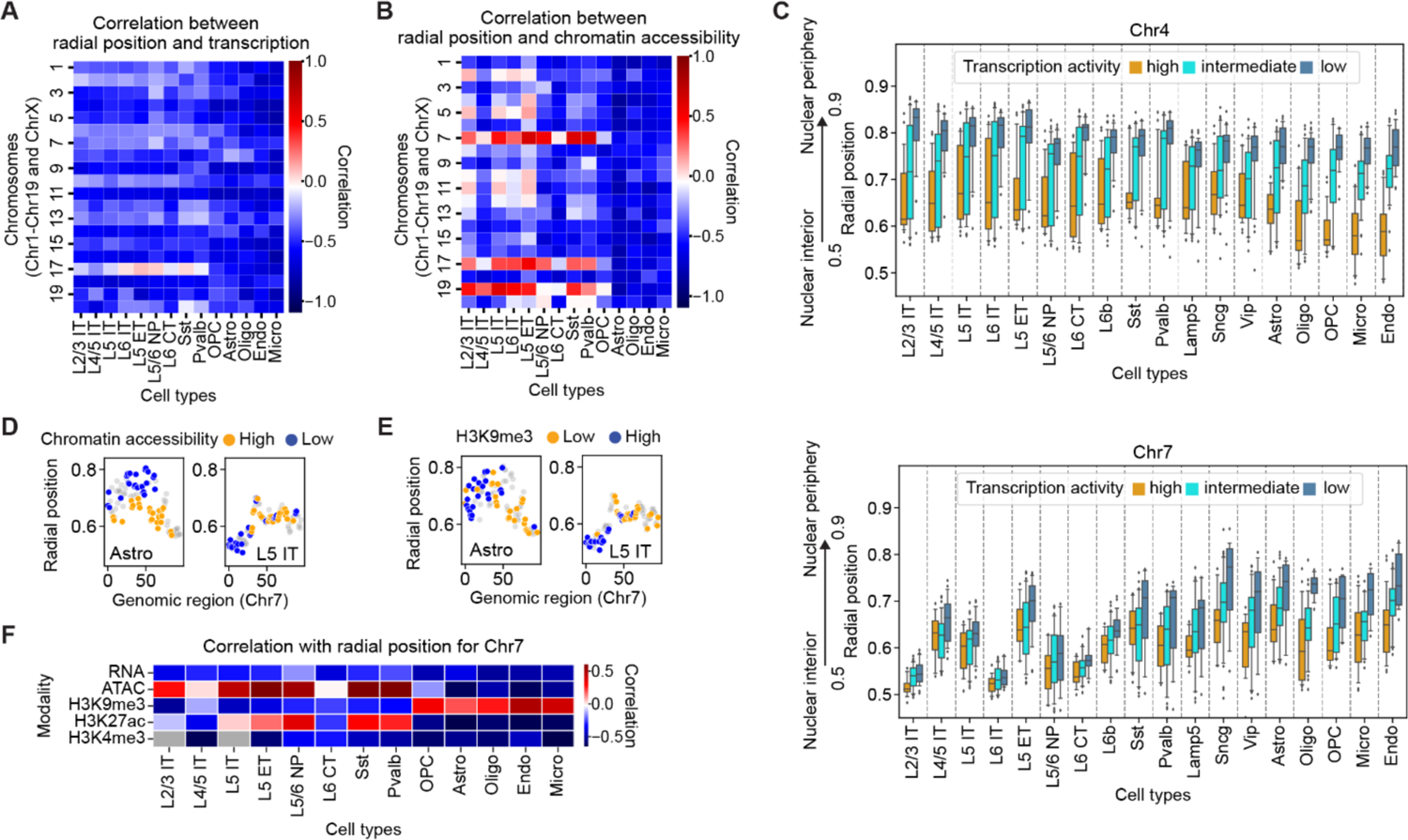
Relationship between nuclear radial positioning and various chromatin quantities across different cell types for different chromosomes. (**A**) Correlation between cell-type medians of normalized radial positions and transcriptional activity (calculated from snRNA-seq data (*52*)) of all loci from the indicated chromosome (y-axis) in the indicated cell type (x-axis). (**B**) Correlation between cell-type medians of normalized radial positions and chromatin accessibility (calculated from snATAC-seq data (*52*)) of all loci from the indicated chromosome (y-axis) in the indicated cell type (x-axis). (**C**) Boxplots for the distribution of normalized nuclear radial positions of loci in Chr4 (top) or Chr7 (bottom) loci for various cell types, grouped by transcriptional activity of the loci. Individual loci are grouped by their cell-type-median transcriptional activity as follows: high - top 25^th^ percentile; low - bottom 25^th^ percentile; intermediate - 25^th^-75^th^ percentile in transcriptional activity across all imaged loci (calculated from snRNA-seq data (*52*)). The center line, box, and whiskers in the boxplots represent the median, 25^th^-75^th^ percentile, and the 5^th^-95^th^ percentile, respectively. (**D**) Inverted radial organization of chromatin accessibility for Chr7 when comparing L5 IT excitatory neurons (right) to astrocytes (left). The dots are arranged by their genomic coordinates on the x-axis. The dots are colored by their chromatin accessibility (orange: top 25^th^ percentile in chromatin accessibility; blue: bottom 25^th^ percentile in chromatin accessibility; gray: all other loci), calculated from snATAC-seq data (*52*). (**E**) Similar to (D), but color coded by their H3K9me3 level. H3K9me3 level was calculated from Paired-Tag data (*63*). (**F**) Correlation between cell-type medians of normalized nuclear radial positions and cell-type medians of various other quantities (transcription level (RNA), chromatin accessibility (ATAC), H3K9me3 level, H3K27ac level, and H3K4me3 level) for genomic loci imaged on Chr7. Each pixel in the heatmap represents the Spearman correlation coefficient between the radial positioning and the indicated quantity (rows) of imaged loci on Chr7 in the indicated cell type (columns). Transcription and chromatin accessibility were calculated from snRNA-seq data and snATAC-seq data (*52*). H3K9me3, H3K27ac, and H3K4me3 levels were calculated from Paired-Tag data (*63*).

**Fig. S15.**
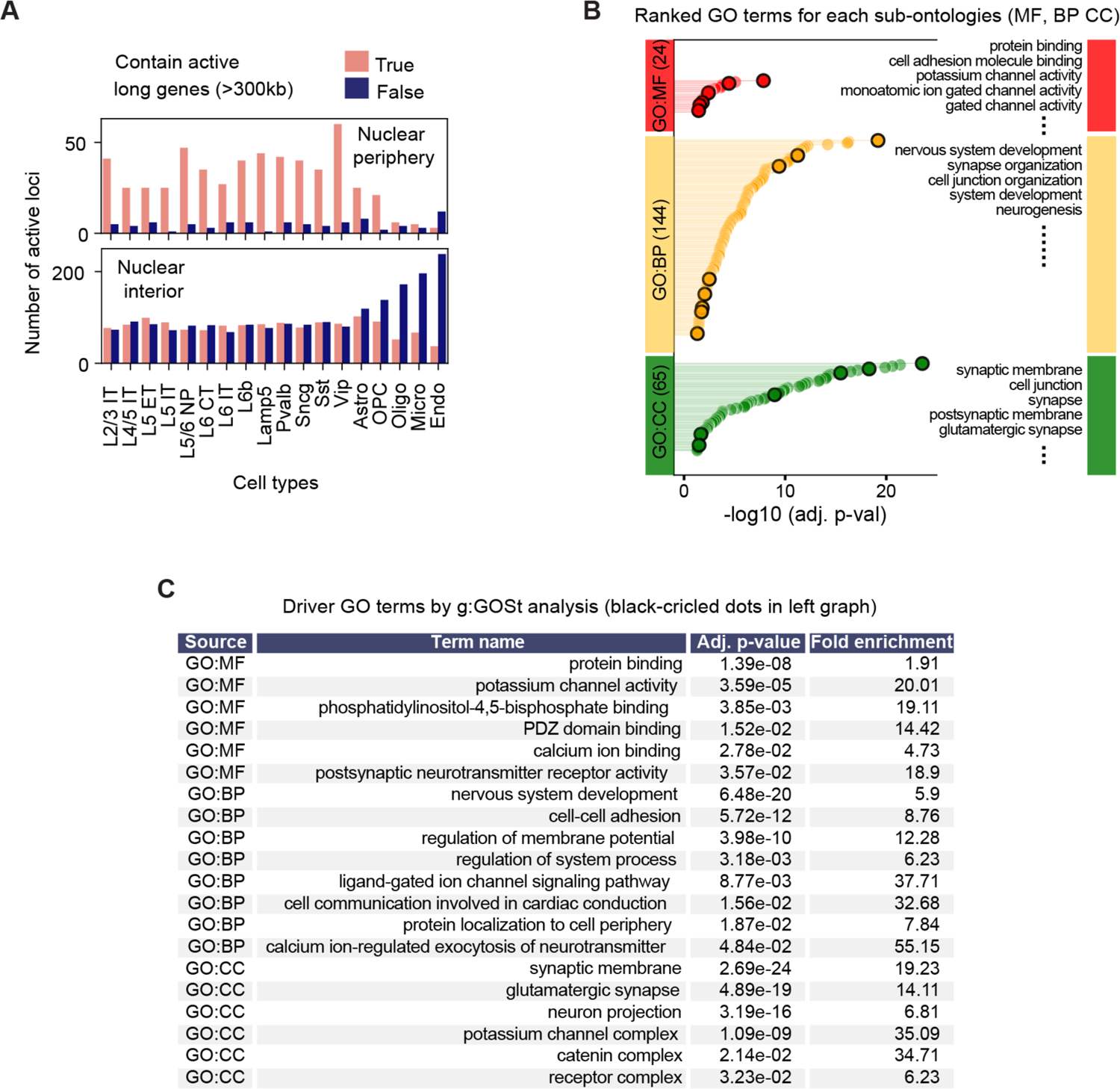
Active expression of long genes in the nuclear periphery in neurons. (**A**) Preferential activation of long genes near the nuclear periphery in neurons. Top: Number of transcriptionally active loci near the nuclear periphery, grouped by whether they harbor actively expressed long genes in each cell type (see Materials and Methods). Transcriptionally active loci are defined as loci in the top 25^th^ percentile of cell-type-median transcriptional level across all imaged loci, calculated from snRNA-seq data (*52*). Nuclear periphery loci are defined as loci in the top 20^th^ percentile of cell-type-median normalized nuclear radial position across all imaged loci. Long genes are defined as genes with >300 kb length. Bottom: similar to the top panel but for loci near the nuclear interior (defined as loci in the bottom 20^th^ percentile of cell-type-median normalized nuclear radial position across all imaged loci). (**B**) Gene Ontology (GO) analysis of actively expressed long genes in the nuclear periphery in neurons (see Materials and Methods). Ranked significant GO terms (p <0.05) identified from g:GOSt analysis (*102*) for each GO sub-ontologies are plotted. The total numbers of significant GO terms are indicated on the left bars, with the top 5 terms listed on the right. MF: molecular function; BP: biological process; CC: cellular component. Circled dots are driver GO terms. (**C**) Driver GO terms identified from g:GOSt analysis in (B) for each GO sub-ontologies.

**Fig. S16.**
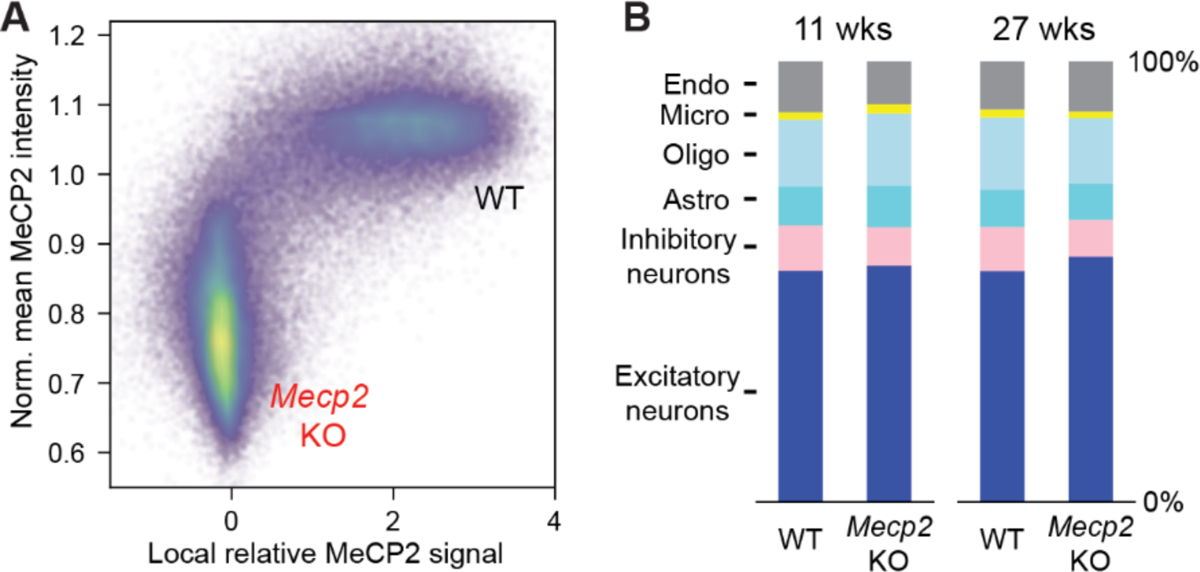
Cell-type compositions of *Mecp2* WT and KO cells. (**A**) Identification of *Mecp2* WT and KO cells based on protein signals of MeCP2 in *Mecp2* +/- female mice. The dots in the scatterplot were colored by the estimated kernel density of the dot distribution. Normalized MeCP2 intensity (y axis) is the mean intensity of MeCP2 antibody signals per cell normalized by the mean signals of all cells belonging to the same cell types in the same experimental replicate. Local relative MeCP2 signal (x axis) is the relative MeCP2 antibody signal per cell against its local background. (**B**) Composition of six major cell types in WT and *Mecp*2 KO cells from the MOp region of *Mecp2* +/- female mice, at the indicated ages of the animals.

**Fig. S17.**
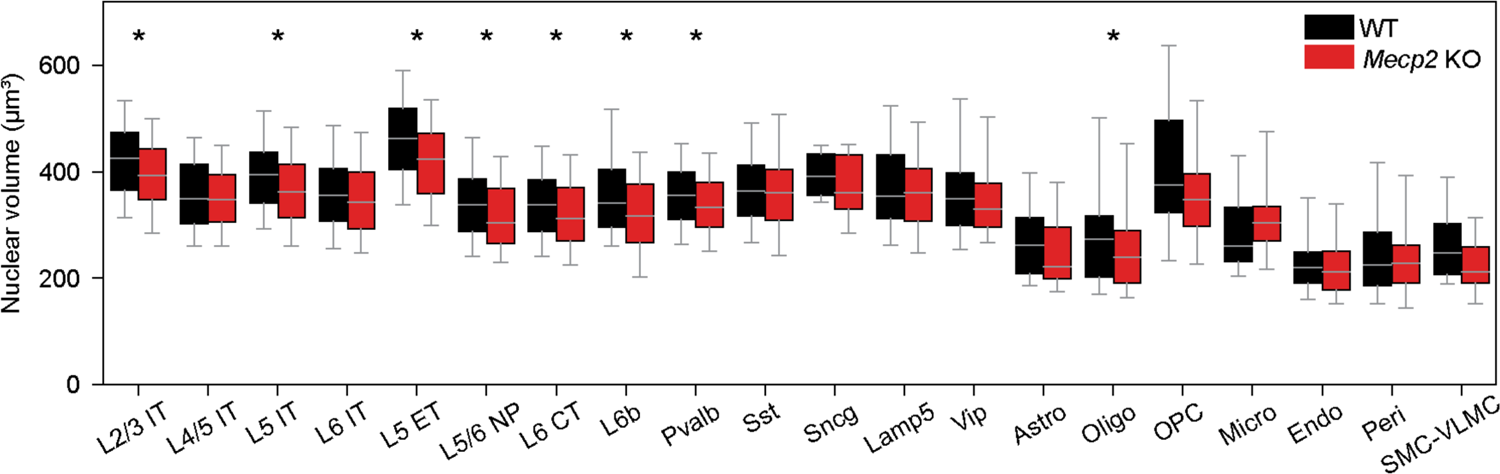
The effects of *Mecp2* deletion on the nucleus sizes in different cell types. Boxplots for the distribution of nuclear volume of individual cells for WT and *Mecp2* KO cells in each indicated cell type. The center line, box, and whiskers in the boxplots represent the median, 25^th^-75^th^ percentile, and the 5^th^-95^th^ percentile, respectively. Mann-Whitney U test, corrected by the Benjamini-Hochberg method, was used for statistical significance analysis. *: p<=0.05.

**Fig. S18.**
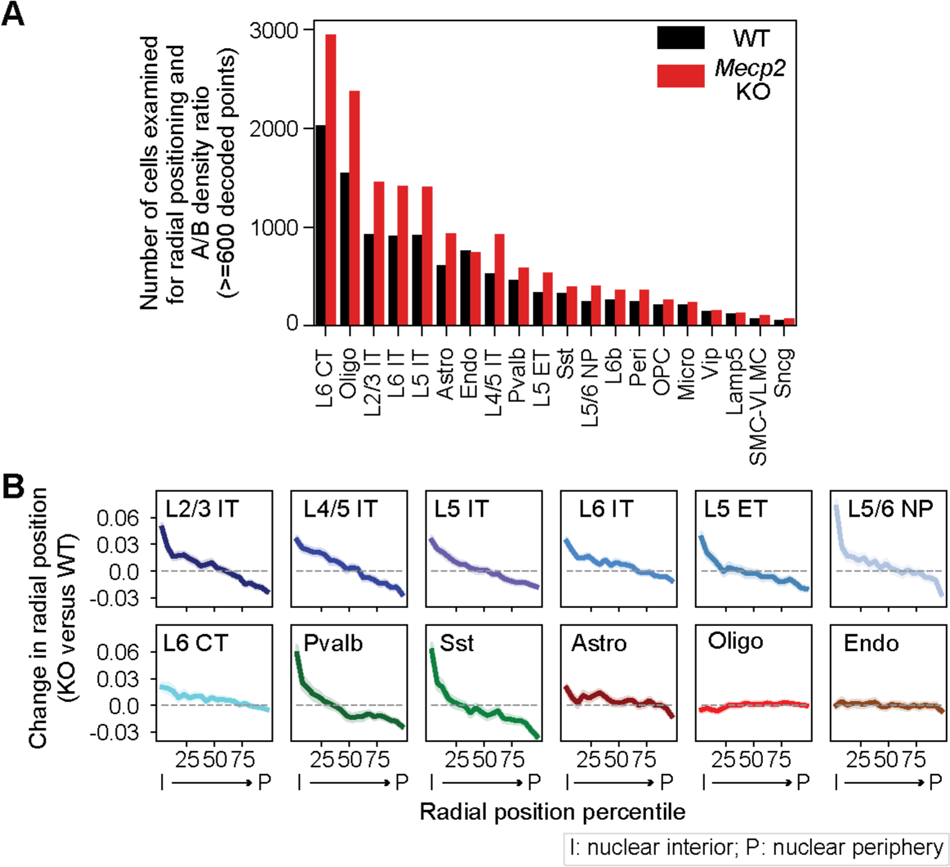
Cell-type-dependent effects of *Mecp2* deletion on radial positioning. (**A**) The number of quantified cells (with > 600 decoded chromatin loci per cell) for each cell type, for WT and *Mecp2* KO cells. (**B**) Changes in normalized nuclear radial positions upon *Mecp2* deletion as a function of the normalized radial position of the imaged genomic loci in various cell types. All imaged loci were grouped into 20 equal bins based on their cell-type median of normalized radial positions in the WT cells. For each locus, radial position change was calculated as the difference in the medians of normalized nuclear radial position in *Mecp2* KO over WT cells. The line and shaded area represent the mean and the 95% confidence interval, respectively.

**Fig. S19.**
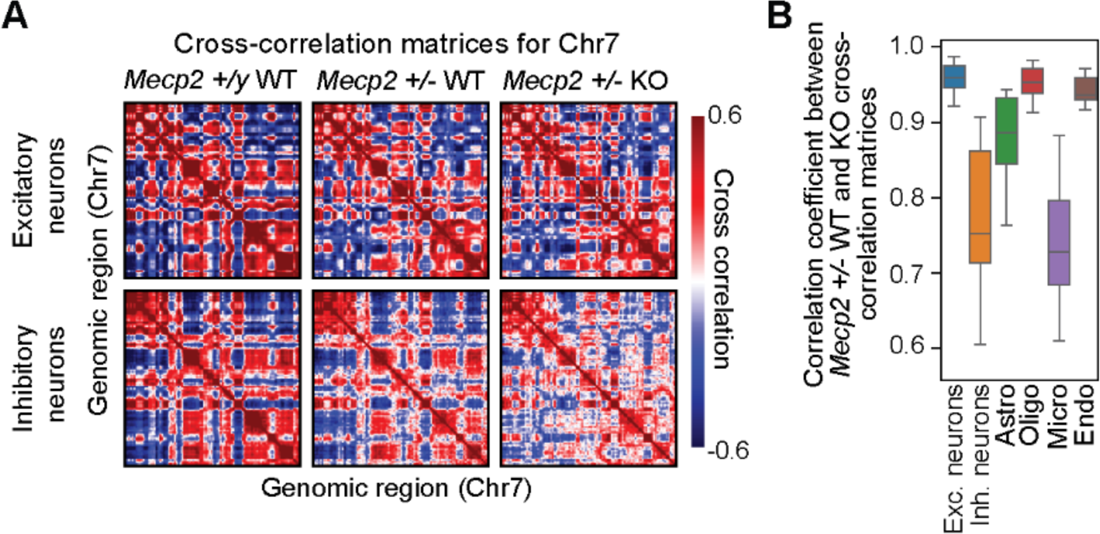
Cell-type-dependent effects of *Mecp2* deletion on A/B compartmentalization. (**A**) Cross-correlation matrices of Chr7 (4.5Mb – 143.8Mb) for excitatory neurons (top) and inhibitory neurons (bottom) for different genotypes. Left: WT cells from WT mice; Middle: WT cells from *Mecp2* +/- female mice; Right: *Mecp2* KO cells from *Mecp2* +/-female mice. Cross-correlation matrices were obtained as in Fig. 3A. No obvious changes in compartmental features were observed between WT and *Mecp2* KO excitatory neurons, and minor differences were observed for inhibitory neurons. (**B**). Boxplots for the distribution of the Pearson correlation coefficients of the cross-correlation matrices between the WT and *Mecp2* KO cells from *Mecp2* +/- mice. For each chromosome (e.g., Chr1), the Pearson correlation coefficient was calculated between the vectorized upper triangular elements of cross-correlation matrix from WT cells and those from *Mecp2* KO cells. The boxplot shows the distribution across the 19 chromosomes (Chr1-Chr19). The center line, box, and whiskers in the boxplots represent the median, 25^th^-75^th^ percentile, and 5^th^-95^th^ percentile, respectively.

**Fig. S20.**
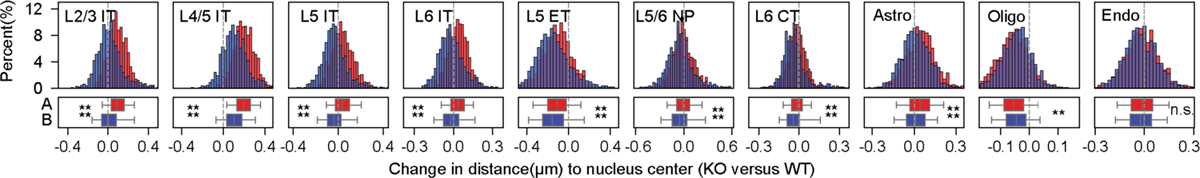
Changes in nuclear radial positioning of A- and B-compartment loci upon *Mecp2* deletion. Histograms of nuclear radial position change upon *Mecp2* deletion for compartment-A (red) or compartment-B (blue) loci in various cell types. The corresponding boxplots of the distributions for A and B loci are shown at the bottom. The center line, box, and whiskers of the boxplots represent the median, 25^th^-75^th^ percentile, and 5^th^-95^th^ percentile, respectively. Student’s t-tests with Bonferroni corrections were used for statistical significance characterization. *: p<=0.05; ***: p<=0.001; ****: p<=0.0001; n.s.: p>0.05.

## Supplementary table captions

**Table S1. Encoding-probe libraries for integrated RNA- and DNA-MERFISH.** A list of all the template oligonucleotides used to make the encoding probes for RNA- and DNA-MERFISH imaging. The file contains five spreadsheets – “RNA MERFISH probes”, “Genome chromatin probes”, “TSS chromatin probes”, “Super-enhancer chromatin probes”, and “TSS chromatin pbs (probes) by MERFISH”. (1) “RNA MERFISH probes” contains all template sequences for making encoding probes for RNA-MERFISH. (2) “Genome chromatin probes” contains all template sequences for making encoding probes for DNA-MERFISH imaging of the 988 genomic loci evenly distributed across the genome. (3) “TSS chromatin probes” contains all template sequences for making encoding probes for sequential DNA-FISH imaging of the TSSs of 28 cell-type marker genes. (4) “Super-enhancer chromatin probes” contains all template sequences for making encoding probes for DNA-MERFISH imaging of the 965 super-enhancer loci. (5) “TSS chromatin pbs (probes) by MERFISH” contains all template sequences for making encoding probes for DNA-MERFSH imaging of TSSs of 28 cell-type marker genes.

**Table S2. Adaptor probes for integrated RNA- and DNA-MERFISH.** The file contains four spreadsheets – “RNA MERFISH”, “Genome chromatin”, “TSS chromatin” and “Super-enhancer chromatin”. Each spreadsheet corresponds to a list of the adaptor probes that were used to hybridize the readout sequences on the encoding probes, from each library described in Table S1. The names of these adaptor probes are formatted in the following way unless otherwise described. Using the first adaptor probe name from the “RNA MERFISH” sheet as an example (“Bit-1-RS0015_2xStv_82rc”), “Bit-1” corresponds to the first bit in the 22-bit RNA-MERFISH imaging. “RS0015” corresponds to the name of a unique readout sequence on the encoding probes (see “name” column in Table S1), and “2xStv_79rc” corresponds to two binding sequences of the dye-conjugated common readout probe named “Stv_79” (see “name” column in Table S3). The last adaptor probe name in the “RNA MERFISH” sheet (“Bit-polyT-RS2179_2xStv_82rc”) corresponds to the adaptor probe that target polyA-anchor probe (/5Acryd/TTGAGTGGATGGAGTGTAATT+TT+TT+TT+TT+TT+TT+TT+TT+TT+T).

**Table S3. Readout probes and PCR primers for integrated RNA- and DNA-MERFISH.** The file contains two spreadsheets – “Common readouts” and “Primers”. (1). “Common readouts” contains the names and sequences of three dye-conjugated common readout probes, which were used to hybridize the two binding sequences on each adaptor probe, as described in Table S2. (2) “Primers” contains all forward and reverse primer pairs used for amplification of the template oligonucleotides, for each library described in Table S1. For “Genome chromatin” library, the template oligonucleotides were divided into three sets (“sets 1 and 2”, “set 3”, and “set 4) by a ratio of ∼ 2:1:1 for the total number of probes from each set. References

## Notes

https://data.4dnucleome.org/experiment-set-replicates/4DNESPE924IP/

https://data.4dnucleome.org/experiment-set-replicates/4DNESMTNNB3N/

